# Inhibition of the ATM kinase rescues planarian regeneration after lethal radiation

**DOI:** 10.1101/2022.09.28.509579

**Authors:** Divya A. Shiroor, Kuang-Tse Wang, Bhargav D. Sanketi, Justin K. Tapper, Carolyn E. Adler

**Author notes:** These authors contributed equally (Divya A. Shiroor, Kuang-Tse Wang).

## Abstract

As stem cells divide, they acquire mutations that can be passed on to daughter cells. To limit the possibility of propagating mutations, cells activate the DNA damage response (DDR) network, which dictates whether cells repair DNA or undergo apoptosis. At the helm of the DDR are three PI3-like kinases including <u>A</u>taxia <u>T</u>elangiectasia <u>M</u>utated (ATM). We report here that knockdown of ATM in planarian flatworms enables stem cells, which normally undergo apoptosis after radiation exposure, to survive lethal doses of radiation. In this context, stem cells circumvent apoptosis, replicate their DNA, and recover function using homologous recombination-mediated DNA repair. Despite radiation exposure, *atm* knockdown animals survive long-term and regenerate new tissues. These effects occur independently of ATM’s canonical downstream effector p53. Together, our results demonstrate that ATM’s primary function is to drive apoptosis, and suggest that inhibition of ATM could therefore potentially favor cell survival after radiation without adverse effects.

## Introduction

Stem cells fuel the production of new cells during homeostasis and tissue regeneration (Tanaka and Reddien, 2011), despite continual exposure to environmental and organismal insults (e.g. aging) that increase the probability of acquiring mutations with each round of cell division. Mutations can alter the output of stem cells, often adversely, by driving senescence, premature differentiation, or promoting tumorigenesis (Blanpain *et al*., 2011; Mandal, Blanpain and Rossi, 2011). Whether stem cells undergo cell death or repair their DNA depends on their cell cycle status, the intrinsic turnover rate of tissues, and the probability of entering quiescence (Borges, Linden and Wang, 2008). To combat the accumulation of DNA damage and prevent tumorigenesis, adult stem cells engage different arms of the DNA damage response (DDR) pathway depending on the tissue where they reside (Mandal, Blanpain and Rossi, 2011; Soteriou and Fuchs, 2018). For example, mouse hair follicle stem cells rely on increased expression of DNA repair proteins and anti-apoptotic factors to repair and evade apoptosis (Sotiropoulou *et al*., 2010), while stem cells in the small intestine exhibit increased radiation sensitivity due to reduced expression of anti-apoptotic proteins (Merritt *et al*., 1995). The difference among stem cell types raises the question of how stem cells cope with DNA damage and the underlying mechanisms that determine their fates.

In many species, the choice to repair DNA or initiate apoptosis is coordinated by three apical kinases <u>A</u>taxia <u>T</u>elangiectasia <u>M</u>utated (ATM), <u>AT</u>M and <u>R</u>ad3-related (ATR), and the <u>DNA</u>-dependent <u>p</u>rotein <u>k</u>inase (DNA-pk), which are members of the phosphoinositide 3-kinase (PI3K)-related kinase family (Fig. EV1A) (Blackford and Jackson, 2017). ATM and ATR serve as important checkpoint regulators that stall the cell cycle upon DNA damage (Blackford and Jackson, 2017). Cells and patients lacking ATM are often hypersensitive to radiation (Xu and Baltimore, 1996; Shiloh and Ziv, 2013). However, some cell types, including thymocytes and postmitotic neurons, are radiation-resistant, suggesting that dependence on ATM can vary (Xu and Baltimore, 1996; Herzog *et al*., 1998). Importantly, recent data comparing the ability of hematopoietic stem cells to reconstitute the blood shows that cells heterozygous for ATM can restore blood faster than homozygotes or wild-type controls (Ito *et al*., 2004; Fortin *et al*., 2021). These results suggest that tuning apical kinase activity in different cell types could be a strategy used to manipulate stem cell outcomes.

Planarian flatworms have an abundant population of adult stem cells that actively divide to maintain tissue homeostasis and fuel regeneration (Rink, 2013). Exposing planarians to ionizing radiation introduces double-strand breaks (DSBs) that induce apoptosis (Pellettieri *et al*., 2010). High-dose radiation (6,000 rads, or 60 Gray) depletes all the stem cells within 2 days (Reddien *et al*., 2005; Eisenhoffer, Kang and Sánchez Alvarado, 2008; Wagner, Ho and Reddien, 2012), while lower doses of radiation (2,000 rads, or 20 Gray) allow a few stem cells to persist. Although these cells slowly resume dividing, the animals still succumb (Wagner, Wang and Reddien, 2011; Lei *et al*., 2016), suggesting that irreversible damage has occurred. This eventual death suggests that despite survival at the cellular level, stem cells are unable to counterbalance DNA damage at sufficient rates to fuel long-term animal viability.

Even at radiation doses of 2,000 rads, the fact that some stem cells survive while others perish within the same animal suggests that factors such as cell cycle state play an important role in driving radiation resistance. Despite conservation of key cell cycle regulators and components of the DDR pathway (Grohme *et al*., 2018; Sahu *et al*., 2021), molecular mechanisms that govern planarian stem cell responses to radiation are as yet unclear (Barghouth *et al*., 2019). Here, we capitalize on the radiation sensitivity of planarian stem cells to understand how these cells decide whether to repair their DNA or undergo apoptosis. We find that ATM drives these cells to undergo apoptosis after exposure. Depletion of ATM prevents stem cell death after radiation. The stem cells that persist escape into the S and G2 stage of the cell cycle. These cells subsequently rely on homologous-recombination mediated repair to overcome sustained DNA damage and can successfully fuel animal regeneration and repair. Together, our data uncovers a specific role for ATM in driving apoptosis after radiation, and highlights how the function of this conserved protein might have evolved in planarian stem cells. This study therefore reveals new mechanisms for promoting stem cell rejuvenation and preventing tumorigenesis in less resilient animals.

## Results

To determine whether apical kinases regulate stem cell responses to radiation in planarians, we used RNA interference (RNAi) to knock down orthologs of *atm, atr* and *dna-pk* and the MRN complex components *<u>m</u>re11, <u>r</u>ad50*, and *<u>n</u>bs1*, which are enriched in stem cells (Fig. EV1B). Two days after radiation exposure at a dosage of 2,000 rads, we examined stem cell distribution and abundance with *in situ* hybridization and qRT-PCR for the canonical stem cell marker *piwi-1* (*Reddien et al., 2005*). Knockdown of *atm*, *atr*, and MRN complex components caused *piwi-1^+^* cells to persist after radiation (Fig. 1, A and B), while virtually no stem cells remained in control animals. Among these genes, *atm*(RNAi) preserved the most stem cells, as shown by *piwi-1* expression levels in qRT-PCR (Fig. 1B) regardless of the RNAi trigger (Fig. EV2A-C). Based on these results, we conclude that the apical kinases *atm* and *atr*, but not *dna-pk*, indeed regulate radiation-induced stem cell loss.

**Figure 1:**
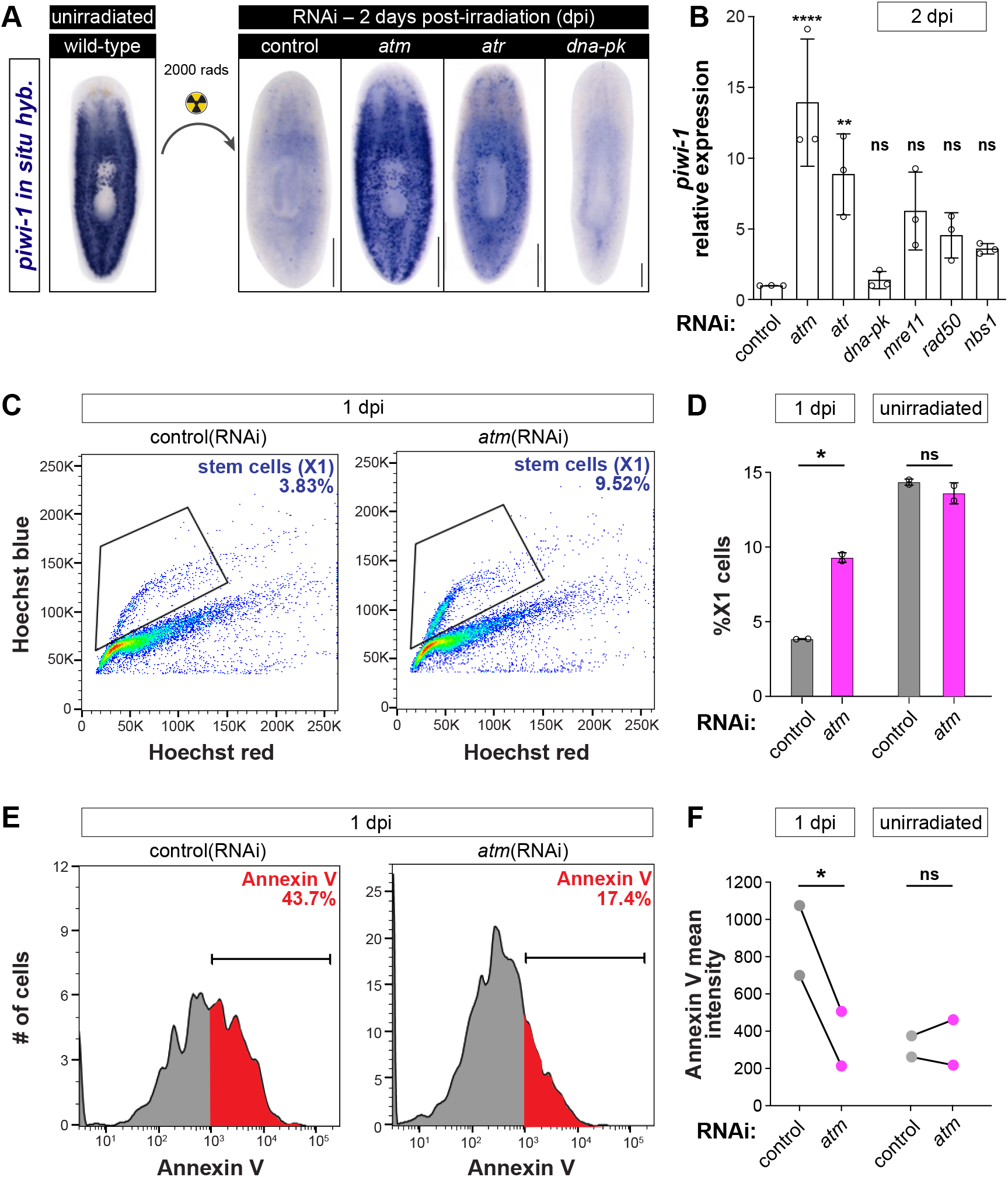
ATM drives stem cell apoptosis after radiation. **A**, Experimental design and *piwi-1* whole mount *in situ* hybridization (WISH) of RNAi animals, 2 days post-irradiation (dpi). **B**, qRT-PCR of *piwi-1* relative to GAPDH in RNAi animals as indicated, 2dpi. Statistics represent ordinary one-way ANOVA compared to controls; ****P<0.0001, **P<0.005. **C**, FACS plots of Hoechst-stained X1 cells from control and *atm*(RNAi) animals, 1 dpi. Black polygon=stem cells (X1), percentage indicated in blue. **D**, Percentage of X1 cells from control and *atm*(RNAi) animals, 1dpi or unirradiated (Fig. S2E). **E**, Histograms of X1s from (C) stained with Annexin V (percentage in red). Brackets and red shaded area=Annexin V mean fluorescent intensity staining >10^3^. **F**, Mean fluorescent intensity of Annexin V calculated from bracketed region in (E) or (Fig. S2F); lines connect results from individual experiments. For all panels, radiation dose=2000 rads. Data are averages ± SD; circles represent biological replicates; ns=not significant. In (D) and (F), statistics represent paired t-test, *P<0.05. Scale bars=250μm.

To evaluate if stem cells in radiated *atm*(RNAi) animals might be evading cell death, we purified stem cells (X1s) using fluorescence activated cell sorting (FACS) (Reddien *et al*., 2005; Hayashi *et al*., 2006) and stained them with the cell death marker Annexin V (Peiris, García-Ojeda and Oviedo, 2016; Shiroor, Bohr and Adler, 2020). One day after radiation, *atm*(RNAi) animals had significantly more X1s as compared to controls (9.52% vs 3.83%) (Fig. 1, C and D), which was accompanied by a reduction in Annexin V staining (Fig. 1, E and F). By contrast, in unirradiated RNAi animals, there was no difference in either stem cell abundance or Annexin-V-positive X1s (Fig. 1, D, F; Fig. EV2, D-G), suggesting that ATM drives stem cell apoptosis primarily after radiation.

To examine the transcriptional response that enables stem cell survival after radiation, we purified stem cells from *atm*(RNAi) animals and subjected them to RNA-sequencing (Fig. EV3A). Following radiation, 431 transcripts were upregulated in *atm*(RNAi) stem cells, while in unirradiated animals, the only differentially-expressed transcript was *atm* (Fig. EV3, B and C, Supplementary Tables 1-3). Classification of upregulated transcripts using Gene Ontology (GO) analysis revealed that *atm*(RNAi) stem cells were enriched for terms related to cell cycle regulation, DNA replication, and gene expression (Fig. EV3D). To determine if these genes might function similarly to *atm*, we knocked down transcripts either significantly upregulated in *atm*(RNAi) stem cells or those associated with DDR terms (Fig. EV3E). Among these, only the conserved ATM effector, *checkpoint kinase 2* (*chk2*), phenocopied *atm*(RNAi) (Fig. EV3, F and G) (Thompson, 2012). Notably, knockdown of the classic ATM effector *p53* did not influence stem cell persistence following radiation. This finding is consistent with the function of planarian p53 having diverged away from monitoring genotoxic stress (Pearson and Sánchez Alvarado, 2010; Cheng *et al*., 2018; Wendt *et al*., 2022). Together, these data indicate that the MRN-ATM-Chk2 axis controls stem cell abundance after radiation, independently of canonical p53 activity.

Based on ATM’s known roles in activating cell cycle checkpoints (Waterman, Haber and Smolka, 2020), we analyzed cell cycle progression in *atm*(RNAi) animals following radiation. First, to label cells undergoing DNA replication, we administered the thymidine analog F-ara-EdU (EdU) (Neef and Luedtke, 2011; Bohr, Shiroor and Adler, 2021) for 24 hours on the first or second day after radiation exposure (Fig. 2A). In stark contrast to controls, most *atm*(RNAi) animals had abundant EdU signal one day after radiation (Fig. 2, A and B), indicating continued DNA replication and a failure to activate the intra-S checkpoint. This transient progression into S phase after *atm* knockdown phenocopies the radioresistant DNA synthesis observed in human cells with mutations in ATM (Painter, 1981). Second, to determine whether cells have also progressed into G2, we quantified DNA content with DAPI staining, and observed an increase in G2/M cells in *atm*(RNAi) animals (Fig. 2C). However, staining for the mitotic marker phosphohistone-H3 (pH3) 2 days after radiation showed no cells in M phase (Fig. 2, D and E), suggesting that stem cells either fail to *atm*(RNAi) animals accumulate in S and G2. Therefore, upon DNA damage, ATM is necessary for G1/S and intra-S checkpoints (Xu *et al*., 2002), but not the metaphase checkpoint.

**Figure 2:**
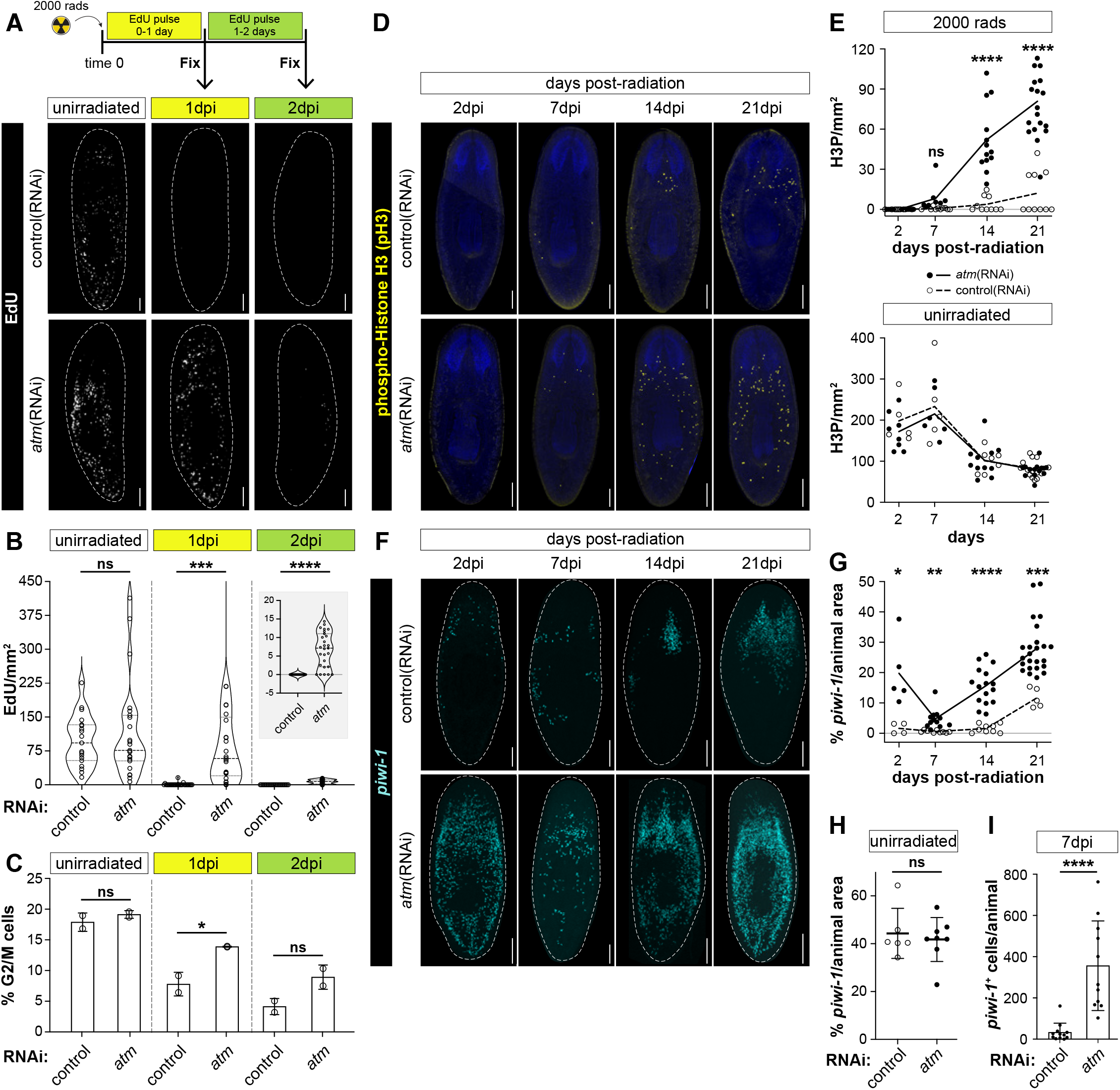
*atm*(RNAi) enables DNA replication and stem cell repopulation after radiation. **A**, Top: Strategy for EdU pulse and animal fixation. Bottom: Confocal images of EdU-stained *atm* and control RNAi animals. **B**, Quantification by area of EdU^+^ cells from A. Inset, 2dpi with altered scale. **C**, Graph showing percentage of G2/M cells in *atm* and control RNAi animals on indicated dpi. **D**, Confocal images of phosphohistone H3 (pH3)-stained RNAi animals after radiation exposure. **E**, Quantification by area of pH3 cells in radiated (top) and unirradiated (bottom) animals after *atm* (solid line) and control (dashed line) RNAi. **F**, Confocal images of *piwi-1* FISH in RNAi animals after radiation exposure. **G and H**, Percentage of animal area occupied by *piwi-1* FISH, divided by total animal area, in radiated (**G**) or unirradiated (**H**) RNAi animals. **I**, Total number of *piwi-1^+^* cells in RNAi animals, 7 dpi. Data are averages ± SD; circles represent biological replicates; statistics represent unpaired t-test between control and *atm*(RNAi), ****P<0.0001, ***P<0.0004, **P<0.0002, *P<0.05, ns=not significant. Scale bars=250μm.

To determine whether the stem cells that persist in *atm*(RNAi) animals are capable of resuming mitosis and repopulating the animal, we monitored stem cells for three weeks after radiation. We detected the reappearance of phosphohistone-H3-positive cells in *atm*(RNAi) animals beginning one week after radiation, with their numbers increasing significantly thereafter (Fig. 2, D and E). We confirmed stem cell expansion in *atm*(RNAi) animals by quantifying *piwi-1^+^* cells (Fig. 2, F-I), with significant repopulation beginning 7 days after radiation. The initial decline of stem cells in *atm*(RNAi) animals between 2 and 7 days after radiation (Fig. 2G) implies that either ATM is not required for apoptosis during this time, or that there may be residual ATM activity eliminating stem cells with severe DNA damage. Together, these findings indicate that *atm*(RNAi) stem cells can ultimately pass through cell cycle checkpoints, implying that DNA damage has been repaired.

Planarians require stem cell division and differentiation for tissue renewal and regeneration. Radiation normally results in animal death in about 6 weeks (Wagner, Wang and Reddien, 2011; Li, Taylor and van Wolfswinkel, 2021). To test whether the stem cells that repopulate *atm*(RNAi) animals are functional, we monitored animal survival after exposing them to a dosage of 2,000 rads (Fig. 3A). In control animals, only a few stem cells persist after this dose, and these cells are insufficient to support regeneration (Wagner, Wang and Reddien, 2011; Shiroor, Bohr and Adler, 2020). Unexpectedly, over 90% of radiated *atm*(RNAi) animals survived long-term without developing gross abnormalities (Fig. 3B) (Pearson and Sánchez Alvarado, 2010). To directly examine regeneration, we decapitated animals immediately after radiation. Consistent with the survival data, 80% of *atm*(RNAi) animals fully regenerated new heads, including photoreceptors and the central nervous system (Fig. 3, C and D). The recovery of regenerative capacity indicates that *atm*(RNAi) animals have completely restored all stem cell function, including normal differentiation and patterning.

**Figure 3:**
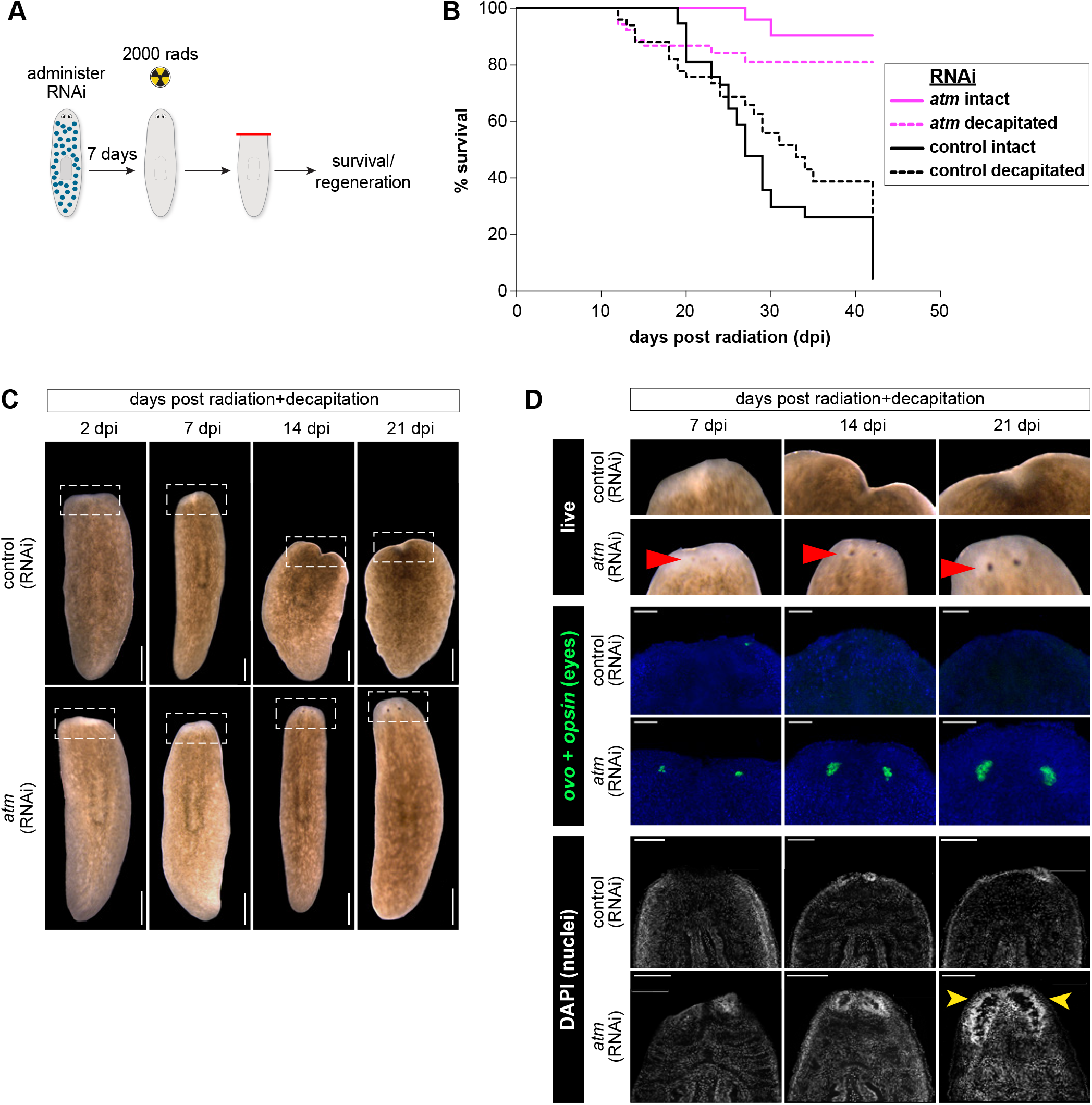
Surviving stem cells fuel regeneration and long-term animal survival. **A**, Schematic of experimental design. **B**, Kaplan-Meier survival curves of intact (solid line) and decapitated (dashed line) RNAi animals after radiation. n = 45 animals, including two biological replicates. **C**, Live images of regenerating control and *atm* RNAi animals after radiation exposure. Scale bars=250μm. **D**, Images of heads in RNAi animals after radiation. Top, boxed region from (C); Middle, *ovo*+*opsin* FISH for eyes (green) counterstained with DAPI (blue); bottom, DAPI staining of the brain. Red arrowheads, eyes; yellow arrowheads, brain. Scale bars=100μm.

Cells repair DSBs through activation of homologous recombination (HR) and non-homologous end joining (NHEJ) (Ciccia and Elledge, 2010; Thompson, 2012; Santivasi and Xia, 2014). To test the dependence of surviving stem cells on these DNA repair pathways, we simultaneously knocked down components of HR or NHEJ in *atm*(RNAi) animals. We used the recombinase *rad51* or its interacting partner *brca2* as a proxy for HR (Peiris *et al*., 2016; Sahu *et al*., 2021), and *dna-pk* as a proxy for NHEJ. Knockdown of either *rad51* or *brca2* accelerated animal death after radiation (Fig. 4A). This accelerated death was unaffected by simultaneous knockdown of *atm*, suggesting that although *atm* preserves stem cells initially after radiation by preventing apoptosis, long-term survival requires homologous recombination-mediated DNA repair. By contrast, knockdown of the apical kinase *dna-pk*, which mediates NHEJ, did not substantially alter the timing of animal death after radiation as compared to controls (Fig. 4B). However, animals in which *dna-pk* was knocked down along with *atm* survived long-term. Together, these results demonstrate that even when apoptosis is inhibited by *atm*(RNAi), NHEJ-mediated DNA repair is insufficient to support stem cell recovery from the deleterious effects of radiation. Rather, in stem cells that escape apoptosis, homologous recombination-mediated repair acts to restore stem cell function.

**Figure 4:**
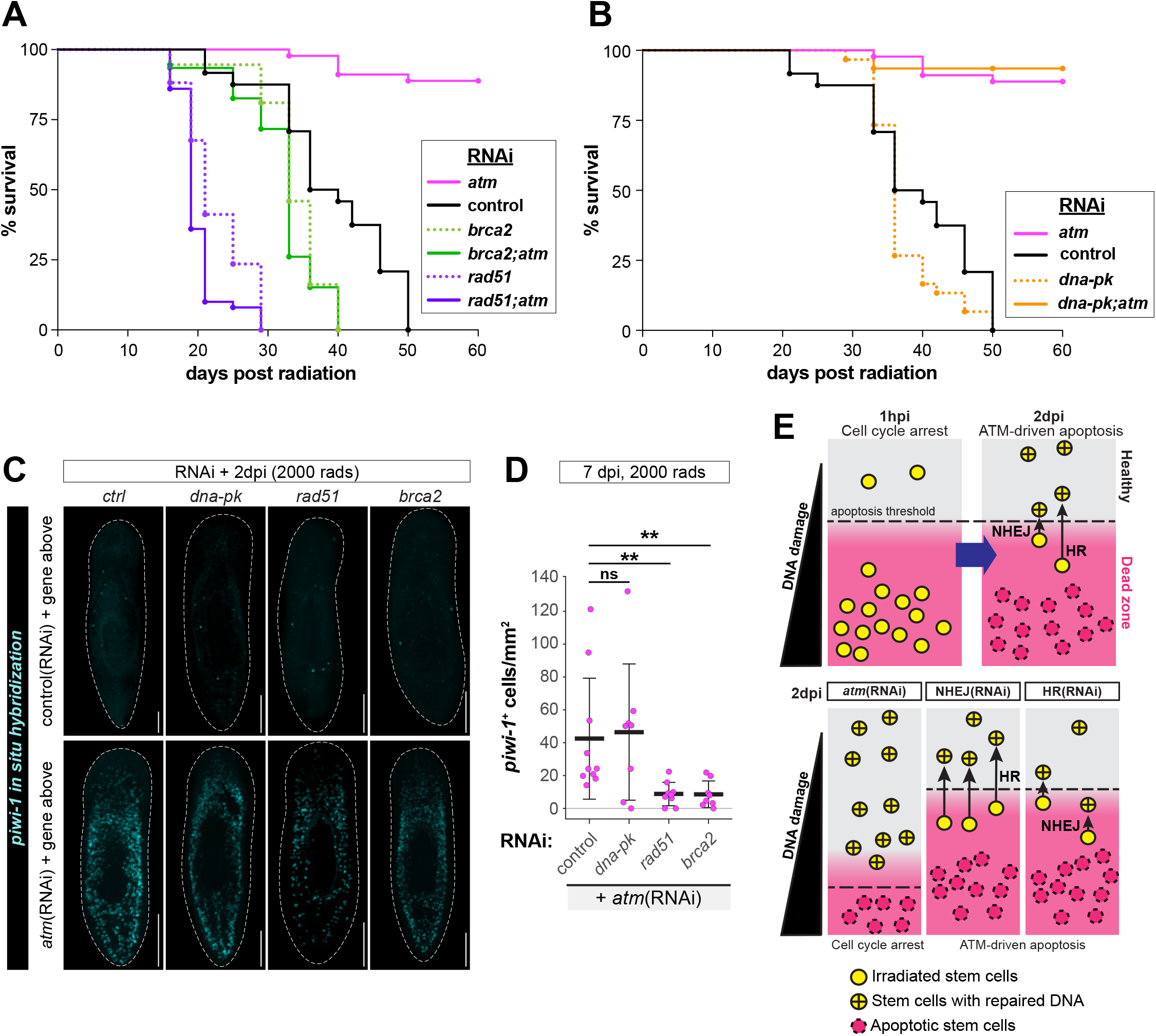
Animal survival after radiation relies on homologous recombination. **A** and **B**, Kaplan-Meier survival curves of intact RNAi animals after radiation, n=25 animals. Both experiments were performed simultaneously but split into two graphs for clarity; control and *atm*(RNAi) values are replicated in both panels. **C**, Confocal images of *piwi-1* FISH in RNAi animals after exposure to 2000 rads, 2dpi. Scale bars=250μm. **D**, Quantification by area of *piwi-1^+^* cells in RNAi animals after 2000 rads exposure, 7dpi. Data are averages ± SD; circles represent biological replicates; ordinary one-way ANOVA compared to control **P<0.005, ns=not significant. **E**, Model depicting mechanism of stem cell survival after radiation. ATM drives stem cell apoptosis by setting a threshold for cell death based on levels of DNA damage at 2dpi. “Healthy zone” refers to stem cell persisting at 2dpi; “Dead Zone” refers to stem cells with severe DNA damage that die between 2dpi and 7dpi. HR-mediated DNA repair, moreso than NHEJ-mediated repair, enables stem cell survival after radiation.

To confirm that long-term survival after perturbation of NHEJ and HR components was due to aberrant persistence of stem cells, we analyzed their distribution. Two days after radiation, stem cell abundance was not significantly altered by knockdown of HR or NHEJ components (Fig. 4C, Fig. EV4A), although single knockdown of NHEJ-specific components *artemis, lig4, ku70*, and *ku80* caused modest increases in stem cell survival and proliferation (Fig. EV4, B-E). Seven days after radiation exposure, when proliferation normally resumes, we found that stem cells were preserved following knockdown of *dna-pk*, but not *brca2* or *rad51* (Fig. 4D), further indicating a reliance on HR at this stage. The failure of stem cell persistence in *brca2* or *rad51* knockdown suggests that stem cells in *atm*(RNAi) animals employ HR, but not NHEJ, to resume proliferation.

## Discussion

In this study, we perturbed expression of apical DDR kinases and assessed their function in regulating planarian stem cell behavior after ionizing radiation. We find that ATM is a key driver of stem cell apoptosis, but strictly in response to massive DNA damage. Stem cells in *atm* knockdown animals continue replicating their DNA after radiation, escaping radiation-induced checkpoints that would otherwise arrest the cell cycle immediately. The cells that manage to replicate their DNA generate a template for the homologous recombination machinery to carry out repair. Therefore, we propose a model in which ATM sets a threshold for activation of stem cell apoptosis based on the extent of DNA damage (Fig. 4D).

Our results demonstrate that planarian stem cells depend on homologous recombination for stem cell recovery, the primary mode of high-fidelity repair (Krenning, van den Berg and Medema, 2019). In wildtype animals, while both HR- and NHEJ-mediated repair strategies are presumably activated by radiation, cells with DNA damage exceeding the threshold set by ATM are driven to undergo apoptosis. Conversely, despite bearing a DNA damage load equivalent to wildtype animals, stem cells in ATM knockdown animals do not undergo apoptosis, providing an opportunity for DNA repair. In *atm*;NHEJ double knockdown animals (Fig. 4), high-fidelity HR is the sole DNA repair mechanism, driving robust stem cell recovery. By contrast, in *atm*;HR double knockdown animals, stem cells can only deploy error-prone NHEJ pathways, resulting in cell death due to activation of cell cycle checkpoints as they undergo mitosis. The probability of undergoing cell death may also be regulated by cell cycle status at the time of radiation, leading to the stochastic persistence of small numbers of cells after low radiation doses (Krenning, van den Berg and Medema, 2019; Raz, Wurtzel and Reddien, 2021). Together, our model proposes that DNA damage is not well tolerated by planarian stem cells, and that they are more likely to undergo apoptosis than propagate mutations. This finding may explain in part the anecdotal observation that planarians do not form tumor-like growths (Pearson and Sánchez Alvarado, 2008).

Despite conservation of core apoptotic pathways, identifying the mechanisms responsible for stem cell death in planarians have remained elusive (Pellettieri and Sánchez Alvarado, 2007; Bender *et al*., 2012). Elimination of damaged cells in other animals normally depends on activation of ATM, along with its canonical downstream effectors Chk2 and p53 (Shiloh and Ziv, 2013). However, in planarians, ATM seems to operate independently of p53 in driving cell death (Fig. EV3F) (Wendt *et al*., 2022). This difference may be due to a functional loss of planarian p53. Despite conservation of the protein, planarian p53 acts primarily to regulate epithelial cell production rather than to respond to genotoxic stress (Pearson and Sánchez Alvarado, 2010; Cheng *et al*., 2018; Wendt *et al*., 2022). On the other hand, some aspects of the ATM pathway remain intact, because knockdown of its canonical effector Chk2 also causes stem cell persistence after radiation (Fig. EV3G). Identifying downstream targets mediating these p53-independent outcomes in planarians may reveal how these pathways have evolved in different animal contexts.

Unlike in other eukaryotes where ATM regulates cell cycle checkpoints in response to replication stress (Waterman, Haber and Smolka, 2020), planarian ATM seems to act strictly after radiation, without any obvious role in unirradiated contexts. This difference in function could be due to altered protein interactions, because planarian ATM lacks the N-terminal solenoid domains found in mammalian ATM (Figure S2A), where interactions with the MRN complex are thought to occur (Baretić and Williams, 2014; Stakyte *et al*., 2021). Intriguingly, planarians lack several components of the DDR and cell cycle checkpoints (Fig. EV1B), suggesting that while individual components of these pathways may be conserved, their collective functions, or their interactions, may have changed over the course of evolution (Grohme *et al*., 2018; Sahu *et al*., 2021). The findings in this study therefore lay a foundation to investigate whether conserved genes might have co-opted divergent roles in planarians to regulate cell death and DNA repair in response to genotoxic stress.

## Author contributions

Conceptualization: DAS, BDS, KTW, CEA

Methodology: DAS, BDS, KTW

Investigation: DAS, BDS, KTW, JKT, CEA

Visualization: DAS, KTW, CEA

Funding acquisition: CEA

Project administration: CEA

Supervision: CEA

Writing – original draft: DAS, KTW, CEA

Writing – review & editing: DAS, BDS, KTW, JKT, CEA

## Acknowledgments

We thank the Cornell University Biotechnology Resource Center’s Flow Cytometry, Imaging, and Genomics cores; members of the Adler laboratory for insight; R. Cerione, R. Weiss, E. Alani, G. Hollopeter, and M. Smolka for critical reading of the manuscript.

## Materials and Methods

### Planarian care

Asexual planarians from the *Schmidtea mediterranea* clonal line CIW4 were were maintained in a recirculating system containing Montjuïc salts (Arnold *et al*., 2016; Merryman, Sánchez Alvarado and Jenkin, 2018) with constant UV sanitization of water. Experimental animals were transferred to a static culture of planaria water supplemented with 50 μg/mL gentamicin sulfate. Animals were used for experiments 5-7 days after the last RNAi feed.

### Irradiation

A J.L. Shepherd & Associates Mark I-68 Irradiator was used to radiate planarians at the doses indicated in the figures. Animals were rinsed immediately after exposure and transferred into fresh planarian water on the following day. Radiated animals maintained longer than 2 days were rinsed on alternate days.

### RNA interference

RNAi was carried out as previously described, either using *in vitro* synthesized RNA (Rouhana *et al*., 2013) or by feeding animals with double stranded RNA-containing bacteria (Adler and Sánchez Alvarado, 2018). For *in vitro* synthesized RNAi, double stranded RNA (dsRNA) was generated using PCR products of genes as a template and mixed with a 4:1 liver:water paste containing 4μg of dsRNA per 10μL of liver. Animals were fed every other day, for a total of 6 feeds. For bacterial RNAi, cDNA for genes of interest were cloned into either pJC53.2 or T4P vector and transformed into HT115 competent cells. Cultures were grown in 2XYT broth at 37°C, shaking at 200–250 rpm until reaching an OD_600_ of 0.6, when IPTG was added to a final concentration of 1mM. Bacterial cultures were then spun down and resuspended in a 4:1 liver:water paste. For double RNAi, an equal amount of two bacterial liquid culture was mixed before spun down. Bacterial RNAi was administered for a total of 4 feeds every other day. Animals were used for experiments 5-7 days after the last RNAi feed. All experiments use *unc-22*(RNAi) as control. See Supplementary Table 4 for primer details of genes.

### qRT-PCR

For each biological replicate, 10 animals were homogenized in Trizol (Thermo Fisher 15596018) in Lysing Matrix D Tubes (MP Biomedicals 116913100) followed by homogenization in a Bead Bug homogenizer (Benchmark). RNA was extracted using a standard Trizol extraction protocol. cDNA was synthesized with Superscript VILO (Life Technologies 11754250). PCR mixes were made with TaqMan Gene Expression Master Mix (Life Technologies) and run using custom primers for *smedwi-1* (AI89MBJ) and GAPDH (AI6RPY3) (ThermoFisher) on an Applied Biosystems Viia7 Real Time PCR System. For each biological replicate, 3 technical replicates were performed and Ct values were averaged. Data was analyzed using Ct methods.

### Fixations

Animals were fixed and labeled as previously described (Pearson *et al*., 2009). Briefly, animals were processed in 7.5% N-acetyl-cysteine in PBS for 10 minutes and fixed in 4% paraformaldehyde for 30 minutes at room temperature. Worms were then rinsed twice with PBSTx (PBS + 0.3% Triton X-100), which was replaced with a prewarmed reduction solution (PBS containing 1% NP-40, 50mM DTT, and 0.5% SDS) and incubated at 37°C for 10 minutes. After rinsing twice with PBSTx, animals were dehydrated in serial solutions of 50:50 methanol:PBSTx and 100% methanol, and stored at −20°C.

### Whole-mount *in situ* hybridizations

Colorimetric *in situ* hybridizations were performed as described (Pearson *et al*., 2009) and fluorescent *in situ* hybridizations (King and Newmark, 2013) with some modifications. Briefly, fixed animals were rehydrated, bleached (5% formamide, 1.2% H_2_O_2_ in 0.5x SSC) and treated with proteinase K (4 μg/mL in PBSTx, Thermo Fisher 25530049) for 10 mins followed by fixation in 4% formaldehyde (10 mins). After 2 hours in pre-hybe, probes were added, and incubated overnight at 56°C. The next day, samples were washed 2x in wash hybe (5 min), 1:1 wash hybe:2X SSC-0.1% Tween 20 (10 min), and 2X SSC-0.1% Tween 20 (30 min), 0.2X SSC-0.1% Tween 20 (30 min) at 56°C followed by 3 × 10 min MABT washes (for colorimetric *in situs*) or PBSTx washes (fluorescent *in situs*) at room temperature. Animals were then placed in blocking solution (0.5% Roche Western Blocking Reagent and 5% inactivated horse serum diluted in MABT or PBSTx) for 2 hours at room temperature and incubated with an appropriate antibody overnight at 4°C: 1:3000 anti-DIG-AP (Roche 11093274910), 1:1000 anti-DIG-POD (Roche 11207733910) or 1:1000 anti-FITC-POD (Roche 11426346910) diluted in blocking solution. Subsequent washes and AP or tyramide development were performed as previously described. After development, animals were soaked in ScaleA2 (4M urea, 20% glycerol, 0.1% Triton X-100, 2.5% DABCO)^38^ and mounted in Aqua-Polymount (Polysciences Inc. 18606). Some *in situ* hybridizations were carried out in a CEM InSituPro Hybridization robot up to the development stage, using the same protocol. Animals were imaged either on a Leica M165F with a DFC7000T, a Zeiss 710 confocal microscope or a Keyence BZ-X800 microscope. For all *in situ* hybridizations, representative images are shown from a population of n≥10 animals.

### Phosphohistone H3 labeling

Animals were stained with phosphohistone H3 following fluorescent *in situ* hybridization. After inactivation of peroxidase with 4% H_2_O_2_ in PBSTx for 1 hour at RT, animals were rinsed in PBSTx (6x washes). Then, animals were incubated in anti-phosphohistone H3 (Ser10) antibody (Millipore Sigma, 05-817R-I) 1:300 in blocking solution for 2 days at 4°C. After washing off the primary with PBSTx, animals were incubated in a goat anti-rabbit-HRP secondary antibody (Thermo Fisher 31460) 1:1000 in PBSTx overnight at 4°C. Antibody was washed off with PBSTx and development was carried out as follows. Animals were pre-incubated in rhodamine tyramide (1:5000 in PBSTx) for 10 minutes, followed by development with 0.005% H_2_O_2_ in PBSTx for 15 minutes at room temperature. After development, samples were rinsed in PBSTx, then counterstained with DAPI (1:5000 in PBSTx, Thermo Scientific) before soaking in ScaleA2 and mounting in Aqua-Polymount. Animals were imaged on a Zeiss 710 confocal microscope or a Keyence BZ-X800 microscope.

### F-ara-EdU staining

Animals were soaked for 24 hours in 0.5 mg/mL F-ara-EdU (Sigma T511293) diluted in planaria water containing 3% DMSO. Soaking began either immediately after radiation or 24 hours later. Animals were fixed immediately after 24 hours in F-ara-EdU.

### F-ara-EdU detection

Fixed animals were rehydrated and bleached overnight in 6% H_2_O_2_ in PBSTx. Bleached animals were treated with proteinase K (10 μg/mL proteinase K + 0.1% SDS in PBSTx) for 15 min, and fixed in 4% formaldehyde in PBSTx for 10 min. F-ara-EdU was developed by soaking animals in development solution containing PBS + 1 mM CuS0_4_ and 100 μM Oregon Green 488 azide (Thermo Fisher O10180) with freshly made 100 mM ascorbic acid added immediately before administering. Animals were then incubated in this solution for 30 min in the dark. After rinsing with PBSTx, animals were post-fixed in 4% formaldehyde and rinsed 2x in PBSTx. Animals were then placed in K block (5% inactivated horse serum, 0.45% fish gelatin, 0.3% Triton-X and 0.05% Tween-20 diluted in PBS) at room temperature for 4 hours. This was followed by incubation with 1:1000 anti-Oregon Green-HRP (Thermo Fisher A21253) in K block overnight. The following day, after 6x PBSTx rinses, antibody was developed with FAM tyramide (1:2000 in Borate buffer) for 10 minutes, followed by development with 0.005% H_2_O_2_ in borate buffer for 15 minutes at room temperature.

### Flow cytometry and Annexin V staining

Flow cytometry of Hoechst-stained cells was conducted as previously described (Reddien *et al*., 2005; Hayashi *et al*., 2006) with minor modifications. Thirty animals per group were rinsed in CMFB (CMF containing 0.5% BSA) and dissected into pieces. Fragments were dissociated in 1:100 Liberase™ (2.5mg/mL, Roche 5401135001) in CMFB at 30°C, agitating at 300 rpm for 30 minutes on an Eppendorf ThermoMixer™. Fragments were triturated with a pipette periodically to aid dissociation. Dissociated cells were then diluted with equal volume of CMFB and centrifuged for 5 minutes at 500g. Pelleted cells were diluted in 1mL of CMFB and passed through a 30μm cell strainer (BD 340627). Cell density was assessed with an automated cell counter (Bio-Rad TC20™). 2.8×10^6^ cells/group were stained with 5μg/mL Hoechst 33342 (ThermoFisher H3570) diluted in CMFB for 70 mins in the dark with gentle rotation. Cells were spun at 500g for 5 minutes, then resuspended in Annexin V staining buffer (2μL Annexin V-APC (Thermo Fisher A35110) diluted in 100μL freshly made 1X Annexin V buffer from a 10X stock solution (0.1M HEPES pH 7.4, 1.4M NaCl, and 25 mM CaCl_2_)) and incubated for 15 minutes at RT. Then, 400μL of 1X Annexin V buffer with 1μg/mL Propidium Iodide (Sigma P4170) was added to each tube. Cells were analyzed on a BD FACS Symphony Analyzer and data was processed in FlowJo (TreeStar, Ashland, OR).

### Flow cytometry with DAPI staining

Thirty animals per condition were dissociated in 1:100 Liberase in CMFB as above. After dissociation and filtration, pelleted cells were fixed by gently resuspending in 4% paraformaldehyde in CMFB for 15 minutes with gentle agitation. Cells were then rinsed twice in PBS-1%BSA and permeabilized with 0.2% Tween-20 in 1X PBS with gentle rocking. Cells were then rinsed as before and incubated in 400μL of DAPI (1mg/mL) diluted 1:1000 in 1X PBS-0.1% Triton-X with gentle rocking at room temperature for 30 minutes in the dark. Samples were run on an Attune NxT analyzer and data was processed using the ModFit LT program.

### RNA-seq library preparation and analysis

RNA-seq libraries were generated from stem cells isolated from either *unc-22*(RNAi) or *atm*(RNAi) animals 1 day after radiation. Data was generated from three biological replicates with 5-10 animals. Stem cells were isolated as previously described (Peiris, García-Ojeda and Oviedo, 2016). Briefly, animals were dissociated as described above, cells were stained with Draq5 (ThermoFisher 65-0880-96) and Calcein-AM (ThermoFisher C3100MP), then sorted on a Sony MA900 Cell Sorter. RNA was isolated from a total of 100,000 stem cells per replicate. RNA-seq libraries were constructed using NEBNext® Ultra™ II Directional RNA Library Prep Kit for Illumina®, and run on a NextSeq500 sequencer. Sequencing reads were aligned using STAR aligner against schMed4-2018 genome assembly, and genes were annotated using the v6 Dresden transcriptome (Grohme *et al*., 2018; Rozanski *et al*., 2019). Differential gene expression analysis was carried out using DEseq2. Using the DEseq2 default parameters, the Wald test was used to compare between groups (unirradiated, *atm*(RNAi) vs. *control*(RNAi); or 1dpi, *atm*(RNAi) vs. *control*(RNAi)) and Benjamini-Hochberg for multi-testing correction (Love, Huber and Anders, 2014). For identifying differentially-expressed genes, |log_2_FC|>1 and adjusted p-value <0.01 were used as the cutoff. The upregulated genes in the comparison between *atm*(RNAi) vs. *control*(RNAi) at 1dpi are listed in Supplementary Table S1, and the downregulated genes are listed in Supplementary Table S2. For GO analysis, topGO was used for identifying enriched GO terms among upregulated or downregulated genes (Alexa and Rahnenführer, no date).

### Single-cell RNA-seq analysis

Single-cell RNA-seq analysis utilized published datasets generated from whole animals (Fincher *et al*., 2018). The analysis is conducted using the function “DotPlot” from Seurat (4.0.4) in R (4.0.5) (Hao *et al*., 2021). Following the original cell annotation from the publication, we divided the cells into two categories: those annotated as “‘Neoblast”, and the rest of the cells, which were annotated here as “Differentiated cells”. Dot size represents the percentage of cells in each category that expressed at least 1 normalized count. The color scale represents the average expression of each annotation (neoblast or differentiated cells), calculated across all cells included in the analysis.

### Image quantification

All image quantification was carried out in Fiji (Schindelin *et al*., 2012). For quantification of fluorescent confocal images of *piwi-1*, phosphohistone-H3 and EdU, maximum projections of confocal images were generated. For EdU and pH3, images were thresholded and counted using the Analyze tool in Fiji. DAPI signal was used to threshold images and measure total area of the animal. Quantification of the number of *piwi-1* cells from colourimetric and fluorescent images was done manually using the CellCounter Plugin. To calculate the percentage of animal area occupied by fluorescent signal for *piwi-1*, we determined the area occupied by *piwi-1* by thresholding, and divided this by the total area of the animal, calculated based on the DAPI signal.

### Statistical analysis and reproducibility

Statistical analysis was performed using PRISM-Graphpad version 9. Statistical details of all experiments are noted in each figure legend. Data represents ≥ 2 independent experiments.

## Data and materials availability

All data are available upon request. RNAseq data has been deposited into NCBI (Accession number GSE190454).

---

**Fig. EV1.**
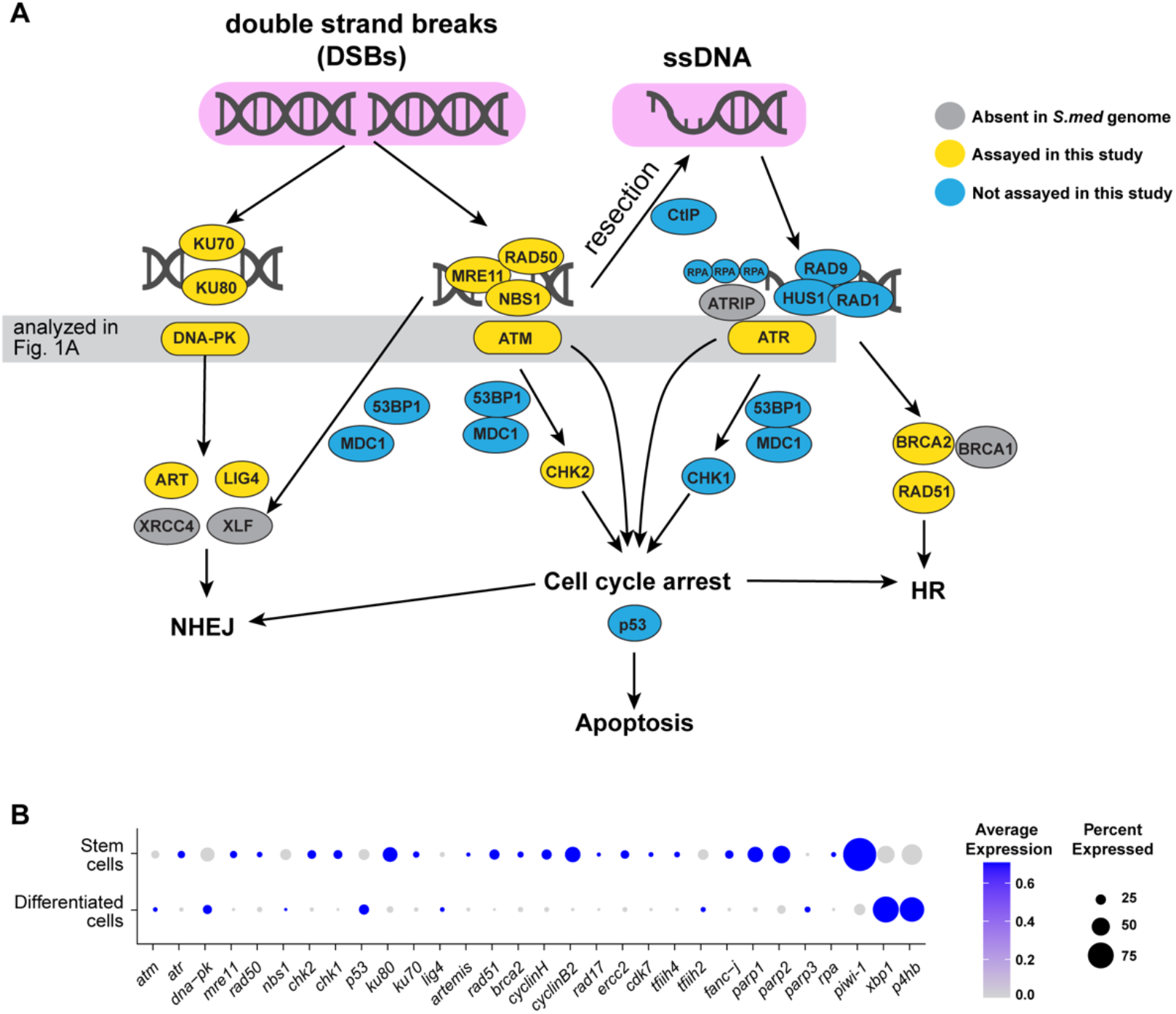
Schematic of DNA damage response pathways. **(A**) Schematic of DNA damage response pathway in humans. Homologs of proteins highlighted in yellow were tested in this study. Those in gray are absent in the *Schmidtea mediterranea* genome (Grohme et al., 2018; Sahu et al., 2021), and those in blue were not examined here. The interactions and relationships among the proteins are unverified in planarians. The *S. mediterranea* p53 homolog is not involved in DNA-damage induced apoptosis (Wendt et al., 2021). **(B**) Dot plot showing expression levels of upregulated genes, and those identified by RNAseq (Fig. S3) and DNA damage response genes in stem cells and differentiated cells in single-cell RNAseq data (Fincher et al., 2018). Cells were grouped into two categories (stem cells vs differentiated cells); in each population, the average expression and percent of cells expressing a given transcript were calculated. *piwi-1* expression is shown as a control for stem cells, and *xbp1* and *p4hb* are included as references for differentiated cells.

**Fig. EV2.**
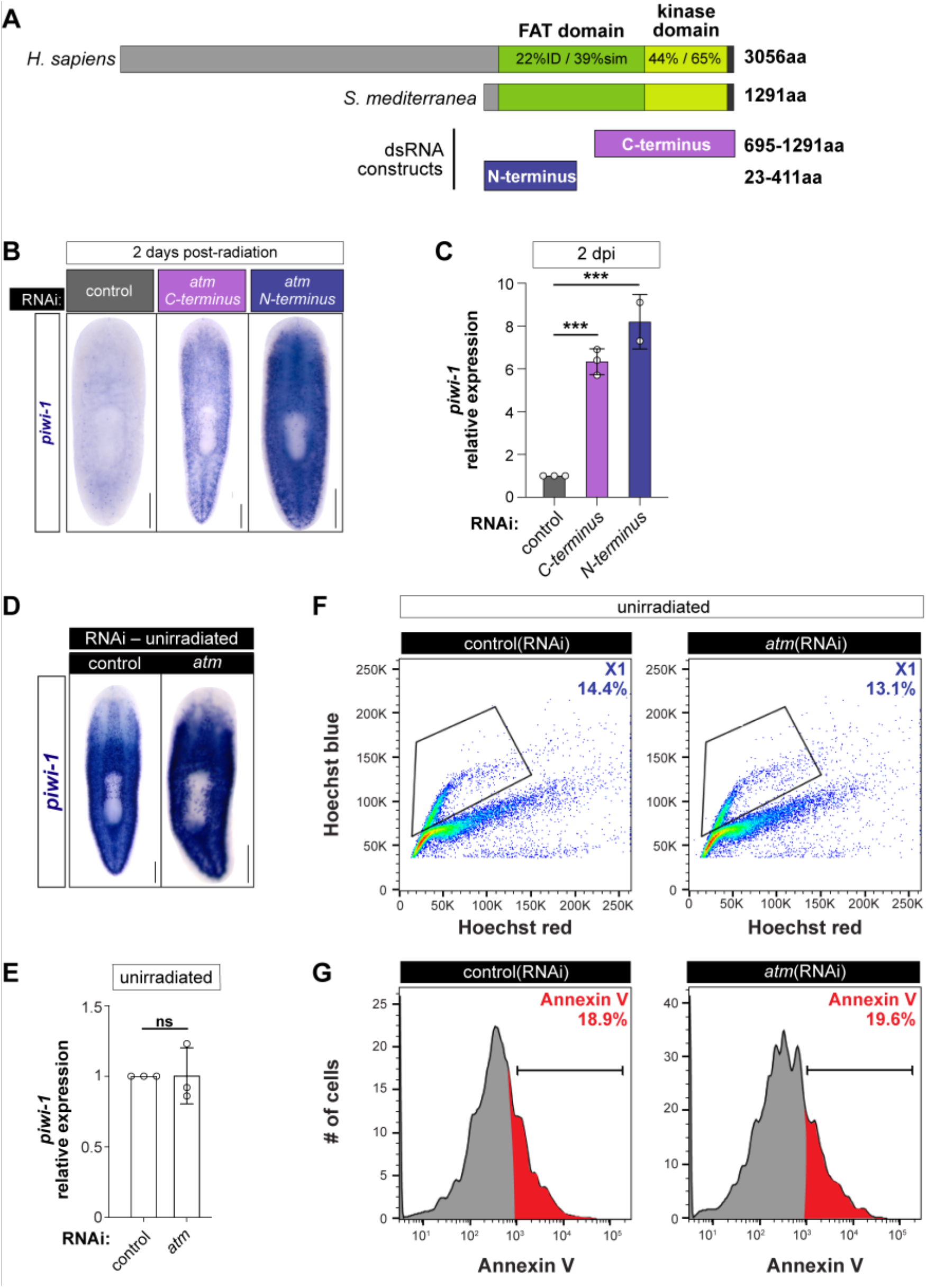
ATM knockdown acts only after radiation. **(A**) Schematic of planarian and human ATM showing domain conservation. Blue and purple boxes show non-overlapping RNAi constructs tested. **(B**) *piwi-1* WISH of RNAi animals as indicated, 2 dpi. Scale bars=250μm. **(C**) qRT-PCR of *piwi-1* relative to GAPDH in RNAi animals. Statistics represent ordinary one-way ANOVA compared to controls, ***P<0.0004. **(D**) *piwi-1* WISH of unirradiated RNAi animals. Scale bars=250μm. **(E**) qRT-PCR of *piwi-1* relative to GAPDH in unirradiated RNAi animals. Statistics represent unpaired t-test; ns=not significant. In C and E, data are averages ± SD; circles represent biological replicates. **(F**) Source data for Fig. 1D. FACS plots of Hoechst-stained X1 cells from unirradiated control and *atm(RNAi*) animals. Black polygon=X1 cells. **(G**) Source data for Fig. 1G. Histograms of X1s from (E) stained with Annexin V. Percent Annexin-V-positive X1s in red. Brackets and red shaded area=Annexin V mean fluorescent intensity staining >10^3^.

**Fig. EV3.**
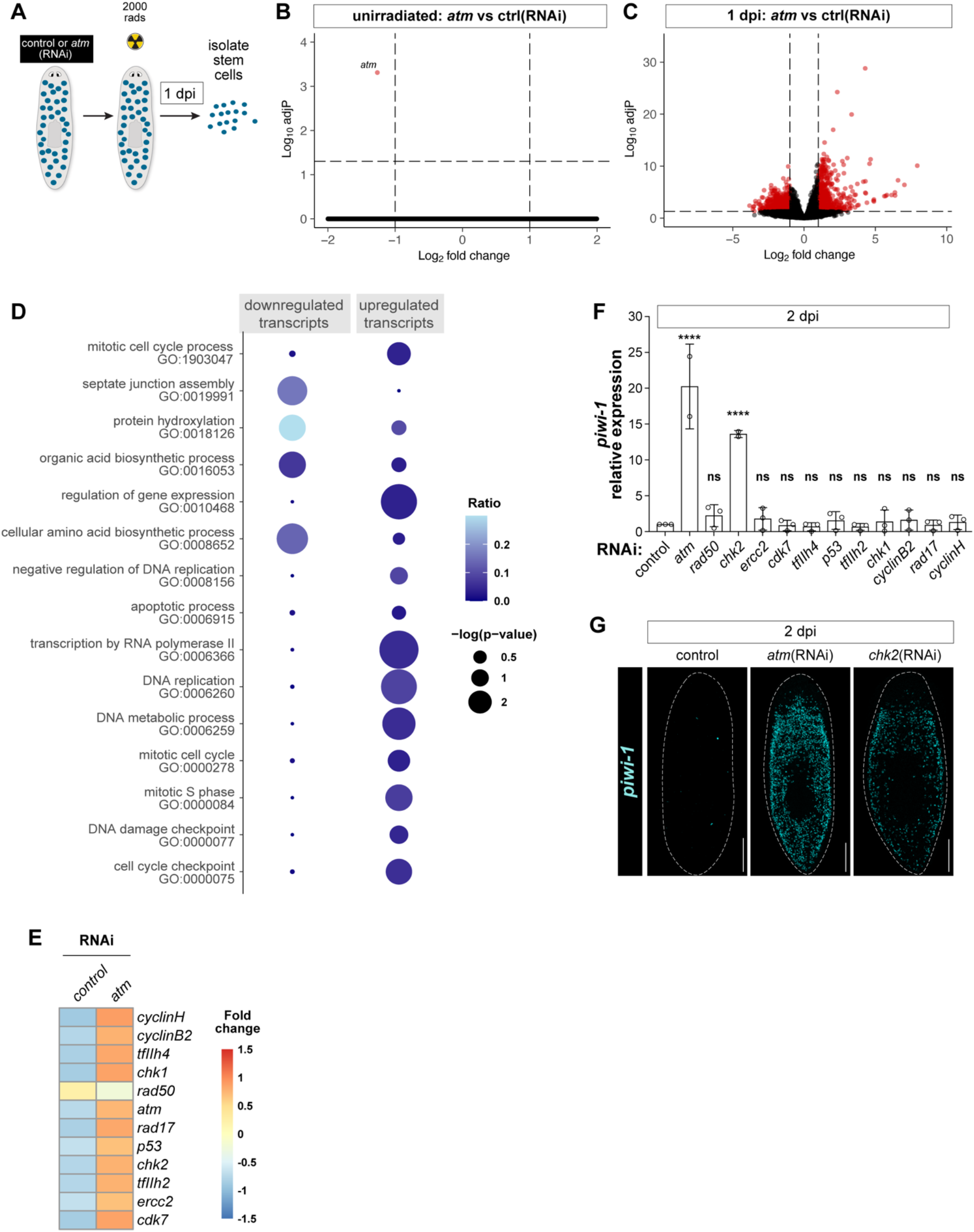
RNA-seq analysis of stem cells following *atm*(RNAi) **(A**) Experimental schematic showing strategy to purify stem cells prior to RNA-seq. **(B**) Volcano plot of differentially expressed transcripts in unirradiated RNAi animals. Red dots indicate transcripts with |log_2_ fold change|<1 and adjusted p-value <0.01. **(C**) Volcano plot of differentially expressed transcripts in RNAi animals 1 day after radiation exposure. Red dots indicate transcripts with |log_2_ fold change|<1 and adjusted p-value <0.01. **(D**) Gene ontology analysis of upregulated transcripts from C. **(E**) Heat map of upregulated transcripts used in RNAi screen in Fig. S2F. **(F**) qRT-PCR of *piwi-1* relative to GAPDH in RNAi animals as indicated, 2dpi. Data are averages ± SD; circles represent biological replicates; statistics represent ordinary one-way ANOVA compared to controls, ****P<0.0001, ns=not significant. **(G**) *piwi-1* fluorescent *in situ* hybridization (FISH) of RNAi animals, 2 dpi.

**Fig. EV4.**
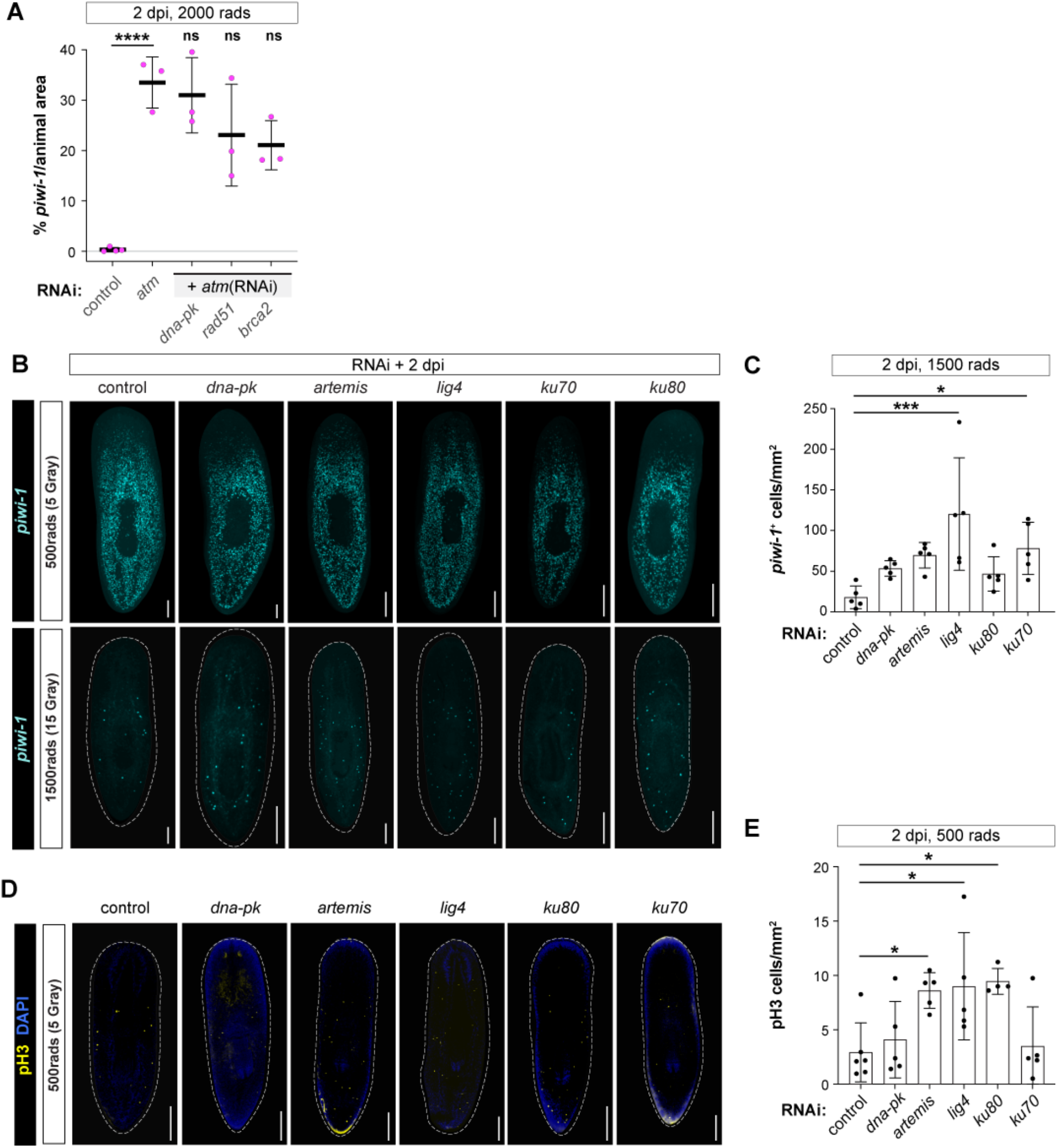
NHEJ is dispensable for stem cell persistence after radiation exposure. **(A**) Percentage of animal area occupied by *piwi-1* FISH divided by total animal area, in radiated RNAi animals as indicated. **(B**) Confocal images of *piwi-1* FISH (cyan) after RNAi of NHEJ genes 2 days after exposure to either 500 or 1500 rads. **(C**) Quantification by area of *piwi-1^+^* cells in RNAi animals (from panel **B**) after 1500 rads exposure, 2dpi. **(D**) Confocal images of animals stained with pH3 (yellow) and DAPI (DNA, blue) after RNAi of NHEJ genes, 2 days after exposure to 500 rads. **(E**) Quantification by area of pH3+ mitotic nuclei in RNAi animals (from panel **D**) after exposure to 500 rads. Data are averages ± SD; circles represent biological replicates; statistics represent ordinary one-way ANOVA compared to *atm*(RNAi) in **(A)**; controls in **(C)** and **(E)**; *P<0.05, ***P<0.0005, ****P<0.0001, ns= not significant. Scale bars=250μm.

