## Supplemental Tables 1-4 for "Inhibition of the ATM kinase rescues planarian regeneration after lethal radiation"

**Table S1: List of upregulated genes in irradiated stem cells (1dpi), atm(RNAi) vs. control(RNAi).**
ID: Planmine transcript ID from dd_Smed_v6 transcriptome.
baseMean: mean of normalized counts for all samples;
log2FC: log2 fold change;
lfcSE: standard error of log2 fold change;
stat: Wald statistic;
pvalue: Wald test p-value;
padj: BH adjusted p-values

Only genes with log2FC >1 and adjusted p-value <0.01 are listed here.

| ID | baseMean | log2FC | lfcSE | stat | pvalue | padj |
|---|---|---|---|---|---|---|
| dd_Smed_v6_15107_0_1 | 122.43 | 7.94 | 1.07 | 7.40 | 1.39E-13 | 8.14E-11 |
| dd_Smed_v6_52262_0_1 | 60.87 | 7.03 | 1.19 | 5.93 | 3.02E-09 | 3.99E-07 |
| dd_Smed_v6_21717_0_1 | 106.67 | 6.59 | 1.05 | 6.29 | 3.13E-10 | 5.33E-08 |
| dd_Smed_v6_28888_0_1 | 16.81 | 6.38 | 1.30 | 4.89 | 9.98E-07 | 4.49E-05 |
| dd_Smed_v6_50870_0_1 | 16.84 | 6.37 | 1.22 | 5.23 | 1.74E-07 | 1.15E-05 |
| dd_Smed_v6_46372_0_1 | 14.48 | 6.14 | 1.24 | 4.94 | 7.62E-07 | 3.65E-05 |
| dd_Smed_v6_56209_0_1 | 17.96 | 5.80 | 1.17 | 4.95 | 7.42E-07 | 3.58E-05 |
| dd_Smed_v6_27803_0_1 | 19.00 | 5.72 | 1.18 | 4.84 | 1.27E-06 | 5.39E-05 |
| dd_Smed_v6_16466_0_1 | 27.12 | 5.63 | 1.17 | 4.82 | 1.44E-06 | 5.96E-05 |
| dd_Smed_v6_64323_0_1 | 18.16 | 4.90 | 1.17 | 4.17 | 3.05E-05 | 6.23E-04 |
| dd_Smed_v6_17951_0_1 | 45.25 | 4.90 | 1.11 | 4.43 | 9.60E-06 | 2.60E-04 |
| dd_Smed_v6_49095_0_1 | 26.13 | 4.87 | 1.14 | 4.28 | 1.87E-05 | 4.36E-04 |
| dd_Smed_v6_31975_0_1 | 60.98 | 4.66 | 0.59 | 7.87 | 3.60E-15 | 4.79E-12 |
| dd_Smed_v6_17938_0_1 | 112.42 | 4.59 | 0.61 | 7.55 | 4.49E-14 | 3.46E-11 |
| dd_Smed_v6_17770_0_1 | 21.70 | 4.41 | 1.10 | 4.01 | 6.16E-05 | 1.07E-03 |
| dd_Smed_v6_18839_0_1 | 29.74 | 4.33 | 0.84 | 5.13 | 2.89E-07 | 1.71E-05 |
| dd_Smed_v6_54952_0_1 | 46.78 | 4.30 | 0.69 | 6.22 | 4.91E-10 | 8.08E-08 |
| dd_Smed_v6_25682_0_1 | 28.14 | 4.29 | 0.83 | 5.20 | 1.98E-07 | 1.27E-05 |
| dd_Smed_v6_10988_0_1 | 911.23 | 4.29 | 0.35 | 12.10 | 1.05E-33 | 1.54E-29 |
| dd_Smed_v6_18815_0_1 | 51.03 | 4.15 | 0.67 | 6.19 | 6.09E-10 | 9.81E-08 |
| dd_Smed_v6_32328_0_1 | 28.93 | 3.82 | 1.04 | 3.68 | 2.35E-04 | 2.96E-03 |
| dd_Smed_v6_29386_0_1 | 15.15 | 3.74 | 1.03 | 3.62 | 2.90E-04 | 3.44E-03 |
| dd_Smed_v6_18297_0_1 | 71.35 | 3.74 | 0.62 | 6.01 | 1.89E-09 | 2.59E-07 |
| dd_Smed_v6_12517_0_1 | 44.98 | 3.62 | 0.73 | 4.98 | 6.22E-07 | 3.13E-05 |
| dd_Smed_v6_11118_0_1 | 18.76 | 3.58 | 1.09 | 3.29 | 1.01E-03 | 8.62E-03 |
| dd_Smed_v6_32297_0_1 | 64.57 | 3.55 | 0.72 | 4.95 | 7.35E-07 | 3.58E-05 |
| dd_Smed_v6_11654_0_1 | 25.96 | 3.52 | 0.83 | 4.24 | 2.22E-05 | 4.92E-04 |
| dd_Smed_v6_22479_0_1 | 87.42 | 3.40 | 0.64 | 5.34 | 9.09E-08 | 6.83E-06 |
| dd_Smed_v6_29468_0_1 | 58.90 | 3.35 | 0.76 | 4.42 | 9.99E-06 | 2.69E-04 |
| dd_Smed_v6_16549_0_1 | 437.01 | 3.33 | 0.33 | 10.18 | 2.34E-24 | 1.14E-20 |

| | | | | | | |
|---|---|---|---|---|---|---|
| dd_Smed_v6_15045_0_1 | 19.63 | 3.33 | 0.94 | 3.56 | 3.71E-04 | 4.14E-03 |
| dd_Smed_v6_44311_0_1 | 56.28 | 3.28 | 0.57 | 5.75 | 8.90E-09 | 9.83E-07 |
| dd_Smed_v6_15035_0_1 | 65.44 | 3.23 | 0.67 | 4.79 | 1.70E-06 | 6.81E-05 |
| dd_Smed_v6_14669_0_1 | 25.76 | 3.22 | 0.82 | 3.93 | 8.48E-05 | 1.37E-03 |
| dd_Smed_v6_25053_0_1 | 17.44 | 3.19 | 0.93 | 3.43 | 6.11E-04 | 6.02E-03 |
| dd_Smed_v6_33402_0_1 | 28.14 | 3.13 | 0.77 | 4.09 | 4.28E-05 | 8.08E-04 |
| dd_Smed_v6_4075_0_1 | 61.74 | 3.09 | 0.67 | 4.60 | 4.26E-06 | 1.41E-04 |
| dd_Smed_v6_12368_0_1 | 435.91 | 3.03 | 0.38 | 7.98 | 1.51E-15 | 2.76E-12 |
| dd_Smed_v6_20374_0_1 | 23.73 | 3.00 | 0.78 | 3.83 | 1.30E-04 | 1.94E-03 |
| dd_Smed_v6_4414_0_1 | 64.18 | 2.98 | 0.65 | 4.58 | 4.59E-06 | 1.51E-04 |
| dd_Smed_v6_22968_0_1 | 23.75 | 2.89 | 0.83 | 3.46 | 5.38E-04 | 5.48E-03 |
| dd_Smed_v6_13860_0_1 | 121.39 | 2.88 | 0.51 | 5.61 | 1.99E-08 | 2.00E-06 |
| dd_Smed_v6_27975_0_1 | 23.78 | 2.82 | 0.86 | 3.26 | 1.10E-03 | 9.26E-03 |
| dd_Smed_v6_23603_0_1 | 26.85 | 2.81 | 0.83 | 3.37 | 7.52E-04 | 6.98E-03 |
| dd_Smed_v6_7227_0_1 | 357.82 | 2.79 | 0.41 | 6.83 | 8.51E-12 | 2.31E-09 |
| dd_Smed_v6_25375_0_1 | 61.48 | 2.79 | 0.58 | 4.78 | 1.76E-06 | 7.00E-05 |
| dd_Smed_v6_16426_0_1 | 225.86 | 2.77 | 0.37 | 7.46 | 8.71E-14 | 5.32E-11 |
| dd_Smed_v6_7688_0_1 | 68.62 | 2.75 | 0.56 | 4.95 | 7.43E-07 | 3.58E-05 |
| dd_Smed_v6_26182_0_1 | 52.99 | 2.75 | 0.54 | 5.12 | 3.13E-07 | 1.81E-05 |
| dd_Smed_v6_9151_0_1 | 235.59 | 2.69 | 0.35 | 7.72 | 1.20E-14 | 1.10E-11 |
| dd_Smed_v6_58387_0_1 | 27.64 | 2.64 | 0.71 | 3.70 | 2.14E-04 | 2.77E-03 |
| dd_Smed_v6_17071_0_1 | 140.54 | 2.64 | 0.57 | 4.64 | 3.49E-06 | 1.23E-04 |
| dd_Smed_v6_20355_0_1 | 72.80 | 2.61 | 0.57 | 4.56 | 5.16E-06 | 1.63E-04 |
| dd_Smed_v6_43024_0_1 | 20.80 | 2.60 | 0.77 | 3.38 | 7.21E-04 | 6.77E-03 |
| dd_Smed_v6_13690_0_1 | 42.79 | 2.59 | 0.65 | 3.97 | 7.07E-05 | 1.19E-03 |
| dd_Smed_v6_30393_0_1 | 26.78 | 2.55 | 0.63 | 4.02 | 5.82E-05 | 1.02E-03 |
| dd_Smed_v6_18122_0_1 | 61.76 | 2.50 | 0.51 | 4.94 | 7.84E-07 | 3.72E-05 |
| dd_Smed_v6_32569_0_1 | 33.54 | 2.49 | 0.74 | 3.39 | 7.01E-04 | 6.63E-03 |
| dd_Smed_v6_36433_0_1 | 51.59 | 2.46 | 0.66 | 3.71 | 2.09E-04 | 2.73E-03 |
| dd_Smed_v6_18317_0_1 | 70.20 | 2.41 | 0.47 | 5.11 | 3.23E-07 | 1.86E-05 |
| dd_Smed_v6_11270_0_1 | 193.00 | 2.40 | 0.36 | 6.68 | 2.31E-11 | 5.29E-09 |
| dd_Smed_v6_8720_0_1 | 964.63 | 2.36 | 0.54 | 4.42 | 9.90E-06 | 2.67E-04 |
| dd_Smed_v6_12204_0_1 | 36.04 | 2.35 | 0.66 | 3.58 | 3.48E-04 | 3.96E-03 |
| dd_Smed_v6_14049_0_1 | 95.58 | 2.34 | 0.42 | 5.53 | 3.15E-08 | 2.95E-06 |
| dd_Smed_v6_13215_0_1 | 219.75 | 2.34 | 0.37 | 6.29 | 3.27E-10 | 5.50E-08 |
| dd_Smed_v6_36340_0_1 | 30.10 | 2.33 | 0.71 | 3.29 | 9.87E-04 | 8.51E-03 |
| dd_Smed_v6_11867_0_1 | 1286.08 | 2.33 | 0.21 | 11.14 | 7.64E-29 | 5.60E-25 |
| dd_Smed_v6_33557_0_1 | 38.30 | 2.32 | 0.60 | 3.84 | 1.24E-04 | 1.88E-03 |
| dd_Smed_v6_41385_0_1 | 32.75 | 2.29 | 0.55 | 4.18 | 2.93E-05 | 6.06E-04 |
| dd_Smed_v6_30887_0_1 | 63.70 | 2.27 | 0.59 | 3.84 | 1.24E-04 | 1.89E-03 |
| dd_Smed_v6_62956_0_1 | 42.77 | 2.25 | 0.53 | 4.26 | 2.06E-05 | 4.65E-04 |
| dd_Smed_v6_15555_0_1 | 174.35 | 2.25 | 0.35 | 6.41 | 1.49E-10 | 2.73E-08 |
| dd_Smed_v6_21432_0_1 | 83.14 | 2.24 | 0.41 | 5.52 | 3.48E-08 | 3.16E-06 |

| | | | | | | |
|---|---|---|---|---|---|---|
| dd_Smed_v6_12909_0_1 | 65.36 | 2.19 | 0.54 | 4.09 | 4.37E-05 | 8.20E-04 |
| dd_Smed_v6_12332_0_1 | 807.94 | 2.19 | 0.31 | 7.02 | 2.26E-12 | 7.35E-10 |
| dd_Smed_v6_12873_0_1 | 128.37 | 2.14 | 0.38 | 5.57 | 2.48E-08 | 2.39E-06 |
| dd_Smed_v6_25428_0_1 | 87.19 | 2.12 | 0.58 | 3.68 | 2.30E-04 | 2.91E-03 |
| dd_Smed_v6_19890_0_1 | 86.78 | 2.11 | 0.57 | 3.68 | 2.36E-04 | 2.97E-03 |
| dd_Smed_v6_18582_0_1 | 80.73 | 2.11 | 0.49 | 4.33 | 1.48E-05 | 3.66E-04 |
| dd_Smed_v6_42967_0_1 | 93.11 | 2.08 | 0.54 | 3.87 | 1.08E-04 | 1.68E-03 |
| dd_Smed_v6_15123_0_1 | 454.53 | 2.07 | 0.28 | 7.47 | 8.12E-14 | 5.18E-11 |
| dd_Smed_v6_17433_0_1 | 66.03 | 2.05 | 0.54 | 3.79 | 1.53E-04 | 2.19E-03 |
| dd_Smed_v6_3026_1_1 | 33.59 | 2.05 | 0.59 | 3.46 | 5.41E-04 | 5.49E-03 |
| dd_Smed_v6_2016_0_1 | 6934.73 | 2.04 | 0.22 | 9.47 | 2.77E-21 | 1.01E-17 |
| dd_Smed_v6_36562_0_1 | 32.59 | 2.02 | 0.62 | 3.27 | 1.09E-03 | 9.21E-03 |
| dd_Smed_v6_15178_0_1 | 323.92 | 2.02 | 0.41 | 4.86 | 1.15E-06 | 5.02E-05 |
| dd_Smed_v6_48433_0_1 | 44.46 | 2.02 | 0.60 | 3.34 | 8.34E-04 | 7.54E-03 |
| dd_Smed_v6_11959_0_1 | 357.19 | 1.96 | 0.33 | 6.02 | 1.70E-09 | 2.37E-07 |
| dd_Smed_v6_8102_0_1 | 215.64 | 1.95 | 0.33 | 6.00 | 1.96E-09 | 2.65E-07 |
| dd_Smed_v6_7850_0_1 | 341.45 | 1.94 | 0.29 | 6.81 | 9.57E-12 | 2.50E-09 |
| dd_Smed_v6_9095_0_1 | 66.64 | 1.92 | 0.49 | 3.96 | 7.37E-05 | 1.23E-03 |
| dd_Smed_v6_29867_0_1 | 102.16 | 1.92 | 0.54 | 3.53 | 4.20E-04 | 4.54E-03 |
| dd_Smed_v6_16388_0_1 | 64.55 | 1.92 | 0.48 | 4.00 | 6.32E-05 | 1.09E-03 |
| dd_Smed_v6_14042_0_1 | 215.36 | 1.91 | 0.33 | 5.75 | 8.92E-09 | 9.83E-07 |
| dd_Smed_v6_17201_0_1 | 115.68 | 1.89 | 0.46 | 4.07 | 4.74E-05 | 8.74E-04 |
| dd_Smed_v6_19900_0_1 | 126.31 | 1.89 | 0.27 | 6.90 | 5.21E-12 | 1.47E-09 |
| dd_Smed_v6_15343_0_1 | 145.49 | 1.88 | 0.52 | 3.64 | 2.70E-04 | 3.27E-03 |
| dd_Smed_v6_32522_0_1 | 43.93 | 1.87 | 0.48 | 3.88 | 1.05E-04 | 1.65E-03 |
| dd_Smed_v6_11494_0_1 | 1345.51 | 1.85 | 0.28 | 6.71 | 2.00E-11 | 4.73E-09 |
| dd_Smed_v6_13871_0_1 | 169.00 | 1.84 | 0.31 | 5.92 | 3.14E-09 | 4.00E-07 |
| dd_Smed_v6_8311_0_1 | 51.09 | 1.84 | 0.41 | 4.48 | 7.41E-06 | 2.18E-04 |
| dd_Smed_v6_13780_0_1 | 117.14 | 1.83 | 0.29 | 6.40 | 1.59E-10 | 2.87E-08 |
| dd_Smed_v6_13567_0_1 | 252.08 | 1.82 | 0.30 | 6.05 | 1.46E-09 | 2.11E-07 |
| dd_Smed_v6_11423_0_1 | 84.97 | 1.82 | 0.48 | 3.81 | 1.41E-04 | 2.06E-03 |
| dd_Smed_v6_21908_0_1 | 104.69 | 1.82 | 0.41 | 4.44 | 9.17E-06 | 2.53E-04 |
| dd_Smed_v6_18547_0_1 | 113.39 | 1.81 | 0.35 | 5.13 | 2.94E-07 | 1.73E-05 |
| dd_Smed_v6_16034_0_1 | 207.29 | 1.80 | 0.39 | 4.63 | 3.70E-06 | 1.28E-04 |
| dd_Smed_v6_15471_0_1 | 231.54 | 1.80 | 0.41 | 4.44 | 8.84E-06 | 2.46E-04 |
| dd_Smed_v6_17385_0_1 | 90.63 | 1.79 | 0.48 | 3.74 | 1.86E-04 | 2.53E-03 |
| dd_Smed_v6_16241_0_1 | 64.98 | 1.79 | 0.53 | 3.38 | 7.36E-04 | 6.86E-03 |
| dd_Smed_v6_15230_0_1 | 94.46 | 1.77 | 0.50 | 3.53 | 4.16E-04 | 4.51E-03 |
| dd_Smed_v6_12461_0_1 | 1037.15 | 1.77 | 0.26 | 6.69 | 2.20E-11 | 5.11E-09 |
| dd_Smed_v6_9087_0_1 | 378.05 | 1.75 | 0.35 | 5.05 | 4.31E-07 | 2.33E-05 |
| dd_Smed_v6_17701_0_1 | 67.19 | 1.74 | 0.46 | 3.80 | 1.44E-04 | 2.09E-03 |
| dd_Smed_v6_21468_0_1 | 55.77 | 1.74 | 0.42 | 4.17 | 3.05E-05 | 6.23E-04 |
| dd_Smed_v6_14459_0_1 | 112.60 | 1.73 | 0.40 | 4.30 | 1.72E-05 | 4.16E-04 |

| | | | | | | |
|---|---|---|---|---|---|---|
| dd_Smed_v6_14387_0_1 | 336.09 | 1.73 | 0.22 | 7.74 | 9.63E-15 | 9.65E-12 |
| dd_Smed_v6_17384_0_1 | 50.56 | 1.73 | 0.48 | 3.63 | 2.82E-04 | 3.37E-03 |
| dd_Smed_v6_13772_0_1 | 296.76 | 1.72 | 0.36 | 4.80 | 1.60E-06 | 6.51E-05 |
| dd_Smed_v6_9050_0_1 | 606.75 | 1.72 | 0.22 | 7.81 | 5.91E-15 | 6.66E-12 |
| dd_Smed_v6_5816_0_1 | 1214.40 | 1.71 | 0.36 | 4.73 | 2.25E-06 | 8.65E-05 |
| dd_Smed_v6_10422_0_1 | 77.41 | 1.71 | 0.43 | 3.94 | 8.08E-05 | 1.32E-03 |
| dd_Smed_v6_9899_0_1 | 140.41 | 1.71 | 0.39 | 4.35 | 1.37E-05 | 3.46E-04 |
| dd_Smed_v6_7534_0_1 | 365.47 | 1.70 | 0.28 | 6.15 | 7.61E-10 | 1.19E-07 |
| dd_Smed_v6_8969_0_1 | 383.36 | 1.69 | 0.23 | 7.25 | 4.24E-13 | 1.83E-10 |
| dd_Smed_v6_14224_0_1 | 63.56 | 1.69 | 0.47 | 3.58 | 3.50E-04 | 3.97E-03 |
| dd_Smed_v6_17143_0_1 | 85.22 | 1.69 | 0.46 | 3.64 | 2.72E-04 | 3.28E-03 |
| dd_Smed_v6_59138_0_1 | 38.38 | 1.68 | 0.45 | 3.73 | 1.92E-04 | 2.58E-03 |
| dd_Smed_v6_14888_0_1 | 525.63 | 1.68 | 0.35 | 4.85 | 1.21E-06 | 5.21E-05 |
| dd_Smed_v6_19563_0_1 | 245.26 | 1.68 | 0.41 | 4.10 | 4.13E-05 | 7.88E-04 |
| dd_Smed_v6_11973_0_1 | 522.83 | 1.68 | 0.28 | 6.09 | 1.15E-09 | 1.71E-07 |
| dd_Smed_v6_12841_0_1 | 857.59 | 1.67 | 0.33 | 5.08 | 3.72E-07 | 2.08E-05 |
| dd_Smed_v6_12814_0_1 | 175.65 | 1.67 | 0.37 | 4.51 | 6.42E-06 | 1.94E-04 |
| dd_Smed_v6_5128_0_1 | 677.08 | 1.67 | 0.31 | 5.44 | 5.32E-08 | 4.51E-06 |
| dd_Smed_v6_31977_0_1 | 91.21 | 1.66 | 0.38 | 4.33 | 1.52E-05 | 3.73E-04 |
| dd_Smed_v6_12831_0_1 | 322.86 | 1.66 | 0.32 | 5.15 | 2.65E-07 | 1.62E-05 |
| dd_Smed_v6_9381_0_1 | 450.13 | 1.65 | 0.25 | 6.46 | 1.06E-10 | 2.08E-08 |
| dd_Smed_v6_9250_0_1 | 1087.18 | 1.65 | 0.25 | 6.47 | 9.62E-11 | 1.93E-08 |
| dd_Smed_v6_16242_0_1 | 381.18 | 1.64 | 0.35 | 4.70 | 2.66E-06 | 9.82E-05 |
| dd_Smed_v6_6046_0_1 | 54.48 | 1.63 | 0.48 | 3.40 | 6.73E-04 | 6.42E-03 |
| dd_Smed_v6_14814_0_1 | 140.06 | 1.63 | 0.32 | 5.08 | 3.72E-07 | 2.08E-05 |
| dd_Smed_v6_6147_0_1 | 337.20 | 1.63 | 0.26 | 6.34 | 2.33E-10 | 4.07E-08 |
| dd_Smed_v6_15776_0_1 | 191.38 | 1.62 | 0.36 | 4.52 | 6.29E-06 | 1.90E-04 |
| dd_Smed_v6_33634_0_1 | 66.04 | 1.62 | 0.46 | 3.56 | 3.65E-04 | 4.08E-03 |
| dd_Smed_v6_18101_0_1 | 525.28 | 1.62 | 0.30 | 5.33 | 9.65E-08 | 7.17E-06 |
| dd_Smed_v6_13579_0_1 | 384.01 | 1.62 | 0.36 | 4.46 | 8.33E-06 | 2.35E-04 |
| dd_Smed_v6_14052_0_1 | 193.37 | 1.61 | 0.34 | 4.70 | 2.57E-06 | 9.55E-05 |
| dd_Smed_v6_10541_0_1 | 187.46 | 1.61 | 0.24 | 6.68 | 2.43E-11 | 5.47E-09 |
| dd_Smed_v6_12569_0_1 | 340.30 | 1.60 | 0.27 | 6.02 | 1.69E-09 | 2.37E-07 |
| dd_Smed_v6_5419_0_1 | 3132.51 | 1.60 | 0.20 | 8.05 | 8.05E-16 | 1.69E-12 |
| dd_Smed_v6_12818_0_1 | 444.84 | 1.59 | 0.28 | 5.60 | 2.15E-08 | 2.12E-06 |
| dd_Smed_v6_18174_0_1 | 360.44 | 1.58 | 0.31 | 5.06 | 4.12E-07 | 2.25E-05 |
| dd_Smed_v6_19513_0_1 | 95.31 | 1.58 | 0.36 | 4.40 | 1.08E-05 | 2.85E-04 |
| dd_Smed_v6_8324_0_1 | 335.80 | 1.57 | 0.23 | 6.83 | 8.48E-12 | 2.31E-09 |
| dd_Smed_v6_11422_0_1 | 467.10 | 1.57 | 0.23 | 6.96 | 3.29E-12 | 1.00E-09 |
| dd_Smed_v6_8483_0_1 | 1479.55 | 1.57 | 0.24 | 6.46 | 1.06E-10 | 2.08E-08 |
| dd_Smed_v6_16224_0_1 | 203.57 | 1.57 | 0.32 | 4.95 | 7.43E-07 | 3.58E-05 |
| dd_Smed_v6_9675_0_1 | 265.69 | 1.57 | 0.34 | 4.60 | 4.20E-06 | 1.40E-04 |
| dd_Smed_v6_12400_0_1 | 130.12 | 1.57 | 0.30 | 5.22 | 1.82E-07 | 1.20E-05 |

| | | | | | | |
|---|---|---|---|---|---|---|
| dd_Smed_v6_7870_0_1 | 786.41 | 1.56 | 0.30 | 5.18 | 2.22E-07 | 1.40E-05 |
| dd_Smed_v6_17726_0_1 | 176.55 | 1.55 | 0.37 | 4.18 | 2.90E-05 | 6.01E-04 |
| dd_Smed_v6_17673_0_1 | 216.74 | 1.54 | 0.29 | 5.28 | 1.30E-07 | 9.25E-06 |
| dd_Smed_v6_9656_0_1 | 355.43 | 1.54 | 0.22 | 6.99 | 2.78E-12 | 8.86E-10 |
| dd_Smed_v6_11441_0_1 | 352.76 | 1.54 | 0.21 | 7.31 | 2.73E-13 | 1.29E-10 |
| dd_Smed_v6_12881_0_1 | 1854.82 | 1.54 | 0.28 | 5.53 | 3.19E-08 | 2.95E-06 |
| dd_Smed_v6_20436_0_1 | 44.60 | 1.52 | 0.45 | 3.39 | 6.95E-04 | 6.59E-03 |
| dd_Smed_v6_10685_0_1 | 184.10 | 1.52 | 0.30 | 5.14 | 2.71E-07 | 1.64E-05 |
| dd_Smed_v6_12340_0_1 | 248.05 | 1.52 | 0.29 | 5.28 | 1.30E-07 | 9.25E-06 |
| dd_Smed_v6_20777_0_1 | 100.02 | 1.52 | 0.35 | 4.37 | 1.25E-05 | 3.23E-04 |
| dd_Smed_v6_11926_1_1 | 276.89 | 1.52 | 0.28 | 5.38 | 7.49E-08 | 5.84E-06 |
| dd_Smed_v6_9422_0_1 | 309.52 | 1.51 | 0.25 | 5.93 | 3.09E-09 | 4.00E-07 |
| dd_Smed_v6_12789_0_1 | 1363.17 | 1.51 | 0.25 | 6.02 | 1.74E-09 | 2.40E-07 |
| dd_Smed_v6_14865_0_1 | 347.17 | 1.50 | 0.28 | 5.37 | 7.82E-08 | 6.00E-06 |
| dd_Smed_v6_24349_0_1 | 84.15 | 1.49 | 0.44 | 3.42 | 6.21E-04 | 6.08E-03 |
| dd_Smed_v6_9068_0_1 | 110.40 | 1.49 | 0.38 | 3.93 | 8.37E-05 | 1.36E-03 |
| dd_Smed_v6_43999_0_1 | 69.91 | 1.48 | 0.44 | 3.35 | 8.08E-04 | 7.34E-03 |
| dd_Smed_v6_12521_0_1 | 435.24 | 1.47 | 0.22 | 6.60 | 4.13E-11 | 9.02E-09 |
| dd_Smed_v6_14806_0_1 | 2438.16 | 1.47 | 0.25 | 5.85 | 4.90E-09 | 5.89E-07 |
| dd_Smed_v6_10090_0_1 | 108.72 | 1.47 | 0.31 | 4.74 | 2.09E-06 | 8.16E-05 |
| dd_Smed_v6_12960_0_1 | 647.72 | 1.46 | 0.25 | 5.93 | 3.11E-09 | 4.00E-07 |
| dd_Smed_v6_11412_0_1 | 1754.57 | 1.46 | 0.21 | 7.06 | 1.70E-12 | 6.08E-10 |
| dd_Smed_v6_10976_0_1 | 287.59 | 1.45 | 0.21 | 7.08 | 1.47E-12 | 5.55E-10 |
| dd_Smed_v6_4079_0_1 | 135.67 | 1.45 | 0.42 | 3.44 | 5.85E-04 | 5.82E-03 |
| dd_Smed_v6_12357_0_1 | 471.14 | 1.45 | 0.19 | 7.49 | 6.74E-14 | 4.70E-11 |
| dd_Smed_v6_12959_0_1 | 90.91 | 1.45 | 0.39 | 3.73 | 1.95E-04 | 2.62E-03 |
| dd_Smed_v6_16183_0_1 | 126.22 | 1.45 | 0.36 | 4.00 | 6.21E-05 | 1.08E-03 |
| dd_Smed_v6_14413_0_1 | 93.24 | 1.45 | 0.38 | 3.82 | 1.32E-04 | 1.96E-03 |
| dd_Smed_v6_12013_0_1 | 1837.89 | 1.44 | 0.26 | 5.60 | 2.10E-08 | 2.09E-06 |
| dd_Smed_v6_4005_0_1 | 2773.81 | 1.44 | 0.16 | 8.84 | 9.77E-19 | 2.86E-15 |
| dd_Smed_v6_19710_0_1 | 209.86 | 1.44 | 0.30 | 4.84 | 1.31E-06 | 5.53E-05 |
| dd_Smed_v6_8679_0_1 | 532.87 | 1.43 | 0.20 | 7.20 | 5.99E-13 | 2.51E-10 |
| dd_Smed_v6_6875_0_1 | 984.53 | 1.43 | 0.20 | 7.02 | 2.19E-12 | 7.30E-10 |
| dd_Smed_v6_18952_0_1 | 155.10 | 1.43 | 0.40 | 3.60 | 3.13E-04 | 3.64E-03 |
| dd_Smed_v6_20152_0_1 | 143.70 | 1.42 | 0.30 | 4.73 | 2.24E-06 | 8.64E-05 |
| dd_Smed_v6_10440_0_1 | 520.35 | 1.42 | 0.18 | 7.85 | 4.23E-15 | 5.17E-12 |
| dd_Smed_v6_9160_0_1 | 177.55 | 1.42 | 0.30 | 4.78 | 1.79E-06 | 7.11E-05 |
| dd_Smed_v6_14261_0_1 | 415.32 | 1.42 | 0.34 | 4.16 | 3.22E-05 | 6.49E-04 |
| dd_Smed_v6_9752_0_1 | 197.39 | 1.42 | 0.27 | 5.27 | 1.37E-07 | 9.53E-06 |
| dd_Smed_v6_9923_0_1 | 533.23 | 1.42 | 0.21 | 6.90 | 5.24E-12 | 1.47E-09 |
| dd_Smed_v6_7953_0_1 | 735.20 | 1.41 | 0.18 | 7.67 | 1.74E-14 | 1.50E-11 |
| dd_Smed_v6_6054_0_1 | 92.16 | 1.41 | 0.34 | 4.17 | 3.10E-05 | 6.32E-04 |
| dd_Smed_v6_8052_0_1 | 289.54 | 1.41 | 0.25 | 5.54 | 3.00E-08 | 2.86E-06 |

| | | | | | | |
|---|---|---|---|---|---|---|
| dd_Smed_v6_11624_0_1 | 352.27 | 1.40 | 0.24 | 5.89 | 3.79E-09 | 4.62E-07 |
| dd_Smed_v6_20423_0_1 | 281.16 | 1.40 | 0.24 | 5.84 | 5.21E-09 | 6.16E-07 |
| dd_Smed_v6_13002_0_1 | 102.28 | 1.39 | 0.35 | 3.96 | 7.48E-05 | 1.24E-03 |
| dd_Smed_v6_9897_0_1 | 336.92 | 1.39 | 0.21 | 6.57 | 5.14E-11 | 1.09E-08 |
| dd_Smed_v6_12829_0_1 | 146.51 | 1.38 | 0.30 | 4.58 | 4.59E-06 | 1.51E-04 |
| dd_Smed_v6_6237_0_1 | 768.46 | 1.38 | 0.17 | 8.23 | 1.86E-16 | 4.55E-13 |
| dd_Smed_v6_13879_0_1 | 314.61 | 1.38 | 0.22 | 6.38 | 1.80E-10 | 3.18E-08 |
| dd_Smed_v6_6452_0_1 | 854.02 | 1.38 | 0.20 | 6.83 | 8.68E-12 | 2.31E-09 |
| dd_Smed_v6_40000_0_1 | 42.21 | 1.38 | 0.41 | 3.37 | 7.46E-04 | 6.92E-03 |
| dd_Smed_v6_9774_0_1 | 129.42 | 1.37 | 0.31 | 4.41 | 1.02E-05 | 2.73E-04 |
| dd_Smed_v6_13582_0_1 | 418.87 | 1.37 | 0.33 | 4.21 | 2.56E-05 | 5.48E-04 |
| dd_Smed_v6_6274_0_1 | 371.11 | 1.37 | 0.30 | 4.62 | 3.87E-06 | 1.33E-04 |
| dd_Smed_v6_10725_0_1 | 538.85 | 1.37 | 0.19 | 7.17 | 7.36E-13 | 3.00E-10 |
| dd_Smed_v6_10268_0_1 | 297.09 | 1.37 | 0.27 | 5.16 | 2.40E-07 | 1.50E-05 |
| dd_Smed_v6_12160_0_1 | 87.19 | 1.36 | 0.39 | 3.46 | 5.42E-04 | 5.50E-03 |
| dd_Smed_v6_22739_0_1 | 196.12 | 1.36 | 0.27 | 4.98 | 6.43E-07 | 3.21E-05 |
| dd_Smed_v6_8684_0_1 | 809.23 | 1.35 | 0.17 | 7.89 | 3.05E-15 | 4.79E-12 |
| dd_Smed_v6_5527_0_1 | 557.67 | 1.35 | 0.19 | 6.97 | 3.21E-12 | 1.00E-09 |
| dd_Smed_v6_5623_0_1 | 1167.01 | 1.35 | 0.18 | 7.47 | 8.07E-14 | 5.18E-11 |
| dd_Smed_v6_23589_0_1 | 203.11 | 1.35 | 0.29 | 4.66 | 3.19E-06 | 1.14E-04 |
| dd_Smed_v6_12773_0_1 | 231.80 | 1.34 | 0.37 | 3.58 | 3.46E-04 | 3.95E-03 |
| dd_Smed_v6_11729_0_1 | 1284.89 | 1.34 | 0.23 | 5.79 | 7.11E-09 | 8.20E-07 |
| dd_Smed_v6_15163_0_1 | 184.55 | 1.33 | 0.31 | 4.34 | 1.44E-05 | 3.58E-04 |
| dd_Smed_v6_12599_0_1 | 367.68 | 1.33 | 0.33 | 4.07 | 4.80E-05 | 8.81E-04 |
| dd_Smed_v6_9007_0_1 | 776.81 | 1.32 | 0.26 | 5.03 | 4.96E-07 | 2.61E-05 |
| dd_Smed_v6_12221_0_1 | 384.20 | 1.32 | 0.22 | 6.03 | 1.65E-09 | 2.35E-07 |
| dd_Smed_v6_30487_0_1 | 69.78 | 1.32 | 0.40 | 3.32 | 9.03E-04 | 7.97E-03 |
| dd_Smed_v6_6443_0_1 | 704.82 | 1.32 | 0.17 | 7.64 | 2.18E-14 | 1.77E-11 |
| dd_Smed_v6_8614_0_1 | 585.35 | 1.32 | 0.18 | 7.27 | 3.48E-13 | 1.58E-10 |
| dd_Smed_v6_15678_0_1 | 293.80 | 1.32 | 0.26 | 5.13 | 2.86E-07 | 1.70E-05 |
| dd_Smed_v6_15012_0_1 | 307.37 | 1.32 | 0.29 | 4.47 | 7.98E-06 | 2.30E-04 |
| dd_Smed_v6_11087_0_1 | 640.64 | 1.31 | 0.23 | 5.78 | 7.60E-09 | 8.63E-07 |
| dd_Smed_v6_10514_0_1 | 664.90 | 1.31 | 0.32 | 4.09 | 4.38E-05 | 8.22E-04 |
| dd_Smed_v6_12126_1_1 | 126.36 | 1.31 | 0.37 | 3.52 | 4.29E-04 | 4.60E-03 |
| dd_Smed_v6_15720_0_1 | 87.44 | 1.31 | 0.32 | 4.09 | 4.32E-05 | 8.14E-04 |
| dd_Smed_v6_10885_0_1 | 158.96 | 1.31 | 0.33 | 3.95 | 7.95E-05 | 1.30E-03 |
| dd_Smed_v6_7968_0_1 | 880.24 | 1.31 | 0.29 | 4.58 | 4.58E-06 | 1.51E-04 |
| dd_Smed_v6_10019_0_1 | 866.71 | 1.31 | 0.31 | 4.24 | 2.21E-05 | 4.89E-04 |
| dd_Smed_v6_9795_0_1 | 468.37 | 1.31 | 0.26 | 4.99 | 6.01E-07 | 3.05E-05 |
| dd_Smed_v6_5511_0_1 | 1024.31 | 1.31 | 0.19 | 6.74 | 1.60E-11 | 3.97E-09 |
| dd_Smed_v6_3018_0_1 | 1248.70 | 1.31 | 0.27 | 4.91 | 9.30E-07 | 4.27E-05 |
| dd_Smed_v6_6078_0_1 | 660.15 | 1.30 | 0.17 | 7.74 | 9.88E-15 | 9.65E-12 |
| dd_Smed_v6_5471_0_1 | 426.75 | 1.30 | 0.30 | 4.30 | 1.74E-05 | 4.19E-04 |

| | | | | | | |
|---|---|---|---|---|---|---|
| dd_Smed_v6_4731_0_1 | 1201.28 | 1.30 | 0.21 | 6.10 | 1.08E-09 | 1.64E-07 |
| dd_Smed_v6_8445_0_1 | 142.29 | 1.30 | 0.35 | 3.69 | 2.22E-04 | 2.85E-03 |
| dd_Smed_v6_11158_0_1 | 297.75 | 1.30 | 0.34 | 3.80 | 1.46E-04 | 2.11E-03 |
| dd_Smed_v6_12665_0_1 | 304.49 | 1.30 | 0.31 | 4.22 | 2.40E-05 | 5.19E-04 |
| dd_Smed_v6_10881_0_1 | 487.67 | 1.30 | 0.35 | 3.67 | 2.40E-04 | 3.01E-03 |
| dd_Smed_v6_3402_0_1 | 79.84 | 1.30 | 0.37 | 3.46 | 5.30E-04 | 5.42E-03 |
| dd_Smed_v6_1420_0_1 | 379.05 | 1.29 | 0.28 | 4.58 | 4.63E-06 | 1.51E-04 |
| dd_Smed_v6_16851_0_1 | 359.99 | 1.29 | 0.26 | 5.02 | 5.18E-07 | 2.70E-05 |
| dd_Smed_v6_11325_0_1 | 193.29 | 1.29 | 0.34 | 3.81 | 1.38E-04 | 2.02E-03 |
| dd_Smed_v6_13168_0_1 | 112.93 | 1.29 | 0.35 | 3.67 | 2.43E-04 | 3.03E-03 |
| dd_Smed_v6_15237_0_1 | 395.25 | 1.29 | 0.25 | 5.21 | 1.86E-07 | 1.22E-05 |
| dd_Smed_v6_14158_0_1 | 220.65 | 1.28 | 0.35 | 3.64 | 2.73E-04 | 3.29E-03 |
| dd_Smed_v6_5058_0_1 | 4402.86 | 1.28 | 0.18 | 7.33 | 2.36E-13 | 1.15E-10 |
| dd_Smed_v6_18287_0_1 | 133.62 | 1.28 | 0.36 | 3.53 | 4.23E-04 | 4.56E-03 |
| dd_Smed_v6_7786_0_1 | 228.78 | 1.27 | 0.29 | 4.40 | 1.11E-05 | 2.90E-04 |
| dd_Smed_v6_5194_0_1 | 452.27 | 1.27 | 0.17 | 7.27 | 3.56E-13 | 1.58E-10 |
| dd_Smed_v6_6349_0_1 | 518.97 | 1.27 | 0.24 | 5.21 | 1.88E-07 | 1.23E-05 |
| dd_Smed_v6_15655_0_1 | 201.06 | 1.26 | 0.39 | 3.24 | 1.18E-03 | 9.72E-03 |
| dd_Smed_v6_7703_0_1 | 928.66 | 1.26 | 0.23 | 5.53 | 3.18E-08 | 2.95E-06 |
| dd_Smed_v6_3417_0_1 | 617.96 | 1.25 | 0.21 | 5.85 | 5.05E-09 | 6.02E-07 |
| dd_Smed_v6_2971_0_1 | 2713.38 | 1.25 | 0.16 | 7.87 | 3.49E-15 | 4.79E-12 |
| dd_Smed_v6_20092_0_1 | 114.44 | 1.25 | 0.37 | 3.36 | 7.80E-04 | 7.14E-03 |
| dd_Smed_v6_21373_0_1 | 128.69 | 1.25 | 0.37 | 3.40 | 6.72E-04 | 6.42E-03 |
| dd_Smed_v6_14167_0_1 | 281.91 | 1.25 | 0.23 | 5.42 | 5.92E-08 | 4.88E-06 |
| dd_Smed_v6_3836_0_1 | 1503.00 | 1.24 | 0.21 | 6.05 | 1.47E-09 | 2.12E-07 |
| dd_Smed_v6_6263_0_1 | 1895.00 | 1.24 | 0.18 | 7.08 | 1.48E-12 | 5.55E-10 |
| dd_Smed_v6_18130_0_1 | 532.82 | 1.24 | 0.32 | 3.92 | 8.88E-05 | 1.43E-03 |
| dd_Smed_v6_10753_0_1 | 304.17 | 1.24 | 0.22 | 5.59 | 2.30E-08 | 2.26E-06 |
| dd_Smed_v6_7576_0_1 | 857.69 | 1.24 | 0.19 | 6.49 | 8.74E-11 | 1.80E-08 |
| dd_Smed_v6_8307_0_1 | 277.47 | 1.24 | 0.29 | 4.28 | 1.87E-05 | 4.36E-04 |
| dd_Smed_v6_9370_0_1 | 273.38 | 1.23 | 0.19 | 6.57 | 4.89E-11 | 1.05E-08 |
| dd_Smed_v6_14758_0_1 | 209.25 | 1.23 | 0.34 | 3.67 | 2.44E-04 | 3.04E-03 |
| dd_Smed_v6_30278_0_1 | 190.18 | 1.23 | 0.28 | 4.44 | 8.90E-06 | 2.47E-04 |
| dd_Smed_v6_45908_0_1 | 190.17 | 1.23 | 0.35 | 3.53 | 4.23E-04 | 4.56E-03 |
| dd_Smed_v6_6965_0_1 | 174.23 | 1.23 | 0.37 | 3.36 | 7.93E-04 | 7.22E-03 |
| dd_Smed_v6_6741_0_1 | 838.78 | 1.23 | 0.18 | 6.75 | 1.48E-11 | 3.75E-09 |
| dd_Smed_v6_8887_0_1 | 540.45 | 1.23 | 0.24 | 5.03 | 4.90E-07 | 2.60E-05 |
| dd_Smed_v6_9445_0_1 | 755.68 | 1.23 | 0.18 | 6.93 | 4.25E-12 | 1.26E-09 |
| dd_Smed_v6_8001_0_1 | 2830.39 | 1.23 | 0.20 | 6.15 | 7.86E-10 | 1.21E-07 |
| dd_Smed_v6_8570_0_1 | 1458.09 | 1.22 | 0.17 | 7.38 | 1.53E-13 | 8.55E-11 |
| dd_Smed_v6_16306_0_1 | 138.70 | 1.22 | 0.29 | 4.17 | 3.01E-05 | 6.17E-04 |
| dd_Smed_v6_9527_0_1 | 300.57 | 1.22 | 0.26 | 4.66 | 3.09E-06 | 1.12E-04 |
| dd_Smed_v6_14097_0_1 | 85.22 | 1.22 | 0.36 | 3.34 | 8.23E-04 | 7.46E-03 |

| | | | | | | |
|---|---|---|---|---|---|---|
| dd_Smed_v6_9454_0_1 | 879.21 | 1.21 | 0.20 | 6.12 | 9.08E-10 | 1.39E-07 |
| dd_Smed_v6_13402_0_1 | 521.39 | 1.21 | 0.32 | 3.76 | 1.69E-04 | 2.34E-03 |
| dd_Smed_v6_6013_0_1 | 162.66 | 1.21 | 0.36 | 3.41 | 6.53E-04 | 6.29E-03 |
| dd_Smed_v6_11450_0_1 | 128.31 | 1.21 | 0.35 | 3.41 | 6.43E-04 | 6.23E-03 |
| dd_Smed_v6_9681_0_1 | 468.11 | 1.21 | 0.23 | 5.16 | 2.49E-07 | 1.54E-05 |
| dd_Smed_v6_7650_0_1 | 390.55 | 1.20 | 0.24 | 5.00 | 5.80E-07 | 2.96E-05 |
| dd_Smed_v6_22244_0_1 | 192.27 | 1.20 | 0.28 | 4.29 | 1.75E-05 | 4.20E-04 |
| dd_Smed_v6_11013_0_1 | 230.75 | 1.20 | 0.29 | 4.11 | 3.95E-05 | 7.63E-04 |
| dd_Smed_v6_11682_0_1 | 180.89 | 1.19 | 0.29 | 4.11 | 4.00E-05 | 7.68E-04 |
| dd_Smed_v6_15397_0_1 | 417.58 | 1.19 | 0.28 | 4.18 | 2.96E-05 | 6.10E-04 |
| dd_Smed_v6_1582_0_1 | 11035.79 | 1.19 | 0.18 | 6.73 | 1.70E-11 | 4.13E-09 |
| dd_Smed_v6_7274_0_1 | 468.06 | 1.18 | 0.17 | 7.02 | 2.15E-12 | 7.30E-10 |
| dd_Smed_v6_10376_0_1 | 488.26 | 1.18 | 0.30 | 3.95 | 7.95E-05 | 1.30E-03 |
| dd_Smed_v6_4878_0_1 | 6555.55 | 1.18 | 0.21 | 5.53 | 3.20E-08 | 2.95E-06 |
| dd_Smed_v6_10370_0_1 | 266.65 | 1.18 | 0.23 | 5.06 | 4.29E-07 | 2.33E-05 |
| dd_Smed_v6_9004_0_1 | 325.18 | 1.18 | 0.22 | 5.28 | 1.31E-07 | 9.25E-06 |
| dd_Smed_v6_20961_0_1 | 98.48 | 1.18 | 0.33 | 3.62 | 3.00E-04 | 3.52E-03 |
| dd_Smed_v6_20433_0_1 | 156.15 | 1.18 | 0.28 | 4.14 | 3.43E-05 | 6.81E-04 |
| dd_Smed_v6_13381_0_1 | 512.83 | 1.18 | 0.25 | 4.79 | 1.69E-06 | 6.81E-05 |
| dd_Smed_v6_6268_0_1 | 869.20 | 1.18 | 0.18 | 6.41 | 1.46E-10 | 2.71E-08 |
| dd_Smed_v6_15866_0_1 | 172.73 | 1.17 | 0.31 | 3.76 | 1.70E-04 | 2.35E-03 |
| dd_Smed_v6_21626_0_1 | 125.86 | 1.17 | 0.35 | 3.32 | 8.88E-04 | 7.90E-03 |
| dd_Smed_v6_7405_0_1 | 2064.06 | 1.17 | 0.22 | 5.31 | 1.10E-07 | 8.05E-06 |
| dd_Smed_v6_10711_0_1 | 335.41 | 1.17 | 0.27 | 4.27 | 1.97E-05 | 4.51E-04 |
| dd_Smed_v6_4419_0_1 | 550.08 | 1.17 | 0.20 | 5.93 | 3.10E-09 | 4.00E-07 |
| dd_Smed_v6_5681_0_1 | 885.01 | 1.16 | 0.22 | 5.23 | 1.73E-07 | 1.15E-05 |
| dd_Smed_v6_25001_0_1 | 91.46 | 1.16 | 0.34 | 3.43 | 6.12E-04 | 6.02E-03 |
| dd_Smed_v6_12491_0_1 | 296.09 | 1.16 | 0.20 | 5.72 | 1.08E-08 | 1.14E-06 |
| dd_Smed_v6_7132_0_1 | 209.96 | 1.16 | 0.31 | 3.72 | 1.97E-04 | 2.64E-03 |
| dd_Smed_v6_12518_0_1 | 191.24 | 1.16 | 0.25 | 4.57 | 4.98E-06 | 1.58E-04 |
| dd_Smed_v6_8212_0_1 | 318.96 | 1.16 | 0.35 | 3.27 | 1.09E-03 | 9.22E-03 |
| dd_Smed_v6_4082_0_1 | 462.34 | 1.16 | 0.26 | 4.39 | 1.14E-05 | 2.98E-04 |
| dd_Smed_v6_2933_0_1 | 672.09 | 1.15 | 0.16 | 7.13 | 1.01E-12 | 3.99E-10 |
| dd_Smed_v6_13140_0_1 | 441.47 | 1.15 | 0.21 | 5.58 | 2.42E-08 | 2.36E-06 |
| dd_Smed_v6_9815_0_1 | 1999.87 | 1.15 | 0.23 | 4.99 | 5.95E-07 | 3.03E-05 |
| dd_Smed_v6_20991_0_1 | 114.64 | 1.15 | 0.35 | 3.26 | 1.10E-03 | 9.24E-03 |
| dd_Smed_v6_4912_0_1 | 704.17 | 1.15 | 0.18 | 6.48 | 9.07E-11 | 1.85E-08 |
| dd_Smed_v6_7988_0_1 | 1100.73 | 1.14 | 0.23 | 4.89 | 9.99E-07 | 4.49E-05 |
| dd_Smed_v6_6828_0_1 | 861.07 | 1.14 | 0.23 | 5.01 | 5.56E-07 | 2.87E-05 |
| dd_Smed_v6_14296_0_1 | 200.57 | 1.14 | 0.25 | 4.52 | 6.05E-06 | 1.85E-04 |
| dd_Smed_v6_12381_0_1 | 669.88 | 1.13 | 0.27 | 4.23 | 2.38E-05 | 5.15E-04 |
| dd_Smed_v6_3562_0_1 | 73.47 | 1.13 | 0.31 | 3.67 | 2.44E-04 | 3.04E-03 |
| dd_Smed_v6_2866_0_1 | 2071.77 | 1.13 | 0.16 | 7.06 | 1.63E-12 | 5.96E-10 |

| | | | | | | |
|---|---|---|---|---|---|---|
| dd_Smed_v6_8047_0_1 | 116.56 | 1.13 | 0.29 | 3.83 | 1.26E-04 | 1.90E-03 |
| dd_Smed_v6_21762_0_1 | 269.99 | 1.13 | 0.32 | 3.48 | 5.08E-04 | 5.26E-03 |
| dd_Smed_v6_6632_0_1 | 79.45 | 1.13 | 0.35 | 3.26 | 1.10E-03 | 9.24E-03 |
| dd_Smed_v6_7834_0_1 | 495.89 | 1.13 | 0.23 | 5.00 | 5.63E-07 | 2.89E-05 |
| dd_Smed_v6_5436_0_1 | 654.68 | 1.13 | 0.22 | 5.20 | 2.04E-07 | 1.30E-05 |
| dd_Smed_v6_22620_0_1 | 136.99 | 1.12 | 0.33 | 3.45 | 5.53E-04 | 5.57E-03 |
| dd_Smed_v6_14271_0_1 | 484.03 | 1.12 | 0.30 | 3.80 | 1.43E-04 | 2.09E-03 |
| dd_Smed_v6_18176_0_1 | 149.88 | 1.12 | 0.33 | 3.41 | 6.55E-04 | 6.29E-03 |
| dd_Smed_v6_6313_0_1 | 510.94 | 1.12 | 0.21 | 5.36 | 8.25E-08 | 6.29E-06 |
| dd_Smed_v6_6628_0_1 | 259.01 | 1.12 | 0.28 | 3.99 | 6.59E-05 | 1.13E-03 |
| dd_Smed_v6_14050_0_1 | 436.63 | 1.12 | 0.22 | 5.12 | 3.08E-07 | 1.80E-05 |
| dd_Smed_v6_16050_0_1 | 285.54 | 1.12 | 0.22 | 5.14 | 2.75E-07 | 1.65E-05 |
| dd_Smed_v6_15527_0_1 | 206.65 | 1.12 | 0.23 | 4.79 | 1.66E-06 | 6.69E-05 |
| dd_Smed_v6_11164_0_1 | 270.98 | 1.12 | 0.21 | 5.41 | 6.29E-08 | 5.04E-06 |
| dd_Smed_v6_17704_0_1 | 803.73 | 1.11 | 0.26 | 4.28 | 1.85E-05 | 4.35E-04 |
| dd_Smed_v6_11308_0_1 | 342.14 | 1.11 | 0.22 | 5.15 | 2.60E-07 | 1.59E-05 |
| dd_Smed_v6_4407_0_1 | 903.02 | 1.11 | 0.20 | 5.47 | 4.41E-08 | 3.87E-06 |
| dd_Smed_v6_10699_0_1 | 557.81 | 1.11 | 0.25 | 4.40 | 1.09E-05 | 2.87E-04 |
| dd_Smed_v6_7307_0_1 | 553.29 | 1.11 | 0.19 | 5.83 | 5.70E-09 | 6.68E-07 |
| dd_Smed_v6_8750_0_1 | 1184.78 | 1.11 | 0.19 | 5.75 | 9.07E-09 | 9.91E-07 |
| dd_Smed_v6_5029_0_1 | 207.49 | 1.11 | 0.29 | 3.79 | 1.49E-04 | 2.14E-03 |
| dd_Smed_v6_9852_0_1 | 248.29 | 1.10 | 0.26 | 4.27 | 2.00E-05 | 4.53E-04 |
| dd_Smed_v6_9132_0_1 | 1069.30 | 1.10 | 0.16 | 6.75 | 1.47E-11 | 3.75E-09 |
| dd_Smed_v6_9940_0_1 | 183.66 | 1.10 | 0.29 | 3.75 | 1.74E-04 | 2.40E-03 |
| dd_Smed_v6_9333_0_1 | 334.48 | 1.10 | 0.23 | 4.83 | 1.35E-06 | 5.65E-05 |
| dd_Smed_v6_11739_0_1 | 244.46 | 1.10 | 0.29 | 3.86 | 1.14E-04 | 1.76E-03 |
| dd_Smed_v6_2966_0_1 | 2287.50 | 1.09 | 0.19 | 5.90 | 3.61E-09 | 4.52E-07 |
| dd_Smed_v6_4840_0_1 | 808.48 | 1.09 | 0.16 | 7.04 | 1.88E-12 | 6.55E-10 |
| dd_Smed_v6_9931_0_1 | 229.05 | 1.09 | 0.24 | 4.60 | 4.13E-06 | 1.38E-04 |
| dd_Smed_v6_24476_0_1 | 117.08 | 1.09 | 0.31 | 3.47 | 5.17E-04 | 5.33E-03 |
| dd_Smed_v6_12197_0_1 | 408.90 | 1.09 | 0.24 | 4.51 | 6.57E-06 | 1.98E-04 |
| dd_Smed_v6_7018_0_1 | 583.77 | 1.09 | 0.24 | 4.48 | 7.49E-06 | 2.19E-04 |
| dd_Smed_v6_9126_0_1 | 187.50 | 1.09 | 0.32 | 3.36 | 7.83E-04 | 7.16E-03 |
| dd_Smed_v6_9029_0_1 | 159.48 | 1.09 | 0.30 | 3.57 | 3.61E-04 | 4.06E-03 |
| dd_Smed_v6_9017_0_1 | 1463.71 | 1.08 | 0.18 | 5.97 | 2.41E-09 | 3.24E-07 |
| dd_Smed_v6_5633_0_1 | 560.45 | 1.08 | 0.19 | 5.78 | 7.67E-09 | 8.64E-07 |
| dd_Smed_v6_18318_0_1 | 163.71 | 1.08 | 0.27 | 3.99 | 6.49E-05 | 1.12E-03 |
| dd_Smed_v6_10404_0_1 | 1110.76 | 1.08 | 0.23 | 4.67 | 2.95E-06 | 1.08E-04 |
| dd_Smed_v6_4991_0_1 | 558.36 | 1.08 | 0.22 | 4.91 | 9.13E-07 | 4.22E-05 |
| dd_Smed_v6_9586_0_1 | 785.83 | 1.08 | 0.23 | 4.70 | 2.56E-06 | 9.53E-05 |
| dd_Smed_v6_1558_0_1 | 395.42 | 1.08 | 0.24 | 4.48 | 7.41E-06 | 2.18E-04 |
| dd_Smed_v6_13224_0_1 | 388.37 | 1.08 | 0.22 | 4.90 | 9.54E-07 | 4.36E-05 |
| dd_Smed_v6_3746_0_1 | 339.08 | 1.08 | 0.26 | 4.10 | 4.21E-05 | 7.99E-04 |

| | | | | | | |
|---|---|---|---|---|---|---|
| dd_Smed_v6_10525_0_1 | 397.25 | 1.08 | 0.27 | 4.07 | 4.78E-05 | 8.80E-04 |
| dd_Smed_v6_12187_0_1 | 937.67 | 1.08 | 0.15 | 7.38 | 1.57E-13 | 8.55E-11 |
| dd_Smed_v6_5659_0_1 | 953.92 | 1.08 | 0.21 | 5.25 | 1.51E-07 | 1.03E-05 |
| dd_Smed_v6_9516_0_1 | 725.61 | 1.08 | 0.22 | 4.89 | 1.01E-06 | 4.52E-05 |
| dd_Smed_v6_11759_0_1 | 488.18 | 1.07 | 0.24 | 4.43 | 9.37E-06 | 2.56E-04 |
| dd_Smed_v6_6715_0_1 | 1055.35 | 1.07 | 0.20 | 5.42 | 5.96E-08 | 4.88E-06 |
| dd_Smed_v6_11920_0_1 | 1012.06 | 1.07 | 0.21 | 5.11 | 3.29E-07 | 1.89E-05 |
| dd_Smed_v6_5422_0_1 | 287.62 | 1.07 | 0.20 | 5.26 | 1.44E-07 | 9.94E-06 |
| dd_Smed_v6_9469_0_1 | 609.57 | 1.07 | 0.23 | 4.63 | 3.65E-06 | 1.27E-04 |
| dd_Smed_v6_8119_0_1 | 290.66 | 1.06 | 0.20 | 5.30 | 1.18E-07 | 8.55E-06 |
| dd_Smed_v6_4353_0_1 | 1096.65 | 1.06 | 0.21 | 5.17 | 2.33E-07 | 1.46E-05 |
| dd_Smed_v6_17265_0_1 | 205.22 | 1.06 | 0.26 | 4.14 | 3.46E-05 | 6.84E-04 |
| dd_Smed_v6_5544_0_1 | 761.30 | 1.06 | 0.20 | 5.26 | 1.45E-07 | 9.98E-06 |
| dd_Smed_v6_4279_0_1 | 428.54 | 1.06 | 0.24 | 4.45 | 8.77E-06 | 2.45E-04 |
| dd_Smed_v6_8469_0_1 | 1722.16 | 1.06 | 0.25 | 4.29 | 1.78E-05 | 4.24E-04 |
| dd_Smed_v6_6821_0_1 | 507.92 | 1.06 | 0.26 | 4.08 | 4.41E-05 | 8.25E-04 |
| dd_Smed_v6_6097_0_1 | 254.98 | 1.06 | 0.24 | 4.38 | 1.17E-05 | 3.04E-04 |
| dd_Smed_v6_4723_0_1 | 634.29 | 1.06 | 0.23 | 4.66 | 3.16E-06 | 1.13E-04 |
| dd_Smed_v6_12446_0_1 | 216.95 | 1.05 | 0.24 | 4.46 | 8.24E-06 | 2.35E-04 |
| dd_Smed_v6_17552_0_1 | 155.63 | 1.05 | 0.29 | 3.64 | 2.74E-04 | 3.29E-03 |
| dd_Smed_v6_5513_0_1 | 657.65 | 1.05 | 0.25 | 4.20 | 2.70E-05 | 5.73E-04 |
| dd_Smed_v6_11860_0_1 | 695.17 | 1.04 | 0.24 | 4.43 | 9.28E-06 | 2.54E-04 |
| dd_Smed_v6_5671_0_1 | 343.65 | 1.04 | 0.28 | 3.74 | 1.83E-04 | 2.49E-03 |
| dd_Smed_v6_11263_0_1 | 1738.99 | 1.04 | 0.25 | 4.17 | 2.99E-05 | 6.15E-04 |
| dd_Smed_v6_12333_0_1 | 647.90 | 1.04 | 0.25 | 4.19 | 2.83E-05 | 5.91E-04 |
| dd_Smed_v6_5573_0_1 | 1701.68 | 1.04 | 0.19 | 5.53 | 3.12E-08 | 2.95E-06 |
| dd_Smed_v6_4010_0_1 | 1607.96 | 1.04 | 0.15 | 6.73 | 1.72E-11 | 4.13E-09 |
| dd_Smed_v6_8162_0_1 | 232.95 | 1.04 | 0.22 | 4.62 | 3.77E-06 | 1.29E-04 |
| dd_Smed_v6_10034_0_1 | 1263.66 | 1.04 | 0.22 | 4.70 | 2.64E-06 | 9.76E-05 |
| dd_Smed_v6_2740_0_1 | 2265.33 | 1.03 | 0.21 | 4.90 | 9.74E-07 | 4.42E-05 |
| dd_Smed_v6_10335_0_1 | 2345.28 | 1.03 | 0.22 | 4.64 | 3.42E-06 | 1.20E-04 |
| dd_Smed_v6_8874_0_1 | 344.97 | 1.03 | 0.25 | 4.11 | 3.99E-05 | 7.67E-04 |
| dd_Smed_v6_8241_0_1 | 275.65 | 1.03 | 0.28 | 3.72 | 1.98E-04 | 2.64E-03 |
| dd_Smed_v6_13727_0_1 | 1086.54 | 1.03 | 0.23 | 4.54 | 5.51E-06 | 1.72E-04 |
| dd_Smed_v6_14977_0_1 | 182.55 | 1.03 | 0.26 | 3.96 | 7.65E-05 | 1.26E-03 |
| dd_Smed_v6_8658_0_1 | 285.01 | 1.03 | 0.22 | 4.75 | 2.02E-06 | 7.92E-05 |
| dd_Smed_v6_11558_0_1 | 1177.37 | 1.03 | 0.24 | 4.21 | 2.55E-05 | 5.46E-04 |
| dd_Smed_v6_8654_0_1 | 701.75 | 1.03 | 0.20 | 5.08 | 3.83E-07 | 2.13E-05 |
| dd_Smed_v6_11279_0_1 | 717.48 | 1.02 | 0.19 | 5.34 | 9.39E-08 | 7.02E-06 |
| dd_Smed_v6_12962_0_1 | 565.57 | 1.02 | 0.26 | 3.98 | 6.95E-05 | 1.18E-03 |
| dd_Smed_v6_17731_0_1 | 331.03 | 1.02 | 0.26 | 3.97 | 7.31E-05 | 1.22E-03 |
| dd_Smed_v6_12568_0_1 | 292.41 | 1.02 | 0.27 | 3.78 | 1.59E-04 | 2.24E-03 |
| dd_Smed_v6_5183_0_1 | 3549.62 | 1.02 | 0.16 | 6.41 | 1.45E-10 | 2.71E-08 |

| | | | | | | |
|---|---|---|---|---|---|---|
| dd_Smed_v6_10439_0_1 | 117.10 | 1.02 | 0.27 | 3.78 | 1.57E-04 | 2.22E-03 |
| dd_Smed_v6_6643_0_1 | 409.73 | 1.02 | 0.22 | 4.58 | 4.66E-06 | 1.51E-04 |
| dd_Smed_v6_3953_0_1 | 1521.81 | 1.02 | 0.25 | 4.07 | 4.74E-05 | 8.74E-04 |
| dd_Smed_v6_6567_0_1 | 1283.89 | 1.02 | 0.22 | 4.66 | 3.22E-06 | 1.15E-04 |
| dd_Smed_v6_22716_0_1 | 158.15 | 1.02 | 0.27 | 3.81 | 1.40E-04 | 2.05E-03 |
| dd_Smed_v6_6291_0_1 | 108.28 | 1.02 | 0.31 | 3.24 | 1.21E-03 | 9.91E-03 |
| dd_Smed_v6_6270_0_1 | 874.36 | 1.01 | 0.21 | 4.80 | 1.56E-06 | 6.37E-05 |
| dd_Smed_v6_5465_0_1 | 1769.86 | 1.01 | 0.15 | 6.65 | 2.93E-11 | 6.50E-09 |
| dd_Smed_v6_12403_0_1 | 291.83 | 1.01 | 0.25 | 4.07 | 4.79E-05 | 8.81E-04 |
| dd_Smed_v6_12195_0_1 | 298.79 | 1.01 | 0.30 | 3.40 | 6.79E-04 | 6.47E-03 |
| dd_Smed_v6_12310_0_1 | 822.37 | 1.01 | 0.23 | 4.47 | 7.84E-06 | 2.26E-04 |
| dd_Smed_v6_5197_0_1 | 1215.64 | 1.00 | 0.18 | 5.73 | 1.03E-08 | 1.10E-06 |
| dd_Smed_v6_9001_0_1 | 445.64 | 1.00 | 0.23 | 4.30 | 1.71E-05 | 4.15E-04 |
| dd_Smed_v6_11458_0_1 | 728.80 | 1.00 | 0.20 | 4.97 | 6.61E-07 | 3.27E-05 |

**Table S2: List of downregulated genes in irradiated stem cells (1dpi), atm(RNAi) vs. control(RNAi).**
ID: Planmine transcript ID from dd_Smed_v6 transcriptome.
baseMean: mean of normalized counts for all samples;
log2FC: log2 fold change;
lfcSE: standard error of log2 fold change;
stat: Wald statistic;
pvalue: Wald test p-value;
padj: BH adjusted p-values

Only genes with log2FC <1 and adjusted p-value <0.01 are listed here.

| ID | baseMean | log2FC | lfcSE | stat | pvalue | padj |
|---|---|---|---|---|---|---|
| dd_Smed_v6_8471_0_1 | 14.74 | -3.82 | 1.07 | -3.57 | 3.54E-04 | 4.00E-03 |
| dd_Smed_v6_6494_0_1 | 30.62 | -3.55 | 1.05 | -3.40 | 6.77E-04 | 6.45E-03 |
| dd_Smed_v6_4103_0_1 | 20.46 | -3.52 | 1.02 | -3.44 | 5.77E-04 | 5.76E-03 |
| dd_Smed_v6_2843_0_1 | 29.06 | -3.41 | 0.81 | -4.21 | 2.51E-05 | 5.39E-04 |
| dd_Smed_v6_9220_0_1 | 29.08 | -3.37 | 0.87 | -3.90 | 9.76E-05 | 1.55E-03 |
| dd_Smed_v6_6151_0_1 | 27.78 | -3.16 | 0.84 | -3.77 | 1.62E-04 | 2.27E-03 |
| dd_Smed_v6_5528_0_1 | 20.13 | -3.15 | 0.75 | -4.20 | 2.70E-05 | 5.72E-04 |
| dd_Smed_v6_4104_0_1 | 30.33 | -3.09 | 0.82 | -3.79 | 1.53E-04 | 2.19E-03 |
| dd_Smed_v6_10774_0_1 | 40.86 | -3.03 | 0.84 | -3.61 | 3.06E-04 | 3.58E-03 |
| dd_Smed_v6_2690_0_1 | 46.72 | -2.95 | 0.63 | -4.69 | 2.72E-06 | 9.97E-05 |
| dd_Smed_v6_9917_0_1 | 21.63 | -2.90 | 0.90 | -3.24 | 1.21E-03 | 9.91E-03 |
| dd_Smed_v6_3508_0_1 | 51.09 | -2.88 | 0.62 | -4.66 | 3.14E-06 | 1.13E-04 |
| dd_Smed_v6_6024_0_1 | 22.01 | -2.85 | 0.78 | -3.66 | 2.55E-04 | 3.14E-03 |
| dd_Smed_v6_3065_0_1 | 278.13 | -2.80 | 0.51 | -5.45 | 4.93E-08 | 4.26E-06 |
| dd_Smed_v6_7565_0_1 | 48.81 | -2.68 | 0.78 | -3.45 | 5.56E-04 | 5.59E-03 |
| dd_Smed_v6_3873_0_1 | 37.22 | -2.65 | 0.66 | -4.03 | 5.56E-05 | 9.89E-04 |
| dd_Smed_v6_11520_0_1 | 20.81 | -2.61 | 0.80 | -3.28 | 1.04E-03 | 8.88E-03 |
| dd_Smed_v6_4370_0_1 | 28.98 | -2.61 | 0.75 | -3.46 | 5.35E-04 | 5.45E-03 |
| dd_Smed_v6_617_0_1 | 150.66 | -2.60 | 0.68 | -3.79 | 1.49E-04 | 2.14E-03 |
| dd_Smed_v6_2097_0_1 | 35.63 | -2.58 | 0.62 | -4.18 | 2.92E-05 | 6.05E-04 |
| dd_Smed_v6_1240_0_1 | 29.89 | -2.58 | 0.60 | -4.27 | 1.93E-05 | 4.45E-04 |
| dd_Smed_v6_14638_0_1 | 76.04 | -2.56 | 0.57 | -4.54 | 5.75E-06 | 1.78E-04 |
| dd_Smed_v6_9626_0_1 | 18.54 | -2.54 | 0.72 | -3.53 | 4.09E-04 | 4.44E-03 |
| dd_Smed_v6_35044_0_1 | 58.36 | -2.53 | 0.60 | -4.21 | 2.50E-05 | 5.38E-04 |
| dd_Smed_v6_13538_0_1 | 54.60 | -2.51 | 0.60 | -4.19 | 2.79E-05 | 5.86E-04 |
| dd_Smed_v6_1406_0_1 | 34.10 | -2.51 | 0.67 | -3.77 | 1.61E-04 | 2.27E-03 |
| dd_Smed_v6_9723_0_1 | 33.28 | -2.51 | 0.67 | -3.75 | 1.77E-04 | 2.43E-03 |
| dd_Smed_v6_4876_0_1 | 40.46 | -2.51 | 0.71 | -3.52 | 4.31E-04 | 4.61E-03 |
| dd_Smed_v6_10569_0_1 | 34.18 | -2.51 | 0.77 | -3.26 | 1.11E-03 | 9.32E-03 |
| dd_Smed_v6_22503_0_1 | 37.45 | -2.44 | 0.71 | -3.45 | 5.69E-04 | 5.69E-03 |
| dd_Smed_v6_4668_0_1 | 114.31 | -2.44 | 0.55 | -4.44 | 8.91E-06 | 2.47E-04 |

| | | | | | | |
|---|---|---|---|---|---|---|
| dd_Smed_v6_13707_0_1 | 43.34 | -2.44 | 0.73 | -3.32 | 8.93E-04 | 7.92E-03 |
| dd_Smed_v6_2668_0_1 | 26.60 | -2.43 | 0.69 | -3.50 | 4.70E-04 | 4.95E-03 |
| dd_Smed_v6_10048_0_1 | 32.97 | -2.42 | 0.51 | -4.72 | 2.39E-06 | 9.06E-05 |
| dd_Smed_v6_1009_1_1 | 43.57 | -2.40 | 0.71 | -3.37 | 7.58E-04 | 7.01E-03 |
| dd_Smed_v6_16722_0_1 | 25.28 | -2.39 | 0.69 | -3.47 | 5.28E-04 | 5.41E-03 |
| dd_Smed_v6_797_0_1 | 135.79 | -2.38 | 0.57 | -4.16 | 3.14E-05 | 6.36E-04 |
| dd_Smed_v6_20782_0_1 | 126.82 | -2.37 | 0.70 | -3.38 | 7.36E-04 | 6.86E-03 |
| dd_Smed_v6_6609_0_1 | 62.66 | -2.37 | 0.47 | -5.10 | 3.40E-07 | 1.93E-05 |
| dd_Smed_v6_55569_0_1 | 21.10 | -2.36 | 0.71 | -3.32 | 9.02E-04 | 7.96E-03 |
| dd_Smed_v6_16616_0_1 | 91.64 | -2.34 | 0.61 | -3.86 | 1.15E-04 | 1.77E-03 |
| dd_Smed_v6_87_0_1 | 43.56 | -2.33 | 0.53 | -4.40 | 1.07E-05 | 2.83E-04 |
| dd_Smed_v6_42490_0_1 | 17.04 | -2.32 | 0.71 | -3.25 | 1.14E-03 | 9.49E-03 |
| dd_Smed_v6_75_0_1 | 418.07 | -2.32 | 0.41 | -5.61 | 1.98E-08 | 2.00E-06 |
| dd_Smed_v6_1975_0_1 | 127.28 | -2.32 | 0.55 | -4.19 | 2.73E-05 | 5.77E-04 |
| dd_Smed_v6_2841_0_1 | 24.50 | -2.31 | 0.71 | -3.24 | 1.18E-03 | 9.74E-03 |
| dd_Smed_v6_500_0_1 | 197.78 | -2.31 | 0.39 | -5.89 | 3.75E-09 | 4.62E-07 |
| dd_Smed_v6_9875_0_1 | 117.70 | -2.30 | 0.67 | -3.43 | 6.02E-04 | 5.96E-03 |
| dd_Smed_v6_9781_0_1 | 27.83 | -2.29 | 0.70 | -3.29 | 9.97E-04 | 8.57E-03 |
| dd_Smed_v6_2516_0_1 | 126.23 | -2.28 | 0.52 | -4.37 | 1.22E-05 | 3.15E-04 |
| dd_Smed_v6_13910_0_1 | 64.94 | -2.28 | 0.64 | -3.58 | 3.42E-04 | 3.91E-03 |
| dd_Smed_v6_7746_0_1 | 57.11 | -2.27 | 0.42 | -5.42 | 6.00E-08 | 4.88E-06 |
| dd_Smed_v6_6337_0_1 | 183.69 | -2.26 | 0.64 | -3.54 | 3.94E-04 | 4.34E-03 |
| dd_Smed_v6_1688_0_1 | 70.89 | -2.26 | 0.55 | -4.08 | 4.46E-05 | 8.31E-04 |
| dd_Smed_v6_8133_0_1 | 133.00 | -2.25 | 0.68 | -3.29 | 1.01E-03 | 8.64E-03 |
| dd_Smed_v6_15909_0_1 | 42.36 | -2.25 | 0.56 | -4.03 | 5.68E-05 | 1.01E-03 |
| dd_Smed_v6_7311_0_1 | 139.22 | -2.22 | 0.68 | -3.25 | 1.16E-03 | 9.63E-03 |
| dd_Smed_v6_2084_0_1 | 171.10 | -2.21 | 0.50 | -4.44 | 8.92E-06 | 2.47E-04 |
| dd_Smed_v6_15254_0_1 | 22.80 | -2.20 | 0.67 | -3.27 | 1.08E-03 | 9.11E-03 |
| dd_Smed_v6_5233_0_1 | 118.55 | -2.20 | 0.60 | -3.69 | 2.25E-04 | 2.88E-03 |
| dd_Smed_v6_100_0_1 | 511.07 | -2.19 | 0.43 | -5.13 | 2.93E-07 | 1.73E-05 |
| dd_Smed_v6_194_0_1 | 580.61 | -2.18 | 0.43 | -5.09 | 3.52E-07 | 1.98E-05 |
| dd_Smed_v6_5261_0_1 | 65.17 | -2.18 | 0.51 | -4.26 | 2.06E-05 | 4.66E-04 |
| dd_Smed_v6_9402_0_1 | 115.67 | -2.16 | 0.60 | -3.63 | 2.89E-04 | 3.43E-03 |
| dd_Smed_v6_6092_0_1 | 102.04 | -2.15 | 0.56 | -3.87 | 1.08E-04 | 1.68E-03 |
| dd_Smed_v6_10359_0_1 | 90.99 | -2.15 | 0.40 | -5.38 | 7.64E-08 | 5.89E-06 |
| dd_Smed_v6_652_0_1 | 600.56 | -2.14 | 0.58 | -3.67 | 2.43E-04 | 3.03E-03 |
| dd_Smed_v6_6028_0_1 | 63.88 | -2.14 | 0.57 | -3.72 | 2.02E-04 | 2.67E-03 |
| dd_Smed_v6_13542_0_1 | 43.18 | -2.13 | 0.60 | -3.58 | 3.44E-04 | 3.93E-03 |
| dd_Smed_v6_2258_0_1 | 58.71 | -2.13 | 0.60 | -3.56 | 3.70E-04 | 4.13E-03 |
| dd_Smed_v6_583_0_1 | 219.31 | -2.12 | 0.46 | -4.58 | 4.73E-06 | 1.52E-04 |
| dd_Smed_v6_9425_1_1 | 38.43 | -2.12 | 0.60 | -3.54 | 4.03E-04 | 4.41E-03 |
| dd_Smed_v6_11120_0_1 | 21.12 | -2.12 | 0.58 | -3.66 | 2.56E-04 | 3.14E-03 |
| dd_Smed_v6_4905_0_1 | 176.47 | -2.11 | 0.44 | -4.83 | 1.35E-06 | 5.65E-05 |

| | | | | | | |
|---|---|---|---|---|---|---|
| dd_Smed_v6_1_0_1 | 1322.14 | -2.10 | 0.39 | -5.40 | 6.69E-08 | 5.30E-06 |
| dd_Smed_v6_1774_0_1 | 61.77 | -2.08 | 0.62 | -3.37 | 7.63E-04 | 7.05E-03 |
| dd_Smed_v6_3640_0_1 | 90.30 | -2.08 | 0.63 | -3.28 | 1.03E-03 | 8.80E-03 |
| dd_Smed_v6_6483_0_1 | 82.88 | -2.06 | 0.60 | -3.43 | 6.05E-04 | 5.97E-03 |
| dd_Smed_v6_6221_0_1 | 142.75 | -2.06 | 0.57 | -3.61 | 3.08E-04 | 3.59E-03 |
| dd_Smed_v6_44_0_1 | 1153.21 | -2.06 | 0.38 | -5.48 | 4.32E-08 | 3.82E-06 |
| dd_Smed_v6_10244_0_1 | 31.49 | -2.06 | 0.48 | -4.27 | 1.92E-05 | 4.43E-04 |
| dd_Smed_v6_9565_0_1 | 207.60 | -2.06 | 0.59 | -3.49 | 4.77E-04 | 5.00E-03 |
| dd_Smed_v6_11493_0_1 | 68.61 | -2.05 | 0.58 | -3.54 | 4.06E-04 | 4.43E-03 |
| dd_Smed_v6_267_0_1 | 796.26 | -2.04 | 0.38 | -5.36 | 8.52E-08 | 6.47E-06 |
| dd_Smed_v6_48_0_1 | 2177.38 | -2.04 | 0.36 | -5.64 | 1.73E-08 | 1.80E-06 |
| dd_Smed_v6_3774_0_1 | 274.49 | -2.04 | 0.56 | -3.62 | 2.99E-04 | 3.52E-03 |
| dd_Smed_v6_4006_0_1 | 120.06 | -2.03 | 0.60 | -3.36 | 7.77E-04 | 7.14E-03 |
| dd_Smed_v6_5339_0_1 | 70.40 | -2.02 | 0.57 | -3.57 | 3.59E-04 | 4.05E-03 |
| dd_Smed_v6_172_1_1 | 421.53 | -2.02 | 0.49 | -4.15 | 3.27E-05 | 6.58E-04 |
| dd_Smed_v6_3948_0_1 | 177.81 | -2.01 | 0.53 | -3.78 | 1.55E-04 | 2.20E-03 |
| dd_Smed_v6_396_0_1 | 346.64 | -2.01 | 0.43 | -4.74 | 2.17E-06 | 8.43E-05 |
| dd_Smed_v6_14698_0_1 | 39.86 | -2.01 | 0.54 | -3.75 | 1.77E-04 | 2.43E-03 |
| dd_Smed_v6_13333_0_1 | 184.36 | -2.01 | 0.59 | -3.43 | 6.13E-04 | 6.04E-03 |
| dd_Smed_v6_6169_0_1 | 166.13 | -2.00 | 0.42 | -4.72 | 2.40E-06 | 9.08E-05 |
| dd_Smed_v6_1581_0_1 | 861.46 | -2.00 | 0.41 | -4.89 | 1.03E-06 | 4.58E-05 |
| dd_Smed_v6_427_0_1 | 136.51 | -2.00 | 0.47 | -4.22 | 2.45E-05 | 5.28E-04 |
| dd_Smed_v6_13519_0_1 | 64.92 | -1.99 | 0.59 | -3.40 | 6.71E-04 | 6.41E-03 |
| dd_Smed_v6_831_0_1 | 108.46 | -1.99 | 0.57 | -3.52 | 4.34E-04 | 4.63E-03 |
| dd_Smed_v6_1054_0_1 | 1534.26 | -1.99 | 0.60 | -3.30 | 9.66E-04 | 8.40E-03 |
| dd_Smed_v6_257_0_1 | 179.78 | -1.99 | 0.59 | -3.37 | 7.43E-04 | 6.90E-03 |
| dd_Smed_v6_8077_0_1 | 36.90 | -1.99 | 0.61 | -3.27 | 1.09E-03 | 9.22E-03 |
| dd_Smed_v6_636_0_1 | 330.89 | -1.98 | 0.44 | -4.53 | 6.02E-06 | 1.85E-04 |
| dd_Smed_v6_2305_0_1 | 232.02 | -1.98 | 0.48 | -4.10 | 4.19E-05 | 7.96E-04 |
| dd_Smed_v6_399_0_1 | 430.07 | -1.98 | 0.45 | -4.37 | 1.26E-05 | 3.24E-04 |
| dd_Smed_v6_3919_0_1 | 51.43 | -1.97 | 0.42 | -4.66 | 3.17E-06 | 1.13E-04 |
| dd_Smed_v6_1570_0_1 | 131.74 | -1.96 | 0.48 | -4.11 | 3.93E-05 | 7.60E-04 |
| dd_Smed_v6_1170_0_1 | 620.36 | -1.96 | 0.40 | -4.87 | 1.10E-06 | 4.82E-05 |
| dd_Smed_v6_20344_0_1 | 50.45 | -1.96 | 0.56 | -3.51 | 4.41E-04 | 4.69E-03 |
| dd_Smed_v6_3172_0_1 | 181.19 | -1.96 | 0.56 | -3.47 | 5.30E-04 | 5.42E-03 |
| dd_Smed_v6_1037_0_1 | 191.39 | -1.96 | 0.45 | -4.38 | 1.19E-05 | 3.07E-04 |
| dd_Smed_v6_1114_0_1 | 479.39 | -1.95 | 0.48 | -4.04 | 5.41E-05 | 9.70E-04 |
| dd_Smed_v6_32_0_1 | 823.79 | -1.95 | 0.43 | -4.53 | 5.88E-06 | 1.82E-04 |
| dd_Smed_v6_1388_0_1 | 71.33 | -1.94 | 0.60 | -3.24 | 1.18E-03 | 9.73E-03 |
| dd_Smed_v6_2993_0_1 | 113.56 | -1.94 | 0.60 | -3.24 | 1.21E-03 | 9.88E-03 |
| dd_Smed_v6_1224_0_1 | 662.92 | -1.94 | 0.39 | -4.96 | 7.04E-07 | 3.45E-05 |
| dd_Smed_v6_808_0_1 | 106.99 | -1.93 | 0.57 | -3.39 | 6.91E-04 | 6.56E-03 |
| dd_Smed_v6_184_0_1 | 1331.11 | -1.91 | 0.53 | -3.64 | 2.72E-04 | 3.28E-03 |

| | | | | | | |
|---|---|---|---|---|---|---|
| dd_Smed_v6_1789_0_1 | 571.57 | -1.91 | 0.45 | -4.27 | 1.96E-05 | 4.49E-04 |
| dd_Smed_v6_8833_0_1 | 299.76 | -1.91 | 0.52 | -3.70 | 2.18E-04 | 2.81E-03 |
| dd_Smed_v6_917_0_1 | 154.81 | -1.91 | 0.50 | -3.83 | 1.28E-04 | 1.92E-03 |
| dd_Smed_v6_20_0_1 | 3054.14 | -1.91 | 0.38 | -5.08 | 3.85E-07 | 2.13E-05 |
| dd_Smed_v6_12027_0_1 | 84.70 | -1.90 | 0.38 | -5.07 | 4.04E-07 | 2.22E-05 |
| dd_Smed_v6_1962_0_1 | 225.18 | -1.90 | 0.48 | -3.93 | 8.33E-05 | 1.35E-03 |
| dd_Smed_v6_2554_0_1 | 228.69 | -1.90 | 0.56 | -3.40 | 6.65E-04 | 6.37E-03 |
| dd_Smed_v6_3234_0_1 | 116.54 | -1.90 | 0.55 | -3.47 | 5.28E-04 | 5.41E-03 |
| dd_Smed_v6_6007_0_1 | 225.53 | -1.89 | 0.54 | -3.48 | 4.97E-04 | 5.17E-03 |
| dd_Smed_v6_12212_0_1 | 331.27 | -1.89 | 0.45 | -4.19 | 2.78E-05 | 5.85E-04 |
| dd_Smed_v6_898_0_1 | 79.84 | -1.89 | 0.58 | -3.27 | 1.08E-03 | 9.14E-03 |
| dd_Smed_v6_1643_0_1 | 145.81 | -1.89 | 0.57 | -3.33 | 8.66E-04 | 7.75E-03 |
| dd_Smed_v6_873_0_1 | 1129.97 | -1.88 | 0.57 | -3.31 | 9.19E-04 | 8.10E-03 |
| dd_Smed_v6_13959_0_1 | 42.76 | -1.87 | 0.54 | -3.46 | 5.50E-04 | 5.56E-03 |
| dd_Smed_v6_6348_0_1 | 115.28 | -1.87 | 0.55 | -3.41 | 6.39E-04 | 6.20E-03 |
| dd_Smed_v6_5949_0_1 | 90.72 | -1.87 | 0.37 | -5.07 | 4.07E-07 | 2.23E-05 |
| dd_Smed_v6_272_0_1 | 164.30 | -1.86 | 0.42 | -4.46 | 8.27E-06 | 2.35E-04 |
| dd_Smed_v6_31140_0_1 | 34.23 | -1.86 | 0.57 | -3.26 | 1.13E-03 | 9.40E-03 |
| dd_Smed_v6_2438_0_1 | 396.92 | -1.86 | 0.38 | -4.84 | 1.32E-06 | 5.53E-05 |
| dd_Smed_v6_814_0_1 | 300.25 | -1.86 | 0.51 | -3.67 | 2.44E-04 | 3.04E-03 |
| dd_Smed_v6_452_0_1 | 331.47 | -1.86 | 0.39 | -4.72 | 2.31E-06 | 8.85E-05 |
| dd_Smed_v6_306_0_1 | 225.89 | -1.86 | 0.57 | -3.28 | 1.03E-03 | 8.80E-03 |
| dd_Smed_v6_8821_0_1 | 101.43 | -1.85 | 0.42 | -4.45 | 8.43E-06 | 2.37E-04 |
| dd_Smed_v6_1942_0_1 | 259.45 | -1.85 | 0.56 | -3.32 | 8.93E-04 | 7.92E-03 |
| dd_Smed_v6_1749_0_1 | 223.91 | -1.85 | 0.50 | -3.73 | 1.93E-04 | 2.60E-03 |
| dd_Smed_v6_10387_0_1 | 78.85 | -1.84 | 0.40 | -4.58 | 4.72E-06 | 1.52E-04 |
| dd_Smed_v6_4841_0_1 | 203.22 | -1.84 | 0.52 | -3.55 | 3.88E-04 | 4.28E-03 |
| dd_Smed_v6_1971_0_1 | 313.47 | -1.84 | 0.47 | -3.90 | 9.51E-05 | 1.52E-03 |
| dd_Smed_v6_1088_0_1 | 175.45 | -1.83 | 0.43 | -4.28 | 1.90E-05 | 4.40E-04 |
| dd_Smed_v6_5349_0_1 | 180.64 | -1.83 | 0.49 | -3.71 | 2.06E-04 | 2.71E-03 |
| dd_Smed_v6_6324_0_1 | 38.96 | -1.83 | 0.55 | -3.33 | 8.55E-04 | 7.68E-03 |
| dd_Smed_v6_1726_0_1 | 281.97 | -1.82 | 0.41 | -4.42 | 1.01E-05 | 2.71E-04 |
| dd_Smed_v6_4437_0_1 | 112.96 | -1.82 | 0.56 | -3.26 | 1.13E-03 | 9.44E-03 |
| dd_Smed_v6_12567_0_1 | 96.00 | -1.81 | 0.55 | -3.31 | 9.38E-04 | 8.23E-03 |
| dd_Smed_v6_11_0_1 | 1466.98 | -1.81 | 0.37 | -4.92 | 8.71E-07 | 4.04E-05 |
| dd_Smed_v6_54_0_1 | 1086.50 | -1.81 | 0.33 | -5.45 | 4.95E-08 | 4.26E-06 |
| dd_Smed_v6_2282_0_1 | 377.15 | -1.81 | 0.42 | -4.25 | 2.10E-05 | 4.70E-04 |
| dd_Smed_v6_16492_0_1 | 54.38 | -1.81 | 0.51 | -3.55 | 3.81E-04 | 4.22E-03 |
| dd_Smed_v6_1371_0_1 | 243.14 | -1.81 | 0.50 | -3.64 | 2.69E-04 | 3.26E-03 |
| dd_Smed_v6_6149_0_1 | 153.75 | -1.80 | 0.50 | -3.63 | 2.84E-04 | 3.38E-03 |
| dd_Smed_v6_12105_0_1 | 271.99 | -1.80 | 0.47 | -3.84 | 1.21E-04 | 1.84E-03 |
| dd_Smed_v6_3670_0_1 | 480.06 | -1.79 | 0.48 | -3.70 | 2.17E-04 | 2.80E-03 |
| dd_Smed_v6_9780_0_1 | 69.71 | -1.79 | 0.51 | -3.54 | 4.06E-04 | 4.43E-03 |

| | | | | | | |
|---|---|---|---|---|---|---|
| dd_Smed_v6_642_0_1 | 987.83 | -1.78 | 0.54 | -3.32 | 9.00E-04 | 7.96E-03 |
| dd_Smed_v6_3441_0_1 | 206.87 | -1.77 | 0.50 | -3.57 | 3.63E-04 | 4.07E-03 |
| dd_Smed_v6_3177_0_1 | 381.08 | -1.77 | 0.44 | -4.07 | 4.72E-05 | 8.72E-04 |
| dd_Smed_v6_5168_0_1 | 162.33 | -1.77 | 0.46 | -3.85 | 1.20E-04 | 1.84E-03 |
| dd_Smed_v6_2229_0_1 | 151.33 | -1.76 | 0.53 | -3.30 | 9.74E-04 | 8.44E-03 |
| dd_Smed_v6_2604_0_1 | 170.57 | -1.76 | 0.50 | -3.53 | 4.14E-04 | 4.49E-03 |
| dd_Smed_v6_9521_0_1 | 111.61 | -1.76 | 0.53 | -3.29 | 1.00E-03 | 8.59E-03 |
| dd_Smed_v6_4618_0_1 | 204.61 | -1.76 | 0.39 | -4.49 | 7.04E-06 | 2.09E-04 |
| dd_Smed_v6_81_0_1 | 2115.97 | -1.74 | 0.48 | -3.61 | 3.05E-04 | 3.57E-03 |
| dd_Smed_v6_2929_0_1 | 138.26 | -1.73 | 0.45 | -3.84 | 1.25E-04 | 1.89E-03 |
| dd_Smed_v6_680_0_1 | 434.99 | -1.73 | 0.49 | -3.53 | 4.09E-04 | 4.45E-03 |
| dd_Smed_v6_1616_0_1 | 94.01 | -1.73 | 0.51 | -3.38 | 7.35E-04 | 6.86E-03 |
| dd_Smed_v6_13702_0_1 | 70.09 | -1.73 | 0.48 | -3.57 | 3.55E-04 | 4.01E-03 |
| dd_Smed_v6_2734_0_1 | 229.67 | -1.73 | 0.37 | -4.66 | 3.17E-06 | 1.13E-04 |
| dd_Smed_v6_48911_0_1 | 31.01 | -1.72 | 0.52 | -3.30 | 9.53E-04 | 8.32E-03 |
| dd_Smed_v6_3788_0_1 | 133.21 | -1.72 | 0.47 | -3.70 | 2.18E-04 | 2.81E-03 |
| dd_Smed_v6_3852_0_1 | 249.33 | -1.72 | 0.52 | -3.29 | 1.00E-03 | 8.61E-03 |
| dd_Smed_v6_7495_0_1 | 156.86 | -1.72 | 0.40 | -4.28 | 1.87E-05 | 4.36E-04 |
| dd_Smed_v6_1306_0_1 | 401.18 | -1.72 | 0.50 | -3.45 | 5.58E-04 | 5.61E-03 |
| dd_Smed_v6_983_0_1 | 503.09 | -1.72 | 0.43 | -4.03 | 5.59E-05 | 9.93E-04 |
| dd_Smed_v6_1735_0_1 | 273.64 | -1.71 | 0.52 | -3.30 | 9.60E-04 | 8.36E-03 |
| dd_Smed_v6_241_0_1 | 1581.56 | -1.71 | 0.36 | -4.78 | 1.76E-06 | 7.00E-05 |
| dd_Smed_v6_23532_0_1 | 121.22 | -1.71 | 0.45 | -3.82 | 1.35E-04 | 1.99E-03 |
| dd_Smed_v6_5204_0_1 | 123.04 | -1.70 | 0.48 | -3.54 | 3.98E-04 | 4.38E-03 |
| dd_Smed_v6_1422_0_1 | 420.71 | -1.70 | 0.44 | -3.86 | 1.15E-04 | 1.77E-03 |
| dd_Smed_v6_212_0_1 | 381.86 | -1.70 | 0.46 | -3.69 | 2.26E-04 | 2.88E-03 |
| dd_Smed_v6_6436_0_1 | 136.06 | -1.70 | 0.41 | -4.18 | 2.95E-05 | 6.08E-04 |
| dd_Smed_v6_124_0_1 | 2790.92 | -1.70 | 0.33 | -5.18 | 2.25E-07 | 1.42E-05 |
| dd_Smed_v6_646_0_1 | 653.89 | -1.69 | 0.44 | -3.85 | 1.20E-04 | 1.84E-03 |
| dd_Smed_v6_1426_0_1 | 295.57 | -1.69 | 0.38 | -4.41 | 1.01E-05 | 2.72E-04 |
| dd_Smed_v6_13408_0_1 | 259.72 | -1.68 | 0.38 | -4.48 | 7.58E-06 | 2.20E-04 |
| dd_Smed_v6_2234_0_1 | 346.05 | -1.67 | 0.51 | -3.30 | 9.60E-04 | 8.36E-03 |
| dd_Smed_v6_2237_0_1 | 121.87 | -1.67 | 0.48 | -3.45 | 5.52E-04 | 5.56E-03 |
| dd_Smed_v6_4575_0_1 | 242.27 | -1.67 | 0.48 | -3.47 | 5.15E-04 | 5.32E-03 |
| dd_Smed_v6_1275_0_1 | 367.14 | -1.66 | 0.40 | -4.13 | 3.57E-05 | 7.03E-04 |
| dd_Smed_v6_1320_0_1 | 76.99 | -1.66 | 0.44 | -3.80 | 1.46E-04 | 2.11E-03 |
| dd_Smed_v6_6930_0_1 | 71.21 | -1.66 | 0.38 | -4.41 | 1.05E-05 | 2.80E-04 |
| dd_Smed_v6_7028_0_1 | 129.68 | -1.66 | 0.45 | -3.72 | 2.00E-04 | 2.66E-03 |
| dd_Smed_v6_4914_0_1 | 175.33 | -1.66 | 0.46 | -3.57 | 3.55E-04 | 4.01E-03 |
| dd_Smed_v6_1687_0_1 | 142.98 | -1.66 | 0.50 | -3.29 | 9.95E-04 | 8.56E-03 |
| dd_Smed_v6_1706_0_1 | 345.30 | -1.66 | 0.33 | -4.99 | 6.06E-07 | 3.06E-05 |
| dd_Smed_v6_63_0_1 | 1090.42 | -1.65 | 0.37 | -4.52 | 6.28E-06 | 1.90E-04 |
| dd_Smed_v6_4834_0_1 | 86.38 | -1.65 | 0.39 | -4.28 | 1.89E-05 | 4.39E-04 |

| | | | | | | |
|---|---|---|---|---|---|---|
| dd_Smed_v6_1106_0_1 | 766.43 | -1.64 | 0.42 | -3.91 | 9.25E-05 | 1.48E-03 |
| dd_Smed_v6_234_0_1 | 1628.44 | -1.64 | 0.48 | -3.43 | 6.02E-04 | 5.96E-03 |
| dd_Smed_v6_116_0_1 | 503.12 | -1.64 | 0.46 | -3.58 | 3.49E-04 | 3.96E-03 |
| dd_Smed_v6_4827_0_1 | 62.47 | -1.64 | 0.50 | -3.27 | 1.07E-03 | 9.07E-03 |
| dd_Smed_v6_1245_0_1 | 201.28 | -1.64 | 0.49 | -3.33 | 8.62E-04 | 7.72E-03 |
| dd_Smed_v6_866_0_1 | 334.95 | -1.63 | 0.40 | -4.12 | 3.73E-05 | 7.28E-04 |
| dd_Smed_v6_3051_0_1 | 82.27 | -1.63 | 0.48 | -3.41 | 6.47E-04 | 6.25E-03 |
| dd_Smed_v6_5728_0_1 | 279.77 | -1.63 | 0.46 | -3.58 | 3.48E-04 | 3.96E-03 |
| dd_Smed_v6_485_0_1 | 648.66 | -1.63 | 0.45 | -3.64 | 2.73E-04 | 3.29E-03 |
| dd_Smed_v6_12378_0_1 | 55.09 | -1.62 | 0.39 | -4.16 | 3.17E-05 | 6.42E-04 |
| dd_Smed_v6_1866_0_1 | 268.77 | -1.62 | 0.42 | -3.83 | 1.27E-04 | 1.91E-03 |
| dd_Smed_v6_8169_0_1 | 130.21 | -1.62 | 0.40 | -4.08 | 4.56E-05 | 8.48E-04 |
| dd_Smed_v6_994_0_1 | 193.28 | -1.62 | 0.44 | -3.66 | 2.51E-04 | 3.10E-03 |
| dd_Smed_v6_8131_0_1 | 286.99 | -1.62 | 0.35 | -4.61 | 3.98E-06 | 1.36E-04 |
| dd_Smed_v6_780_0_1 | 511.78 | -1.61 | 0.41 | -3.90 | 9.71E-05 | 1.55E-03 |
| dd_Smed_v6_1120_0_1 | 87.23 | -1.61 | 0.34 | -4.71 | 2.49E-06 | 9.35E-05 |
| dd_Smed_v6_2291_0_1 | 118.73 | -1.61 | 0.36 | -4.46 | 8.22E-06 | 2.35E-04 |
| dd_Smed_v6_8356_0_1 | 198.06 | -1.61 | 0.49 | -3.30 | 9.81E-04 | 8.49E-03 |
| dd_Smed_v6_11281_1_1 | 79.67 | -1.60 | 0.48 | -3.37 | 7.61E-04 | 7.04E-03 |
| dd_Smed_v6_107_0_1 | 338.78 | -1.60 | 0.49 | -3.27 | 1.09E-03 | 9.20E-03 |
| dd_Smed_v6_5884_0_1 | 147.27 | -1.60 | 0.45 | -3.60 | 3.24E-04 | 3.74E-03 |
| dd_Smed_v6_5218_0_1 | 509.41 | -1.60 | 0.45 | -3.55 | 3.85E-04 | 4.25E-03 |
| dd_Smed_v6_2328_0_1 | 219.82 | -1.59 | 0.45 | -3.57 | 3.54E-04 | 4.00E-03 |
| dd_Smed_v6_785_0_1 | 695.81 | -1.59 | 0.43 | -3.70 | 2.16E-04 | 2.80E-03 |
| dd_Smed_v6_2871_0_1 | 149.87 | -1.59 | 0.48 | -3.32 | 8.89E-04 | 7.90E-03 |
| dd_Smed_v6_1565_0_1 | 101.66 | -1.58 | 0.44 | -3.56 | 3.65E-04 | 4.08E-03 |
| dd_Smed_v6_561_0_1 | 429.49 | -1.57 | 0.43 | -3.63 | 2.79E-04 | 3.35E-03 |
| dd_Smed_v6_749_0_1 | 842.37 | -1.57 | 0.37 | -4.29 | 1.81E-05 | 4.27E-04 |
| dd_Smed_v6_2092_0_1 | 373.44 | -1.56 | 0.34 | -4.54 | 5.51E-06 | 1.72E-04 |
| dd_Smed_v6_7008_0_1 | 72.10 | -1.56 | 0.45 | -3.44 | 5.82E-04 | 5.79E-03 |
| dd_Smed_v6_11523_0_1 | 112.00 | -1.56 | 0.47 | -3.36 | 7.93E-04 | 7.22E-03 |
| dd_Smed_v6_2083_0_1 | 131.03 | -1.55 | 0.44 | -3.54 | 4.01E-04 | 4.39E-03 |
| dd_Smed_v6_16007_0_1 | 126.92 | -1.55 | 0.44 | -3.51 | 4.52E-04 | 4.80E-03 |
| dd_Smed_v6_7811_0_1 | 875.16 | -1.54 | 0.38 | -4.04 | 5.38E-05 | 9.67E-04 |
| dd_Smed_v6_7574_0_1 | 79.40 | -1.54 | 0.42 | -3.65 | 2.65E-04 | 3.23E-03 |
| dd_Smed_v6_1760_1_1 | 130.05 | -1.54 | 0.39 | -3.93 | 8.46E-05 | 1.37E-03 |
| dd_Smed_v6_531_0_1 | 198.93 | -1.54 | 0.46 | -3.33 | 8.72E-04 | 7.79E-03 |
| dd_Smed_v6_8623_0_1 | 109.18 | -1.53 | 0.47 | -3.25 | 1.15E-03 | 9.59E-03 |
| dd_Smed_v6_353_0_1 | 595.76 | -1.53 | 0.36 | -4.23 | 2.36E-05 | 5.13E-04 |
| dd_Smed_v6_683_0_1 | 701.90 | -1.53 | 0.44 | -3.48 | 5.02E-04 | 5.21E-03 |
| dd_Smed_v6_361_0_1 | 602.78 | -1.53 | 0.44 | -3.48 | 4.99E-04 | 5.19E-03 |
| dd_Smed_v6_595_0_1 | 170.91 | -1.52 | 0.45 | -3.38 | 7.37E-04 | 6.86E-03 |
| dd_Smed_v6_219_0_1 | 584.84 | -1.52 | 0.32 | -4.74 | 2.09E-06 | 8.16E-05 |

| | | | | | | |
|---|---|---|---|---|---|---|
| dd_Smed_v6_1132_0_1 | 992.61 | -1.52 | 0.41 | -3.70 | 2.12E-04 | 2.75E-03 |
| dd_Smed_v6_367_0_1 | 284.31 | -1.51 | 0.41 | -3.72 | 1.96E-04 | 2.62E-03 |
| dd_Smed_v6_390_0_1 | 646.41 | -1.51 | 0.42 | -3.61 | 3.10E-04 | 3.61E-03 |
| dd_Smed_v6_4404_0_1 | 314.59 | -1.51 | 0.43 | -3.51 | 4.54E-04 | 4.82E-03 |
| dd_Smed_v6_626_0_1 | 405.41 | -1.51 | 0.41 | -3.68 | 2.34E-04 | 2.95E-03 |
| dd_Smed_v6_4297_0_1 | 112.92 | -1.51 | 0.45 | -3.36 | 7.78E-04 | 7.14E-03 |
| dd_Smed_v6_459_0_1 | 554.17 | -1.50 | 0.42 | -3.54 | 3.99E-04 | 4.38E-03 |
| dd_Smed_v6_694_0_1 | 114.01 | -1.49 | 0.45 | -3.29 | 9.94E-04 | 8.55E-03 |
| dd_Smed_v6_5621_0_1 | 118.18 | -1.48 | 0.32 | -4.61 | 4.02E-06 | 1.36E-04 |
| dd_Smed_v6_743_0_1 | 268.84 | -1.47 | 0.43 | -3.44 | 5.79E-04 | 5.78E-03 |
| dd_Smed_v6_3281_0_1 | 198.63 | -1.47 | 0.41 | -3.56 | 3.70E-04 | 4.13E-03 |
| dd_Smed_v6_16850_0_1 | 333.47 | -1.47 | 0.45 | -3.24 | 1.19E-03 | 9.75E-03 |
| dd_Smed_v6_12339_0_1 | 106.00 | -1.47 | 0.30 | -4.83 | 1.38E-06 | 5.73E-05 |
| dd_Smed_v6_3311_0_1 | 161.99 | -1.47 | 0.39 | -3.75 | 1.74E-04 | 2.40E-03 |
| dd_Smed_v6_1291_0_1 | 136.35 | -1.46 | 0.35 | -4.23 | 2.29E-05 | 5.03E-04 |
| dd_Smed_v6_1579_0_1 | 4386.17 | -1.46 | 0.33 | -4.44 | 9.20E-06 | 2.53E-04 |
| dd_Smed_v6_7614_0_1 | 367.16 | -1.45 | 0.23 | -6.44 | 1.19E-10 | 2.29E-08 |
| dd_Smed_v6_8179_0_1 | 105.88 | -1.45 | 0.39 | -3.77 | 1.66E-04 | 2.32E-03 |
| dd_Smed_v6_3419_0_1 | 220.54 | -1.43 | 0.36 | -3.96 | 7.64E-05 | 1.26E-03 |
| dd_Smed_v6_1155_0_1 | 282.72 | -1.43 | 0.34 | -4.20 | 2.65E-05 | 5.65E-04 |
| dd_Smed_v6_3090_0_1 | 169.02 | -1.43 | 0.42 | -3.43 | 5.96E-04 | 5.91E-03 |
| dd_Smed_v6_3572_0_1 | 215.66 | -1.42 | 0.36 | -3.96 | 7.35E-05 | 1.23E-03 |
| dd_Smed_v6_1625_0_1 | 231.71 | -1.42 | 0.28 | -5.02 | 5.09E-07 | 2.66E-05 |
| dd_Smed_v6_12797_0_1 | 251.93 | -1.42 | 0.37 | -3.82 | 1.34E-04 | 1.98E-03 |
| dd_Smed_v6_1266_0_1 | 219.71 | -1.42 | 0.34 | -4.23 | 2.29E-05 | 5.03E-04 |
| dd_Smed_v6_11300_0_1 | 220.10 | -1.42 | 0.39 | -3.66 | 2.52E-04 | 3.11E-03 |
| dd_Smed_v6_862_0_1 | 162.18 | -1.41 | 0.40 | -3.52 | 4.38E-04 | 4.67E-03 |
| dd_Smed_v6_13789_0_1 | 273.50 | -1.41 | 0.32 | -4.36 | 1.29E-05 | 3.30E-04 |
| dd_Smed_v6_2669_0_1 | 147.40 | -1.40 | 0.42 | -3.38 | 7.23E-04 | 6.78E-03 |
| dd_Smed_v6_824_0_1 | 1088.19 | -1.40 | 0.31 | -4.56 | 5.23E-06 | 1.64E-04 |
| dd_Smed_v6_6712_0_1 | 598.71 | -1.40 | 0.27 | -5.15 | 2.55E-07 | 1.57E-05 |
| dd_Smed_v6_564_0_1 | 121.03 | -1.40 | 0.42 | -3.33 | 8.60E-04 | 7.72E-03 |
| dd_Smed_v6_13571_0_1 | 214.48 | -1.39 | 0.32 | -4.35 | 1.39E-05 | 3.47E-04 |
| dd_Smed_v6_14494_0_1 | 63.87 | -1.39 | 0.40 | -3.50 | 4.69E-04 | 4.94E-03 |
| dd_Smed_v6_10555_0_1 | 151.12 | -1.39 | 0.38 | -3.70 | 2.13E-04 | 2.76E-03 |
| dd_Smed_v6_1351_0_1 | 302.19 | -1.39 | 0.40 | -3.47 | 5.24E-04 | 5.38E-03 |
| dd_Smed_v6_8108_0_1 | 173.84 | -1.39 | 0.33 | -4.19 | 2.83E-05 | 5.91E-04 |
| dd_Smed_v6_2037_0_1 | 240.30 | -1.38 | 0.34 | -4.11 | 3.95E-05 | 7.63E-04 |
| dd_Smed_v6_7992_0_1 | 215.66 | -1.38 | 0.34 | -4.01 | 5.98E-05 | 1.05E-03 |
| dd_Smed_v6_6581_1_1 | 60.28 | -1.38 | 0.42 | -3.27 | 1.07E-03 | 9.08E-03 |
| dd_Smed_v6_11223_0_1 | 291.27 | -1.38 | 0.42 | -3.25 | 1.17E-03 | 9.68E-03 |
| dd_Smed_v6_1897_0_1 | 432.32 | -1.37 | 0.32 | -4.27 | 1.95E-05 | 4.48E-04 |
| dd_Smed_v6_16127_0_1 | 94.20 | -1.37 | 0.40 | -3.44 | 5.80E-04 | 5.78E-03 |

| | | | | | | |
|---|---|---|---|---|---|---|
| dd_Smed_v6_1744_0_1 | 170.26 | -1.37 | 0.42 | -3.24 | 1.19E-03 | 9.74E-03 |
| dd_Smed_v6_6127_0_1 | 100.07 | -1.36 | 0.36 | -3.75 | 1.76E-04 | 2.43E-03 |
| dd_Smed_v6_8917_0_1 | 337.70 | -1.36 | 0.41 | -3.33 | 8.65E-04 | 7.75E-03 |
| dd_Smed_v6_1303_0_1 | 115.60 | -1.36 | 0.38 | -3.62 | 2.91E-04 | 3.44E-03 |
| dd_Smed_v6_540_0_1 | 877.19 | -1.36 | 0.31 | -4.40 | 1.06E-05 | 2.82E-04 |
| dd_Smed_v6_1505_0_1 | 244.64 | -1.36 | 0.32 | -4.29 | 1.79E-05 | 4.24E-04 |
| dd_Smed_v6_11958_0_1 | 232.32 | -1.36 | 0.41 | -3.33 | 8.79E-04 | 7.85E-03 |
| dd_Smed_v6_640_0_1 | 365.12 | -1.36 | 0.34 | -4.02 | 5.85E-05 | 1.03E-03 |
| dd_Smed_v6_14584_0_1 | 149.09 | -1.35 | 0.32 | -4.27 | 1.98E-05 | 4.52E-04 |
| dd_Smed_v6_3854_0_1 | 247.26 | -1.35 | 0.33 | -4.12 | 3.85E-05 | 7.46E-04 |
| dd_Smed_v6_2408_0_1 | 226.63 | -1.35 | 0.36 | -3.78 | 1.55E-04 | 2.20E-03 |
| dd_Smed_v6_5169_0_1 | 348.92 | -1.35 | 0.24 | -5.62 | 1.86E-08 | 1.90E-06 |
| dd_Smed_v6_5882_0_1 | 155.11 | -1.35 | 0.36 | -3.70 | 2.12E-04 | 2.75E-03 |
| dd_Smed_v6_270_0_1 | 565.15 | -1.34 | 0.41 | -3.27 | 1.07E-03 | 9.05E-03 |
| dd_Smed_v6_17_0_1 | 1477.13 | -1.34 | 0.32 | -4.19 | 2.74E-05 | 5.79E-04 |
| dd_Smed_v6_7213_0_1 | 111.65 | -1.34 | 0.40 | -3.36 | 7.76E-04 | 7.14E-03 |
| dd_Smed_v6_3201_0_1 | 87.02 | -1.34 | 0.37 | -3.63 | 2.80E-04 | 3.36E-03 |
| dd_Smed_v6_8243_0_1 | 124.27 | -1.33 | 0.37 | -3.60 | 3.14E-04 | 3.64E-03 |
| dd_Smed_v6_6476_0_1 | 174.34 | -1.33 | 0.40 | -3.34 | 8.42E-04 | 7.60E-03 |
| dd_Smed_v6_3591_0_1 | 488.64 | -1.32 | 0.37 | -3.53 | 4.17E-04 | 4.52E-03 |
| dd_Smed_v6_9018_0_1 | 283.82 | -1.32 | 0.26 | -4.98 | 6.40E-07 | 3.20E-05 |
| dd_Smed_v6_3658_0_1 | 366.77 | -1.31 | 0.37 | -3.51 | 4.55E-04 | 4.83E-03 |
| dd_Smed_v6_15371_0_1 | 78.23 | -1.31 | 0.35 | -3.74 | 1.81E-04 | 2.47E-03 |
| dd_Smed_v6_2109_0_1 | 1089.10 | -1.31 | 0.37 | -3.54 | 4.01E-04 | 4.39E-03 |
| dd_Smed_v6_4539_0_1 | 511.60 | -1.31 | 0.31 | -4.20 | 2.69E-05 | 5.71E-04 |
| dd_Smed_v6_1067_0_1 | 158.34 | -1.31 | 0.35 | -3.73 | 1.89E-04 | 2.56E-03 |
| dd_Smed_v6_11425_0_1 | 283.28 | -1.31 | 0.27 | -4.76 | 1.94E-06 | 7.66E-05 |
| dd_Smed_v6_810_0_1 | 1472.95 | -1.30 | 0.36 | -3.67 | 2.40E-04 | 3.01E-03 |
| dd_Smed_v6_8449_0_1 | 59.41 | -1.30 | 0.38 | -3.46 | 5.31E-04 | 5.42E-03 |
| dd_Smed_v6_2391_0_1 | 276.63 | -1.30 | 0.33 | -3.97 | 7.18E-05 | 1.21E-03 |
| dd_Smed_v6_3_0_1 | 25838.32 | -1.30 | 0.29 | -4.49 | 6.96E-06 | 2.07E-04 |
| dd_Smed_v6_576_0_1 | 455.87 | -1.29 | 0.38 | -3.39 | 7.00E-04 | 6.63E-03 |
| dd_Smed_v6_1970_0_1 | 363.70 | -1.29 | 0.29 | -4.42 | 9.79E-06 | 2.65E-04 |
| dd_Smed_v6_14112_0_1 | 40.27 | -1.29 | 0.37 | -3.47 | 5.27E-04 | 5.41E-03 |
| dd_Smed_v6_8138_0_1 | 492.47 | -1.28 | 0.34 | -3.80 | 1.46E-04 | 2.11E-03 |
| dd_Smed_v6_593_1_1 | 429.50 | -1.28 | 0.30 | -4.26 | 2.02E-05 | 4.57E-04 |
| dd_Smed_v6_11837_0_1 | 190.18 | -1.27 | 0.35 | -3.68 | 2.29E-04 | 2.90E-03 |
| dd_Smed_v6_8711_0_1 | 96.85 | -1.27 | 0.36 | -3.52 | 4.34E-04 | 4.63E-03 |
| dd_Smed_v6_7153_0_1 | 473.53 | -1.27 | 0.26 | -4.85 | 1.26E-06 | 5.39E-05 |
| dd_Smed_v6_1294_0_1 | 361.87 | -1.27 | 0.27 | -4.76 | 1.93E-06 | 7.62E-05 |
| dd_Smed_v6_2622_0_1 | 1147.45 | -1.27 | 0.21 | -6.17 | 7.03E-10 | 1.11E-07 |
| dd_Smed_v6_11023_0_1 | 201.51 | -1.27 | 0.25 | -5.01 | 5.54E-07 | 2.87E-05 |
| dd_Smed_v6_6736_0_1 | 645.22 | -1.27 | 0.31 | -4.06 | 4.88E-05 | 8.93E-04 |

| | | | | | | |
|---|---|---|---|---|---|---|
| dd_Smed_v6_10425_0_1 | 206.43 | -1.27 | 0.34 | -3.77 | 1.62E-04 | 2.27E-03 |
| dd_Smed_v6_2803_0_1 | 181.17 | -1.26 | 0.38 | -3.34 | 8.46E-04 | 7.62E-03 |
| dd_Smed_v6_229_0_1 | 256.52 | -1.25 | 0.37 | -3.38 | 7.37E-04 | 6.86E-03 |
| dd_Smed_v6_12305_0_1 | 219.56 | -1.24 | 0.35 | -3.57 | 3.60E-04 | 4.06E-03 |
| dd_Smed_v6_3070_0_1 | 987.98 | -1.24 | 0.28 | -4.47 | 8.01E-06 | 2.30E-04 |
| dd_Smed_v6_7663_0_1 | 321.23 | -1.24 | 0.31 | -3.99 | 6.50E-05 | 1.12E-03 |
| dd_Smed_v6_990_0_1 | 844.68 | -1.24 | 0.34 | -3.67 | 2.42E-04 | 3.03E-03 |
| dd_Smed_v6_4614_0_1 | 174.50 | -1.23 | 0.32 | -3.91 | 9.31E-05 | 1.49E-03 |
| dd_Smed_v6_3230_0_1 | 295.40 | -1.23 | 0.35 | -3.47 | 5.14E-04 | 5.30E-03 |
| dd_Smed_v6_2732_0_1 | 368.99 | -1.23 | 0.23 | -5.27 | 1.36E-07 | 9.53E-06 |
| dd_Smed_v6_1008_0_1 | 221.59 | -1.23 | 0.29 | -4.25 | 2.18E-05 | 4.85E-04 |
| dd_Smed_v6_5524_0_1 | 273.28 | -1.22 | 0.27 | -4.58 | 4.75E-06 | 1.52E-04 |
| dd_Smed_v6_6776_0_1 | 122.70 | -1.22 | 0.33 | -3.66 | 2.56E-04 | 3.15E-03 |
| dd_Smed_v6_5498_0_1 | 485.25 | -1.21 | 0.37 | -3.28 | 1.05E-03 | 8.93E-03 |
| dd_Smed_v6_11948_0_1 | 375.22 | -1.21 | 0.30 | -4.09 | 4.28E-05 | 8.08E-04 |
| dd_Smed_v6_10075_0_1 | 168.60 | -1.21 | 0.33 | -3.65 | 2.61E-04 | 3.19E-03 |
| dd_Smed_v6_354_0_1 | 878.58 | -1.21 | 0.20 | -5.91 | 3.50E-09 | 4.42E-07 |
| dd_Smed_v6_12635_0_1 | 94.12 | -1.21 | 0.37 | -3.27 | 1.08E-03 | 9.16E-03 |
| dd_Smed_v6_9419_0_1 | 341.60 | -1.21 | 0.35 | -3.48 | 5.02E-04 | 5.21E-03 |
| dd_Smed_v6_5065_0_1 | 217.24 | -1.20 | 0.36 | -3.36 | 7.73E-04 | 7.12E-03 |
| dd_Smed_v6_800_0_1 | 418.08 | -1.20 | 0.26 | -4.65 | 3.30E-06 | 1.16E-04 |
| dd_Smed_v6_5767_0_1 | 414.01 | -1.19 | 0.25 | -4.69 | 2.69E-06 | 9.91E-05 |
| dd_Smed_v6_3752_0_1 | 189.59 | -1.19 | 0.24 | -4.94 | 7.65E-07 | 3.65E-05 |
| dd_Smed_v6_11428_0_1 | 142.30 | -1.19 | 0.34 | -3.46 | 5.37E-04 | 5.47E-03 |
| dd_Smed_v6_2523_0_1 | 591.24 | -1.18 | 0.21 | -5.67 | 1.45E-08 | 1.52E-06 |
| dd_Smed_v6_10032_0_1 | 413.19 | -1.18 | 0.28 | -4.19 | 2.83E-05 | 5.91E-04 |
| dd_Smed_v6_6322_0_1 | 102.13 | -1.18 | 0.36 | -3.25 | 1.16E-03 | 9.63E-03 |
| dd_Smed_v6_9267_0_1 | 294.41 | -1.18 | 0.33 | -3.60 | 3.12E-04 | 3.64E-03 |
| dd_Smed_v6_8984_0_1 | 265.99 | -1.17 | 0.33 | -3.52 | 4.27E-04 | 4.59E-03 |
| dd_Smed_v6_12214_0_1 | 311.47 | -1.17 | 0.34 | -3.42 | 6.16E-04 | 6.05E-03 |
| dd_Smed_v6_3941_0_1 | 637.93 | -1.17 | 0.25 | -4.68 | 2.91E-06 | 1.06E-04 |
| dd_Smed_v6_2567_0_1 | 138.04 | -1.17 | 0.31 | -3.77 | 1.61E-04 | 2.26E-03 |
| dd_Smed_v6_639_2_1 | 391.15 | -1.17 | 0.26 | -4.45 | 8.75E-06 | 2.45E-04 |
| dd_Smed_v6_832_0_1 | 3600.99 | -1.17 | 0.34 | -3.44 | 5.81E-04 | 5.79E-03 |
| dd_Smed_v6_7604_0_1 | 160.47 | -1.16 | 0.33 | -3.54 | 3.98E-04 | 4.38E-03 |
| dd_Smed_v6_8523_0_1 | 171.26 | -1.16 | 0.28 | -4.11 | 3.99E-05 | 7.67E-04 |
| dd_Smed_v6_6815_0_1 | 117.48 | -1.16 | 0.26 | -4.52 | 6.07E-06 | 1.85E-04 |
| dd_Smed_v6_5091_0_1 | 125.07 | -1.16 | 0.29 | -4.04 | 5.39E-05 | 9.67E-04 |
| dd_Smed_v6_6575_0_1 | 407.57 | -1.16 | 0.29 | -3.95 | 7.83E-05 | 1.29E-03 |
| dd_Smed_v6_5053_0_1 | 750.13 | -1.15 | 0.26 | -4.43 | 9.48E-06 | 2.58E-04 |
| dd_Smed_v6_3420_0_1 | 299.98 | -1.15 | 0.28 | -4.16 | 3.13E-05 | 6.35E-04 |
| dd_Smed_v6_7193_0_1 | 176.07 | -1.15 | 0.30 | -3.82 | 1.33E-04 | 1.98E-03 |
| dd_Smed_v6_8117_0_1 | 391.16 | -1.15 | 0.28 | -4.16 | 3.24E-05 | 6.52E-04 |

| | | | | | | |
|---|---|---|---|---|---|---|
| dd_Smed_v6_2010_0_1 | 141.34 | -1.15 | 0.23 | -5.02 | 5.04E-07 | 2.65E-05 |
| dd_Smed_v6_271_0_1 | 798.01 | -1.14 | 0.30 | -3.84 | 1.26E-04 | 1.90E-03 |
| dd_Smed_v6_5274_0_1 | 213.70 | -1.14 | 0.31 | -3.64 | 2.69E-04 | 3.26E-03 |
| dd_Smed_v6_7062_0_1 | 190.06 | -1.14 | 0.32 | -3.60 | 3.13E-04 | 3.64E-03 |
| dd_Smed_v6_2627_0_1 | 678.06 | -1.14 | 0.23 | -4.85 | 1.27E-06 | 5.39E-05 |
| dd_Smed_v6_9549_0_1 | 107.09 | -1.13 | 0.35 | -3.26 | 1.12E-03 | 9.37E-03 |
| dd_Smed_v6_7075_0_1 | 98.11 | -1.13 | 0.35 | -3.25 | 1.16E-03 | 9.63E-03 |
| dd_Smed_v6_5242_0_1 | 267.67 | -1.13 | 0.32 | -3.54 | 4.07E-04 | 4.44E-03 |
| dd_Smed_v6_6557_0_1 | 225.93 | -1.13 | 0.29 | -3.95 | 7.79E-05 | 1.28E-03 |
| dd_Smed_v6_7_0_1 | 638759.64 | -1.13 | 0.25 | -4.54 | 5.61E-06 | 1.74E-04 |
| dd_Smed_v6_14_0_1 | 680.05 | -1.12 | 0.35 | -3.24 | 1.18E-03 | 9.72E-03 |
| dd_Smed_v6_442_0_1 | 534.73 | -1.12 | 0.33 | -3.38 | 7.31E-04 | 6.84E-03 |
| dd_Smed_v6_9115_0_1 | 479.23 | -1.12 | 0.23 | -4.86 | 1.16E-06 | 5.04E-05 |
| dd_Smed_v6_5727_0_1 | 486.16 | -1.12 | 0.34 | -3.24 | 1.18E-03 | 9.71E-03 |
| dd_Smed_v6_2825_0_1 | 878.24 | -1.11 | 0.15 | -7.33 | 2.26E-13 | 1.14E-10 |
| dd_Smed_v6_11199_0_1 | 157.74 | -1.11 | 0.29 | -3.86 | 1.13E-04 | 1.75E-03 |
| dd_Smed_v6_8659_0_1 | 228.36 | -1.10 | 0.26 | -4.23 | 2.37E-05 | 5.14E-04 |
| dd_Smed_v6_4392_0_1 | 459.38 | -1.10 | 0.26 | -4.24 | 2.23E-05 | 4.92E-04 |
| dd_Smed_v6_3132_0_1 | 197.87 | -1.10 | 0.21 | -5.23 | 1.74E-07 | 1.15E-05 |
| dd_Smed_v6_1673_0_1 | 430.97 | -1.10 | 0.33 | -3.36 | 7.89E-04 | 7.20E-03 |
| dd_Smed_v6_11004_0_1 | 762.73 | -1.10 | 0.24 | -4.60 | 4.14E-06 | 1.38E-04 |
| dd_Smed_v6_1904_0_1 | 221.74 | -1.10 | 0.31 | -3.49 | 4.75E-04 | 5.00E-03 |
| dd_Smed_v6_4942_0_1 | 455.27 | -1.09 | 0.27 | -4.10 | 4.15E-05 | 7.90E-04 |
| dd_Smed_v6_2210_0_1 | 581.08 | -1.09 | 0.25 | -4.39 | 1.11E-05 | 2.90E-04 |
| dd_Smed_v6_4753_0_1 | 199.17 | -1.09 | 0.29 | -3.79 | 1.53E-04 | 2.19E-03 |
| dd_Smed_v6_5967_0_1 | 123.55 | -1.08 | 0.29 | -3.76 | 1.73E-04 | 2.40E-03 |
| dd_Smed_v6_1399_0_1 | 784.60 | -1.08 | 0.31 | -3.53 | 4.23E-04 | 4.56E-03 |
| dd_Smed_v6_7864_0_1 | 379.30 | -1.08 | 0.25 | -4.32 | 1.57E-05 | 3.86E-04 |
| dd_Smed_v6_7356_0_1 | 467.81 | -1.08 | 0.28 | -3.88 | 1.04E-04 | 1.63E-03 |
| dd_Smed_v6_1916_0_1 | 1500.75 | -1.07 | 0.19 | -5.72 | 1.09E-08 | 1.15E-06 |
| dd_Smed_v6_7590_0_1 | 117.13 | -1.07 | 0.28 | -3.80 | 1.48E-04 | 2.13E-03 |
| dd_Smed_v6_7940_0_1 | 147.10 | -1.07 | 0.33 | -3.26 | 1.10E-03 | 9.24E-03 |
| dd_Smed_v6_4349_0_1 | 1190.44 | -1.07 | 0.17 | -6.39 | 1.67E-10 | 2.99E-08 |
| dd_Smed_v6_6330_0_1 | 180.78 | -1.07 | 0.31 | -3.41 | 6.55E-04 | 6.29E-03 |
| dd_Smed_v6_6303_0_1 | 245.69 | -1.07 | 0.31 | -3.42 | 6.17E-04 | 6.06E-03 |
| dd_Smed_v6_8089_0_1 | 213.57 | -1.06 | 0.31 | -3.46 | 5.48E-04 | 5.54E-03 |
| dd_Smed_v6_4776_0_1 | 172.81 | -1.06 | 0.23 | -4.63 | 3.58E-06 | 1.25E-04 |
| dd_Smed_v6_1060_0_1 | 820.79 | -1.06 | 0.24 | -4.48 | 7.49E-06 | 2.19E-04 |
| dd_Smed_v6_6406_0_1 | 459.64 | -1.06 | 0.24 | -4.46 | 8.34E-06 | 2.35E-04 |
| dd_Smed_v6_12684_0_1 | 115.86 | -1.05 | 0.31 | -3.41 | 6.47E-04 | 6.25E-03 |
| dd_Smed_v6_7127_0_1 | 421.49 | -1.05 | 0.26 | -4.05 | 5.20E-05 | 9.38E-04 |
| dd_Smed_v6_4547_0_1 | 231.47 | -1.05 | 0.29 | -3.64 | 2.74E-04 | 3.30E-03 |
| dd_Smed_v6_6660_0_1 | 400.85 | -1.05 | 0.25 | -4.12 | 3.82E-05 | 7.44E-04 |

| | | | | | | |
|---|---|---|---|---|---|---|
| dd_Smed_v6_603_0_1 | 397.78 | -1.04 | 0.28 | -3.66 | 2.56E-04 | 3.14E-03 |
| dd_Smed_v6_8306_0_1 | 324.08 | -1.04 | 0.27 | -3.82 | 1.32E-04 | 1.97E-03 |
| dd_Smed_v6_10889_0_1 | 466.99 | -1.04 | 0.26 | -4.06 | 4.89E-05 | 8.95E-04 |
| dd_Smed_v6_5327_0_1 | 532.71 | -1.04 | 0.20 | -5.14 | 2.75E-07 | 1.65E-05 |
| dd_Smed_v6_9926_1_1 | 59.15 | -1.03 | 0.29 | -3.54 | 4.03E-04 | 4.41E-03 |
| dd_Smed_v6_8959_0_1 | 301.63 | -1.03 | 0.22 | -4.61 | 4.11E-06 | 1.38E-04 |
| dd_Smed_v6_125_0_1 | 7706.95 | -1.03 | 0.26 | -3.97 | 7.05E-05 | 1.19E-03 |
| dd_Smed_v6_4687_0_1 | 182.55 | -1.03 | 0.27 | -3.77 | 1.65E-04 | 2.30E-03 |
| dd_Smed_v6_4642_0_1 | 178.18 | -1.03 | 0.32 | -3.25 | 1.16E-03 | 9.65E-03 |
| dd_Smed_v6_9460_0_1 | 603.70 | -1.03 | 0.31 | -3.30 | 9.70E-04 | 8.42E-03 |
| dd_Smed_v6_10066_0_1 | 292.50 | -1.03 | 0.31 | -3.32 | 8.85E-04 | 7.88E-03 |
| dd_Smed_v6_2828_0_1 | 567.55 | -1.02 | 0.20 | -5.20 | 2.01E-07 | 1.29E-05 |
| dd_Smed_v6_329_0_1 | 1541.46 | -1.02 | 0.17 | -5.87 | 4.35E-09 | 5.26E-07 |
| dd_Smed_v6_493_0_1 | 3205.05 | -1.02 | 0.32 | -3.25 | 1.17E-03 | 9.70E-03 |
| dd_Smed_v6_2272_0_1 | 762.55 | -1.02 | 0.24 | -4.35 | 1.39E-05 | 3.47E-04 |
| dd_Smed_v6_1518_0_1 | 965.97 | -1.02 | 0.24 | -4.23 | 2.33E-05 | 5.09E-04 |
| dd_Smed_v6_6320_0_1 | 160.75 | -1.02 | 0.27 | -3.80 | 1.43E-04 | 2.09E-03 |
| dd_Smed_v6_13864_0_1 | 103.35 | -1.02 | 0.24 | -4.28 | 1.84E-05 | 4.31E-04 |
| dd_Smed_v6_12866_0_1 | 335.94 | -1.01 | 0.26 | -3.87 | 1.08E-04 | 1.68E-03 |
| dd_Smed_v6_7215_0_1 | 471.50 | -1.01 | 0.25 | -4.07 | 4.67E-05 | 8.65E-04 |
| dd_Smed_v6_4573_0_1 | 616.38 | -1.01 | 0.22 | -4.53 | 6.04E-06 | 1.85E-04 |
| dd_Smed_v6_1893_0_1 | 825.80 | -1.01 | 0.27 | -3.69 | 2.28E-04 | 2.90E-03 |
| dd_Smed_v6_5407_0_1 | 1125.79 | -1.01 | 0.18 | -5.49 | 4.05E-08 | 3.62E-06 |
| dd_Smed_v6_8606_0_1 | 277.59 | -1.01 | 0.24 | -4.13 | 3.64E-05 | 7.14E-04 |
| dd_Smed_v6_13753_0_1 | 275.32 | -1.01 | 0.28 | -3.58 | 3.41E-04 | 3.91E-03 |
| dd_Smed_v6_6231_0_1 | 300.29 | -1.01 | 0.30 | -3.36 | 7.91E-04 | 7.21E-03 |
| dd_Smed_v6_5600_0_1 | 1943.75 | -1.00 | 0.24 | -4.14 | 3.42E-05 | 6.81E-04 |
| dd_Smed_v6_5330_0_1 | 314.05 | -1.00 | 0.24 | -4.15 | 3.32E-05 | 6.64E-04 |
| dd_Smed_v6_11160_0_1 | 381.51 | -1.00 | 0.22 | -4.60 | 4.24E-06 | 1.41E-04 |

**Table S3: List of genes assayed in this paper and their gene expression profile in RNA-seq analysis**
ID: Planmine transcript ID from dd_Smed_v6 transcriptome.
baseMean: mean of normalized counts for all samples;
log2FC: log2 fold change;
lfcSE: standard error of log2 fold change;
stat: Wald statistic;
pvalue: Wald test p-value;
padj: BH adjusted p-values
DE genes: Yes=present in Table S1 or S2; No=DDR pathway component mentioned in Fig. S1B

| ID | Gene | baseMean | log2FC | lfcSE | stat | pvalue | padj | DE genes |
|---|---|---|---|---|---|---|---|---|
| dd_Smed_v6_14586_0_1 | atm | 1184.11 | 0.45 | 0.23 | 1.99 | 4.65E-02 | 1.25E-01 | No |
| dd_Smed_v6_13527_0_1 | rad50 | 103.32 | -1.06 | 0.45 | -2.33 | 1.97E-02 | 6.92E-02 | No |
| dd_Smed_v6_6620_0_1 | chk2 | 2493.37 | 0.27 | 0.18 | 1.49 | 1.35E-01 | 2.61E-01 | No |
| dd_Smed_v6_8198_0_1 | ercc2 | 167.87 | 0.12 | 0.37 | 0.31 | 7.53E-01 | 8.40E-01 | No |
| dd_Smed_v6_8614_0_1 | cdk7 | 585.35 | 1.32 | 0.18 | 7.27 | 3.48E-13 | 1.58E-10 | Yes |
| dd_Smed_v6_11510_0_1 | TFIIH4 | 564.25 | 0.45 | 0.16 | 2.80 | 5.11E-03 | 2.72E-02 | No |
| dd_Smed_v6_7018_0_1 | TFIIH2 | 583.77 | 1.09 | 0.24 | 4.48 | 7.49E-06 | 2.19E-04 | Yes |
| dd_Smed_v6_12789_0_1 | chk1 | 1363.17 | 1.51 | 0.25 | 6.02 | 1.74E-09 | 2.40E-07 | Yes |
| dd_Smed_v6_10725_0_1 | cyclinH | 538.85 | 1.37 | 0.19 | 7.17 | 7.36E-13 | 3.00E-10 | Yes |
| dd_Smed_v6_7837_0_1 | cyclinB2 | 4274.086 | 0.91 | 0.26 | 3.49 | 4.81E-04 | 5.04E-03 | No |
| dd_Smed_v6_20423_0_1 | rad17 | 281.16 | 1.40 | 0.24 | 5.84 | 5.21E-09 | 6.16E-07 | Yes |
| dd_Smed_v6_5563_0_1 | p53 | 2655.18 | 0.13 | 0.18 | 0.74 | 4.60E-01 | 6.06E-01 | No |
| dd_Smed_v6_4961_0_1 | mre11 | 2244.58 | 0.69 | 0.20 | 3.48 | 4.95E-04 | 5.16E-03 | No |
| dd_Smed_v6_13269_0_1 | nbs1 | 1349.49 | 0.71 | 0.32 | 2.25 | 2.43E-02 | 8.02E-02 | No |
| dd_Smed_v6_8754_0_1 | atr | 2220.78 | 0.41 | 0.17 | 2.38 | 1.71E-02 | 6.31E-02 | No |
| dd_Smed_v6_11363_0_1 | lig4 | 222.43 | 0.10 | 0.30 | 0.33 | 7.42E-01 | 8.32E-01 | No |
| dd_Smed_v6_8421_0_1 | ku80 | 2560.21 | 0.33 | 0.18 | 1.84 | 6.63E-02 | 1.60E-01 | No |
| dd_Smed_v6_9725_0_1 | ku70 | 123.75 | -0.13 | 0.27 | -0.49 | 6.24E-01 | 7.43E-01 | No |
| dd_Smed_v6_12096_0_1 | artemis | 133.88 | 1.06 | 0.34 | 3.07 | 2.12E-03 | 1.46E-02 | Yes |
| dd_Smed_v6_8462_0_1 | dna-pk | 124.34 | -0.27 | 0.38 | -0.71 | 4.80E-01 | 6.24E-01 | No |
| dd_Smed_v6_8626_0_1 | rad51 | 1858.18 | 0.35 | 0.22 | 1.59 | 1.13E-01 | 2.29E-01 | No |
| dd_Smed_v6_12551_0_1 | brca2 | 835.75 | 0.76 | 0.21 | 3.58 | 3.49E-04 | 3.96E-03 | No |
| dd_Smed_v6_16638_0_1 | FANC-J | 88.81 | 1.20 | 0.45 | 2.65 | 8.05E-03 | 3.74E-02 | Yes |
| dd_Smed_v6_10338_0_1 | PARP1 | 2523.82 | 0.43 | 0.22 | 1.91 | 5.61E-02 | 1.42E-01 | No |
| dd_Smed_v6_6154_0_1 | PARP2 | 0.61 | -3.86 | 4.25 | -0.91 | 3.64E-01 | NA | No |
| dd_Smed_v6_2611_0_1 | PARP3 | 439.08 | -1.41 | 0.47 | -2.99 | 2.75E-03 | 1.73E-02 | Yes |
| dd_Smed_v6_14158_0_1 | RPA | 220.65 | 1.28 | 0.35 | 3.64 | 2.73E-04 | 3.29E-03 | Yes |

**Table S4. Primers and transcript IDs for genes used in this study.**

| Gene | Smed ID | Forward primer | Reverse primer |
|---|---|---|---|
| *unc-22* | ---- | GACGTAAACGGCCACAAGTT | CTTGTACAGCTCGTCCATGC |
| *atm* | dd_Smed_v6_14586_0_1 | AGAATGCCTGCAAAATGAAGA | ACCACACTATATGAAGGGAACAAA |
| *atr* | dd_Smed_v6_8754_0_1 | ATTCCTGGCCCAATAAAAGAA | GAGCAAACCGAAAATGAGCTA |
| *dna-pk* | dd_Smed_v6_8462_0_1 | GCAAAACGGGGTAGTCGTAA | CGAATTCGGATTTGTTTCGT |
| *mre11* | dd_Smed_v6_4961_0_1 | TTGATGAAACTTCGTAAAGCACT | TTTCTTTTTCCCTTTGCCATC |
| *nbs1* | dd_Smed_v6_13269_0_1 | TGGCGATATCGTTACTTTTGG | GGCACGGAATTACGTTCAATA |
| *rad50* | dd_Smed_v6_11475_0_2 | CCGAGCGAAATGAATTACAGA | TTTTTGAATTTGGAGGCTTGA |
| *chk2* | dd_Smed_v6_6620_0_1 | CAGATAGTAGTTACACAGTTGGAAGA | GCTCGCCAGACAAGGAATTA |
| *ercc2* | dd_Smed_v6_8198_0_1 | GACGCCGACAATTCCATTTC | CTATATCTGACGGGCTCACATC |
| *cdk7* | dd_Smed_v6_8614_0_1 | AGGTGAAGGCCAATTTGCTA | CAAGGTCAGTTATCATTGAATTGGG |
| *tfIIh4* | dd_Smed_v6_11510_0_1 | TGGACCTCAAGGTGAATGTATG | GGCGAAAGGGAAACGTTTATG |
| *p53* | dd_Smed_v6_5563_0_1 | CTGCTTTTAAATCCGACGACA | TGCACATAGCACACATGACAA |
| *tfIIh2* | dd_Smed_v6_7018_0_1 | CTTCGAACGTATGGATAGGGAAG | CACATTCCGAGACCCGAATC |
| *chk1* | dd_Smed_v6_12789_0_1 | CAGCACCTCAAGTCTGAAATTG | CGTCGGAAGTCCACCAAA |
| *cyclinB2* | dd_Smed_v6_7837_0_1 | GCCATTGGTGCTCCGTAAT | TCCCATTATTACCAAACGGATGAG |
| *rad17* | dd_Smed_v6_20423_0_1 | TTCAGACCTAGCTGTTCATAGTAAA | ATGAAACGAAAGCTGAGAGAAATC |
| *cyclinH* | dd_Smed_v6_10725_0_1 | GCAATCTGCCCGATGGAT | TGATTGATAGACTTCATGTTGAGGA |
| *lig4* | dd_Smed_v6_11363_0_1 | CGCGAAGACAAATCTCCATT | GCTCATTACAGCGCATACGA |
| *ku80* | dd_Smed_v6_8421_0_1 | AGCTACCAGGGGAAATTGCT | CATGTTGGTACGGGCTTCTT |
| *ku70* | dd_Smed_v6_9725_0_1 | TTTCTAACTGCACGTTTTTCAA | CAAAATTACGCAAACTTTATTCACA |
| *artemis* | dd_Smed_v6_12096_0_1 | TGAATATCCACAAATTGCCATA | TGGTGTGAAAAACACACTCG |
| *atm_Nterm* | dd_Smed_v6_14856_0_1 | TGAAATTGCTTCTGCCACAG | TTGATCCCAGGAGATTTTGC |
| *atm_Cterm* | dd_Smed_v6_14856_0_1 | TGGTCGAACAATTGGTGAAA | ATGGATCGTGAAGCAAAACC |
